# Progressive decline in the levels of six miRNAs from parents to children in autism

**DOI:** 10.1101/2022.10.20.512859

**Authors:** Minoo Rassoulzadegan, Ecmel Mehmetbeyoglu, Zeynep Yilmaz, Serpil Taheri, Yusuf Ozkul

## Abstract

The growing burden of a gradual increase in births of children with autism has placed it at the center of the concerns of major laboratories. We have previously detected a decrease in the levels of six miRNAs (miR-19a-3p, miR-361-5p, miR-3613-3p, miR-150-5p, miR-126-3p, and miR-499a-5p) in parents and their children inherited at a lower level. Here, we suggest that down-regulation of each of these six miRNAs inherited from parents contributes to the development of children with autism. We compare their levels of distribution in each family between the autistic child and siblings. We find that the distribution of levels of these miRNAs in siblings (undiagnosed as autism) is not always higher than in autistic children, but it is at varying levels. These data support a model in which autistic behavior relies on low levels of the six miRNAs expressed in children potentially associated with autistic syndrome (ASD). The intimate link between miRNAs levels and behavioral characteristics suggests possibilities for understanding the basic circuitry involved in autism and thus advancing partial knowledge of brain functions. An early diagnosis of autism helps provide children an environment conducive to their development.

**Graphical abstract:** 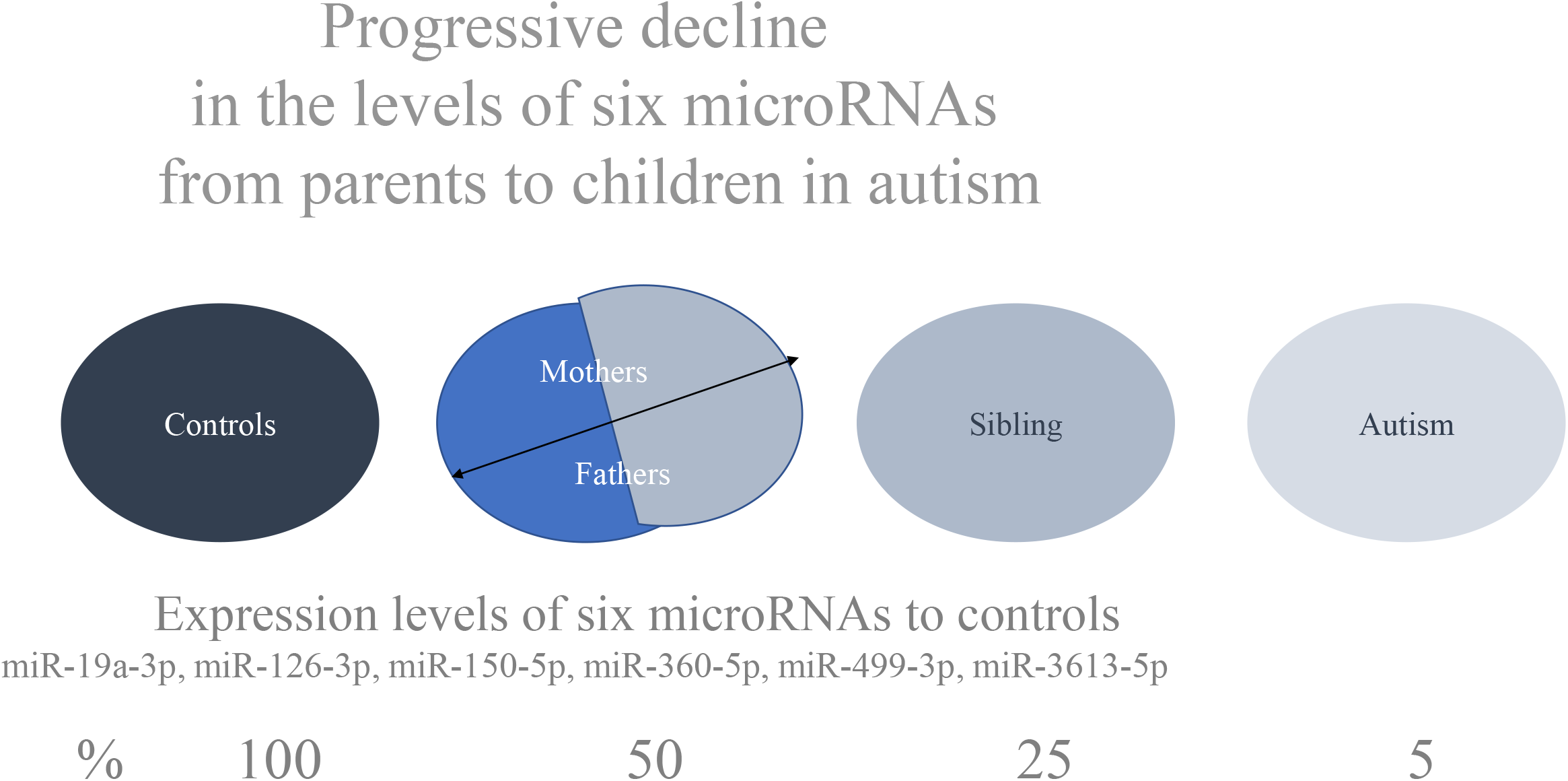

## Introduction

Understanding the Autism Spectrum Disorder (ASD) - impairment in communication and social interaction with stereotypic and repetitive patterns of behavior (reviewed in refs^1,2,3^) - appears more and more as a societal^4^ as well as a scientific challenge. At an increasing frequency, it affects the most complex functions of the human brain, raising critical questions for the neurobiologist, the social scientist and the geneticist. Its genetic origin is clearly indicated by the high concordance rates between monozygotic twins^5,6,7^, but given the complex variations of the phenotype, even in twins indicate a strong involvement of epigenetic mechanisms. In the long term, their understanding will have to be part of the molecular analysis of brain functions, the most advanced levels of biological complexity. In a more immediate perspective, establishing a clear etiology of ASD could lead to useful medical developments for expert medical as well as parental advice and, most important for the physician in charge, to assist in the early identification of affected children, so as to provide them with adequate conditions for their optimal development.

Large to very large cohorts of patients are currently under study in scientific and medical institutes throughout the world, a representative recent study involves no less than 35,584 subjects including 11,986 ASD patients, their families and controls^8,9^. Several lines of genetic research are currently in progress with the most advanced techniques of genome analysis. Protein coding alterations were detected in more than 100 mutated loci in the genomes of autistic patients^28,10^. In addition, genome-wide detection of tandem DNA repeats revealed their expansion in autism^9^ with many of them expressed at early developmental stages in neurons and neuronal precursors. Their identification may hopefully allow us to understand the mechanisms, extent and variability as well as the relationship of the disease with other neurodevelopmental defects. However, none of these identified genetic variants were detected in all and not even in a majority of patients and thus, would qualify as a key determinant of the disease. A distinct approach of the problem makes use of animal models, a hopeful counterpart of clinical studies^11,12^. The available mouse models reproduce characteristic features of the human disease and allow a variety of experimental approaches, which are not even to be considered in the case of human patients for ethical and practical reasons. Similarities between the clinical observations of patients and the psychological and cognitive alterations in the mouse models of ASD support the validity of this approach^12^.

The recent report by the physicians and scientists of Erciyes University^13^ is relevant to these various aspects of current ASD research. A first step was the assembly of a cohort of 45 patients from 37 independent families with for each family genetically related individuals (parents, brothers, sisters) and 37 unrelated controls, for a total of 187 serum samples. The analysis was then centered on the family of genes encoding the 22nt-long non-coding regulatory RNAs designated miRNAs for microRNAs and identified as key actors in cellular differentiation processes^14^. Their main activity at least that so far identified is to decrease expression of sequence-homologous target genes by inducing degradation of the mRNAs and blocking their translation. Several hundreds of them have been identified in higher organisms, each one controlling a variable number of mRNAs in a tissue in a target specific manner. They are involved in multiple processes including brain functions and alterations have been implicated in neuropathological conditions such as ASD^15,16^. In the assembled Kayseri cohort, miRNA PCR Array profiles of the expression of 372 miRNAs (see Ozkul et al.^13^ for expression and list of the microRNAs) led to the striking observation of a reduced expression of six of them among 372 miRNAs compared to healthy controls. The same modified profile was seen in every patient of the cohort. The six miRNAs (miR-19a-3p, miR-361-5p, miR-3613-3p, miR-150-5p, miR-126-3p, and miR-499a-5p) were all down-regulated in the serum of all 45 ASD patients of the Kayseri cohort. The penetrance of these correlations with all six miRNAs validate the significance of these low expression levels as an intrinsic feature of this disease (see raw data in Ozkul et al.^13^). Moreover, an independent observation of interest raises the key question of a possible hereditary transmission of the disease, or at least of a disease-prone state. Although clinically healthy, all first-degree family members of the 45 independent patients (brothers, sisters, fathers and mothers) showed a common pattern of progressive down-regulation. While the six miRNAs were all detected in higher amounts in unrelated individuals, a distinct pattern was registered among the clinically healthy first-degree family members of the patients with serum levels of the six miRNAs, in the range of 40 to 50 per cent of the unrelated individuals. It is likely that, although clinically normal, such people would in turn generate affected progenies, an assumption fully verified previously in animal models. The intermediate value of the “six miRNAs” therefore corresponds to a heritable high-risk state of healthy individuals.

The validity of conclusions extrapolated to autistic patients in general from the analysis of the Turkish cohort was checked by extending them to two of the established mouse models of the disease^12^: male animals treated with valproic acid (VPA) and *Cc2d1a* heterozygotes line. Five (miR-19a-3p, miR-361-5p, miR-150-5p, miR-126-3p, and miR-499a-5p) of these miRNAs are present in the mouse genome and were also down-regulated in sera, hippocampus and sperm of mouse models of autism like (VPA and *Cc2d1a+/-* see reference^13^).

VPA is known to induce a heritable pathological state with characteristic traits of the ASD pathology^17^ and indeed, mice of two inbred strains, *B6D2* and *Balb/c*, that had received one intraperitoneal injection of VPA according to the published protocols showed both the characteristic alterations in behavior and the low levels of expression of the miRNAs characteristic of the human patients. The same was true for a distinct established animal model of ASD, created in the mouse by a mutation of the *Cc2d1a* locus^11,13^, (one of the candidate genes identified in a few patients with autism) a repressor involved in the serotonin pathways.

The aims of the present review are to highlight miRNAs as markers of transgenerational behavioral changes revealing significant decreases (familiar profile) in the levels of six independent miRNAs along with their distribution from parents to autistic children and especially of their siblings (not diagnosed as autistic). Here, we examined the familial cases of six miRNAs distribution, compared to autistic children. From parents or at least one parent low levels are transmitted mainly to children with autism. In healthy siblings, the levels of the six miRNAs are seen at variable levels. This strongly suggests the decrease of all six miRNAs is required in children to express autism.

## Results and discussion

A first obvious comment concerns the novelty of the data reported by Ozkul and his colleagues in the search for a common genetic determinant associated with ASD, an area in which much effort has been invested without, so far, reaching a general conclusion^18,19,20^. The six miRNAs out of 372 microRNAs tested identified by Ozkul et al.^13^ were found expressed at low levels in 45 of the patients of the Kayseri cohort (Ozkul et al.^13^) and were summarized in Supplementary Table 1a. The six down-regulated miRNAs in the serum of 45 patients with ASD are miR-19a-3p, miR-361-5p, miR-3613-3p, miR-150-5p, miR-126-3p, and miR-499a-5p. Here we focus on the same six down-regulated miRNAs in each family, unlike those previously analyzed in Ozkul & al.^13^ globally on average. The six miRNAs have a unique sequence, target many transcripts and have certain targets in common see Table 1 and Figure 1 and Supplementary Table 2. None of the target is common to the six miRNAs. A short number of the target shared by five or four of them is listed in Table 2. Among the targets shared by four or five are many genes already identified in patients with autism.

**Figure 1.**
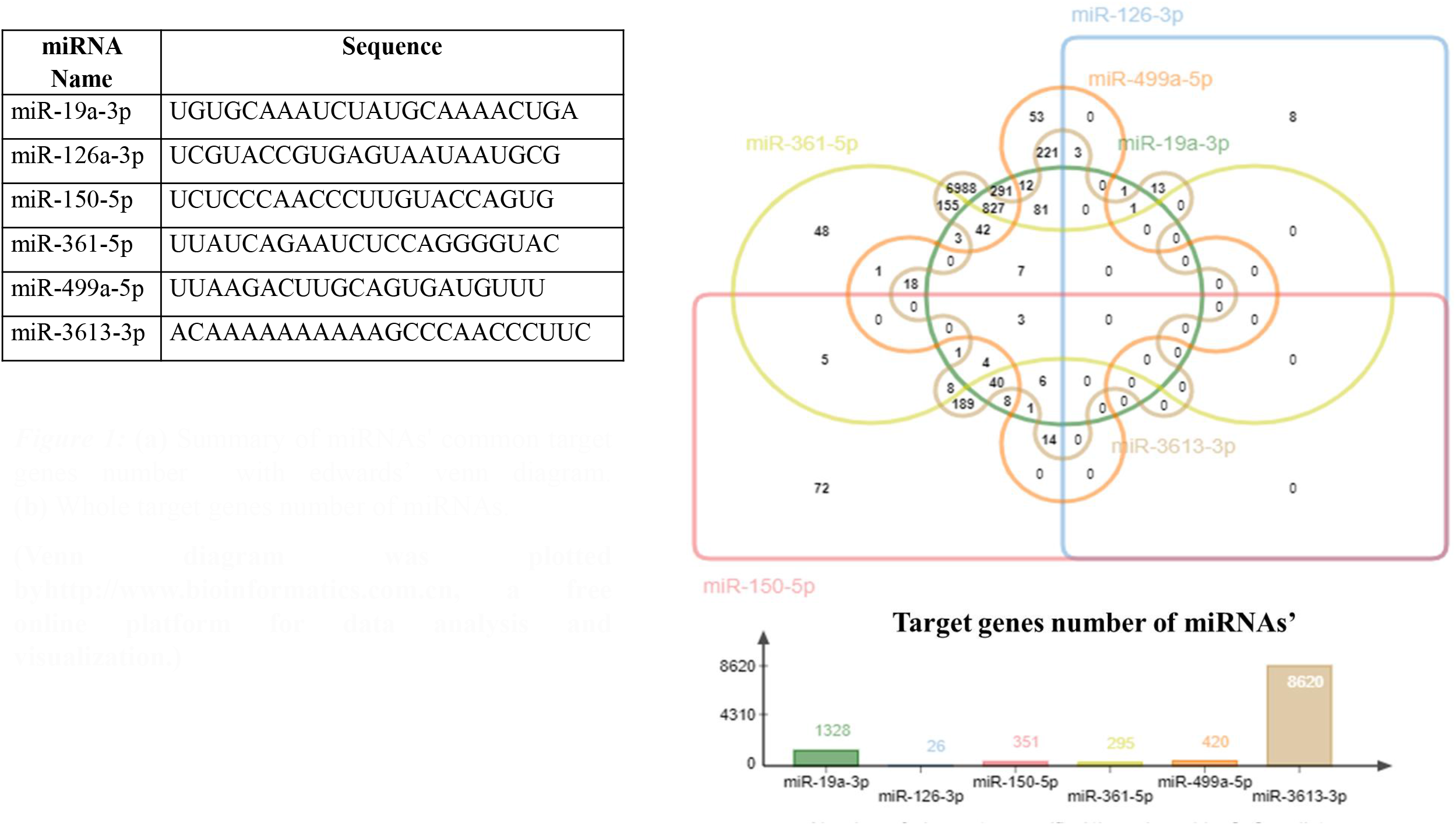
(a) Summary of miRNAs’ common target genes number with edwards’ venn diagram. **(b)** Whole target genes number of miRNAs. **(Venn diagram was plotted byhttp://www.bioinformatics.com.cn, a free online platform for data analysis and visualization.)**

### The hereditary decrease in six miRNAs

At the Erciyes University School of Medicine Hospital, Kayseri, Turkey a cohort (all samples n=189) of 45 patients with autism, their parents (n=74), unaffected siblings (n=33) and healthy members control age-matched children (n=21) and 16 control parents. The serum 372 miRNA expression profiles were compared with significant results (p<0.05).

In Supplementary Table 1a these results of miRNAs expression level data extracted from Ozkul et al.^13^ are presented as the mean of controls, which is arbitrarily considered to be 100 % and the % of changes are indicated for patients, sibling and parents of children. The hereditary decrease expressed as a percentage of controls shows in Supplementary Table 1a, a mean level of 50% for both parents. In Supplementary Table 1b-f shows example for family member that shows the highest value of one of the six miRNAs, in patients (girl or boy), or father, or mother and or sibling compared to 37 controls (healthy child and their parents). Individual family analysis suggests that both parents or at least one parent have low levels of the same miRNAs. On the part of either the mother or father, the down-regulation profile is mainly passed with even lower levels on to the autistic child. Figure 2 show five examples for each parent. It is important to compare the value of a given patient and their parents to all controls to visualize the inheritance of parental down-regulation.

**Figure 2.**
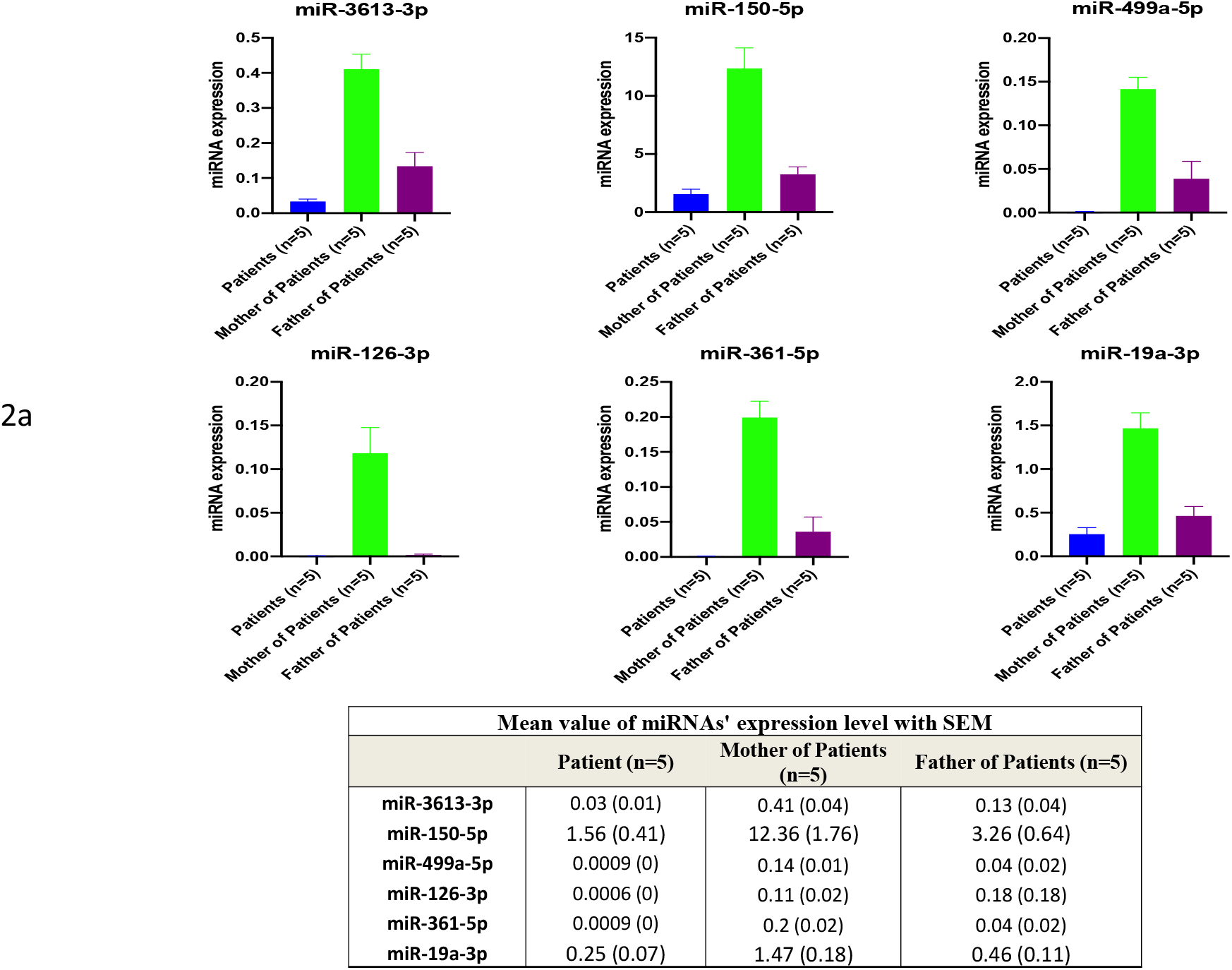
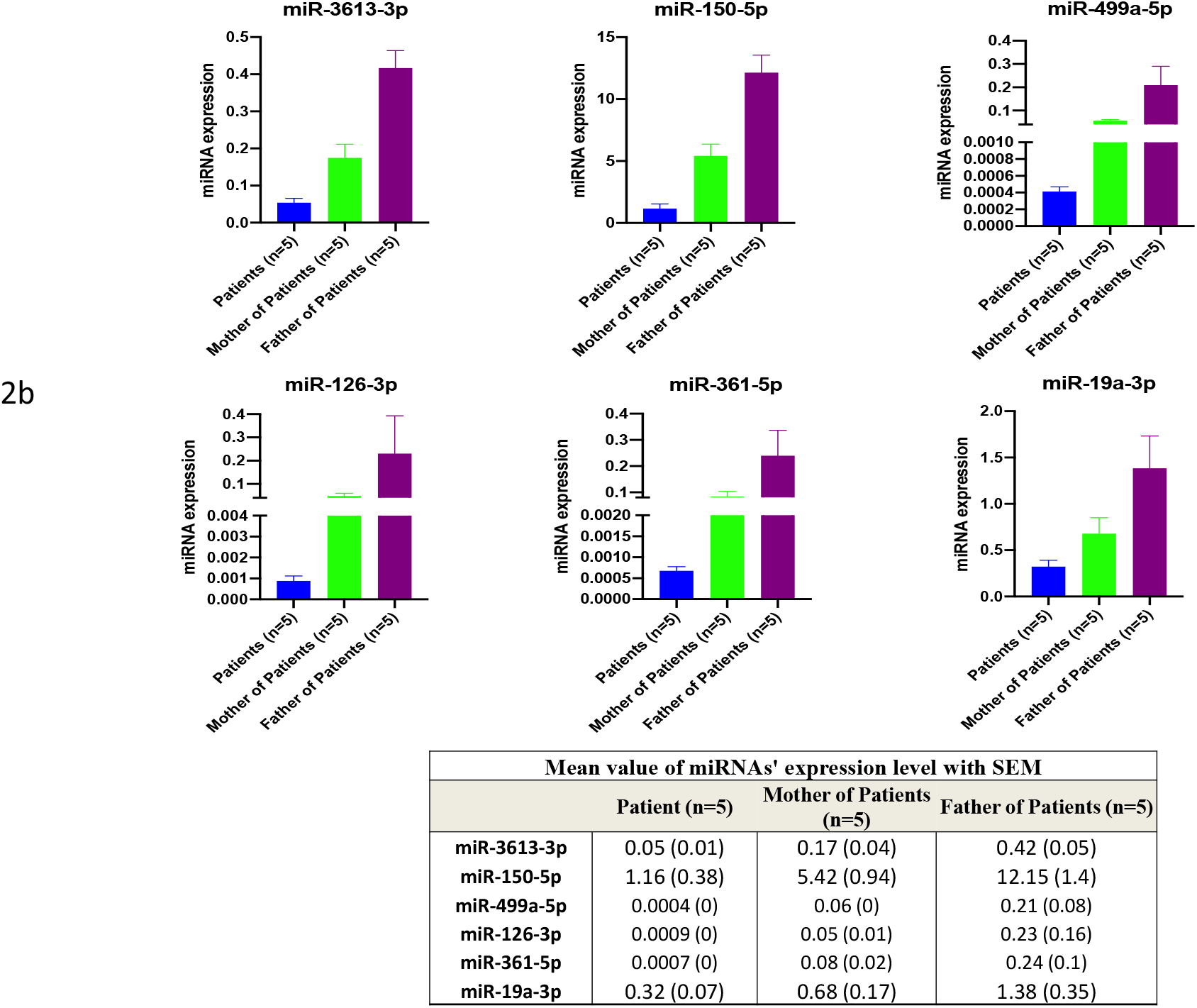
Individual family profiles for the highest values of “Six-miRNAs” (miR-19a-3p, miR-361-5p, miR-3613-3p, miR-150-5p, miR-126-3p, and miR-499a-5p) with respect to the control (mean value). The individual data are taken from the value presented in Table 1 of patients (girls and boys), mothers, fathers, sibling and controls and all raw values are presented in Supplementary tables 1 for each family. Data represents transcription values for “Six-miRNAs” (raw data after normalization)

### Distribution of six miRNAs in the sibling of autistic children

Despite the fact that all six miRNAs are at lower levels in siblings of children with autism compared to controls, their individual analysis in a given family is informative for the profiles of the inherited levels of each of the six miRNAs. Sibling escapes autism, so the question is, could the levels of ‘six miRNAs’ make a difference? Examples of individual family autistic children to siblings are shown in Figure 3a-e with the levels of each one of the six miRNAs in a single family separately compared between the child with autism and his breast siblings (not diagnosed as autistic). RT-Q-PCR tests validated the results of the global cohort analysis.

**Figure 3.**
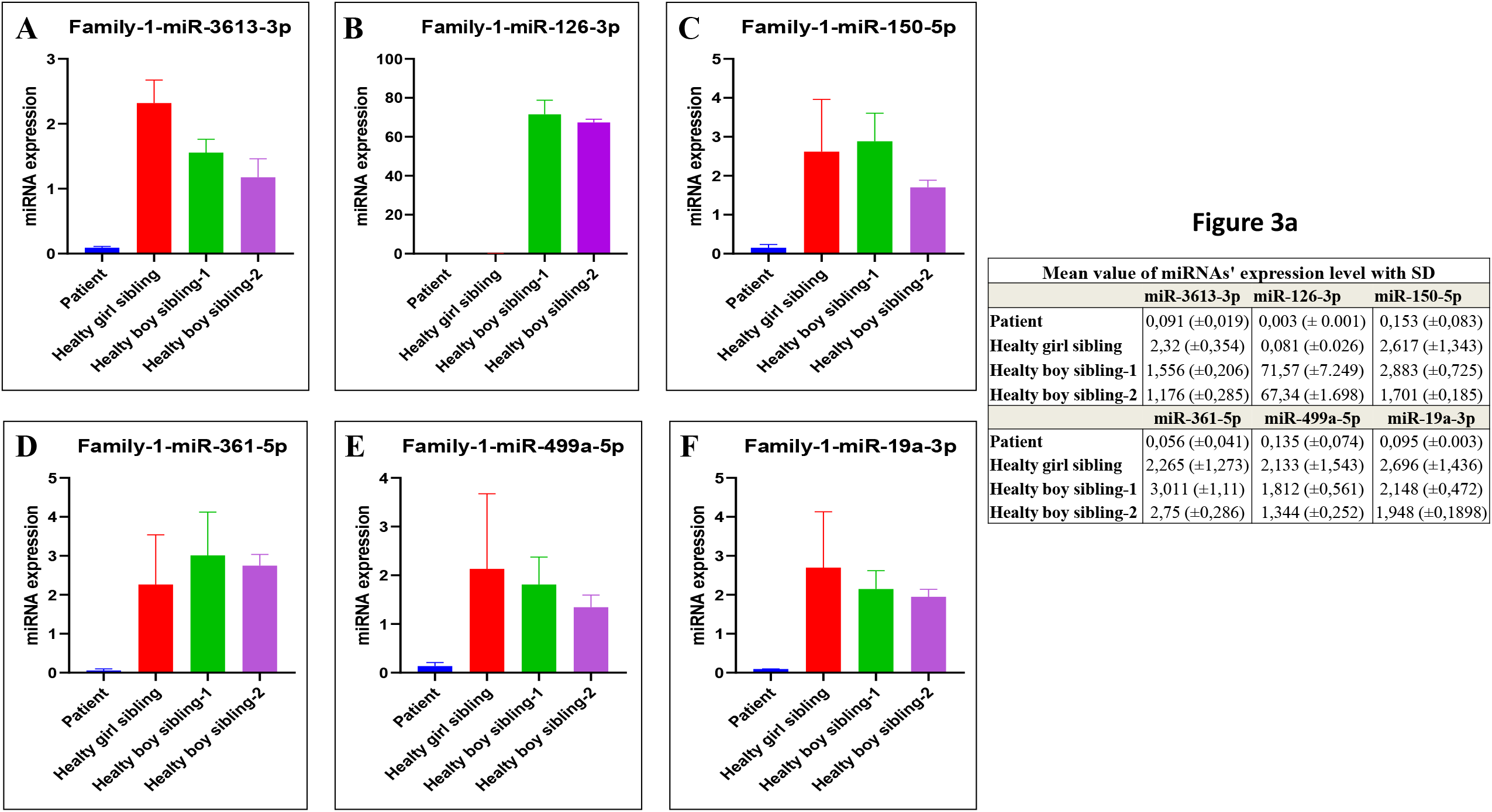
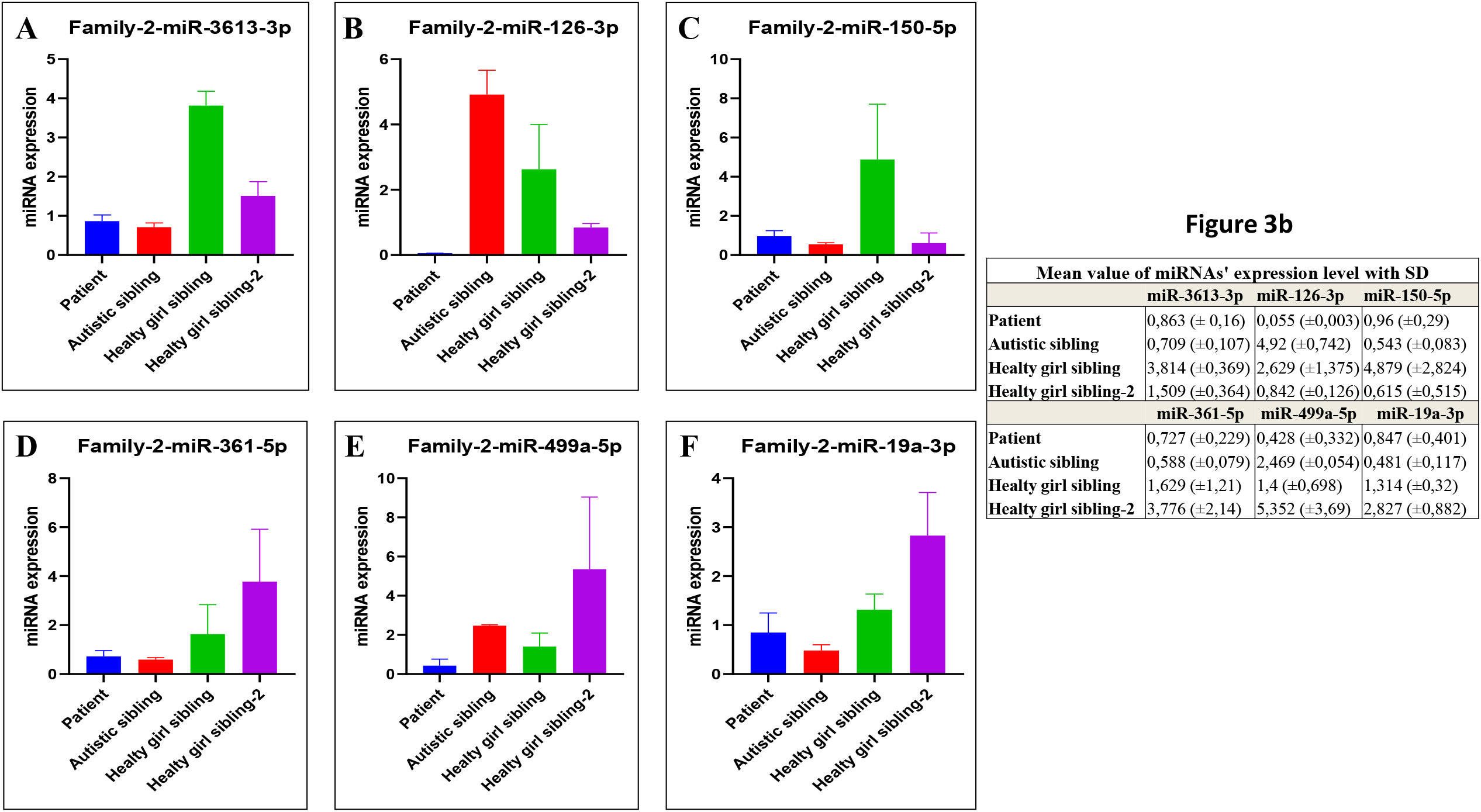
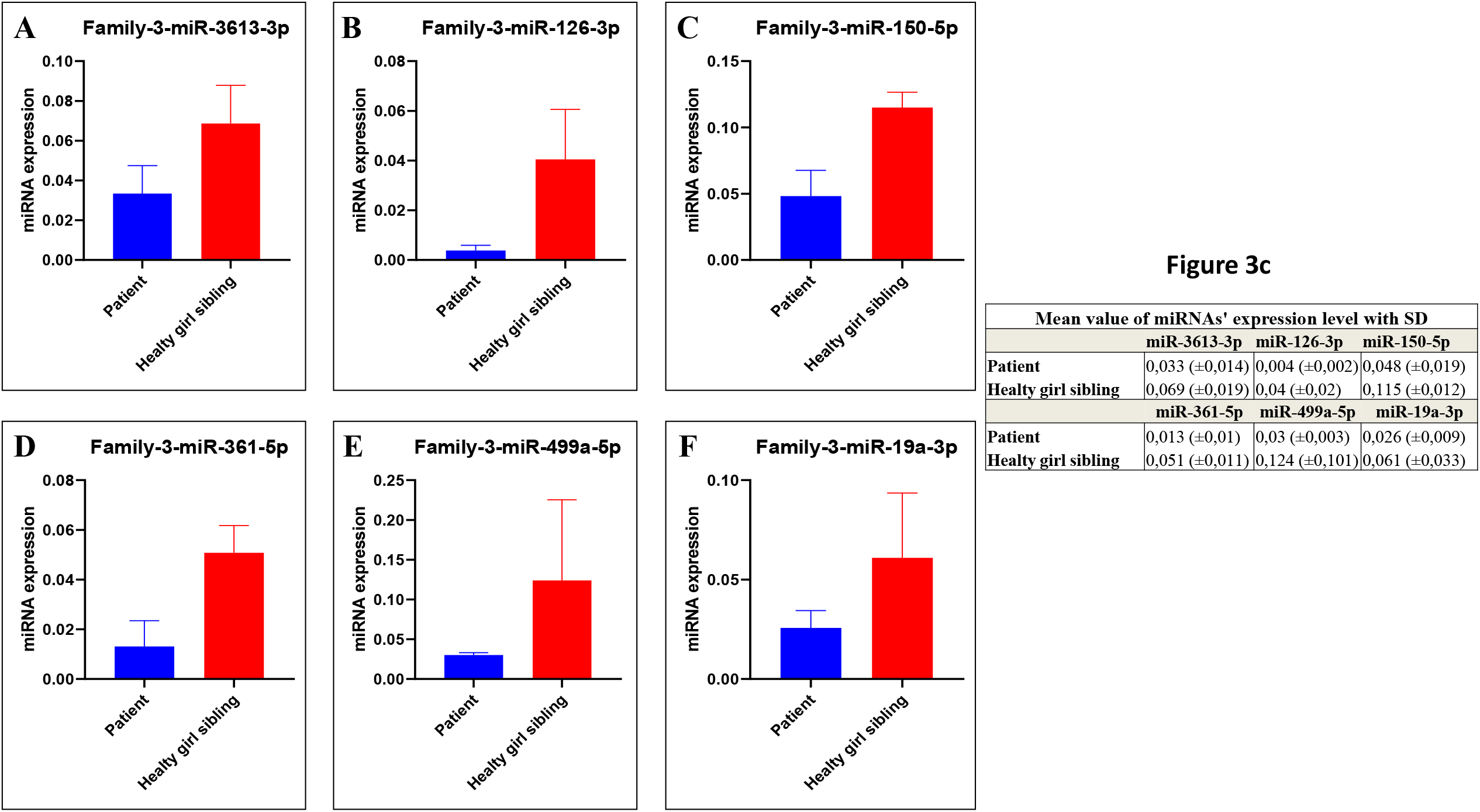
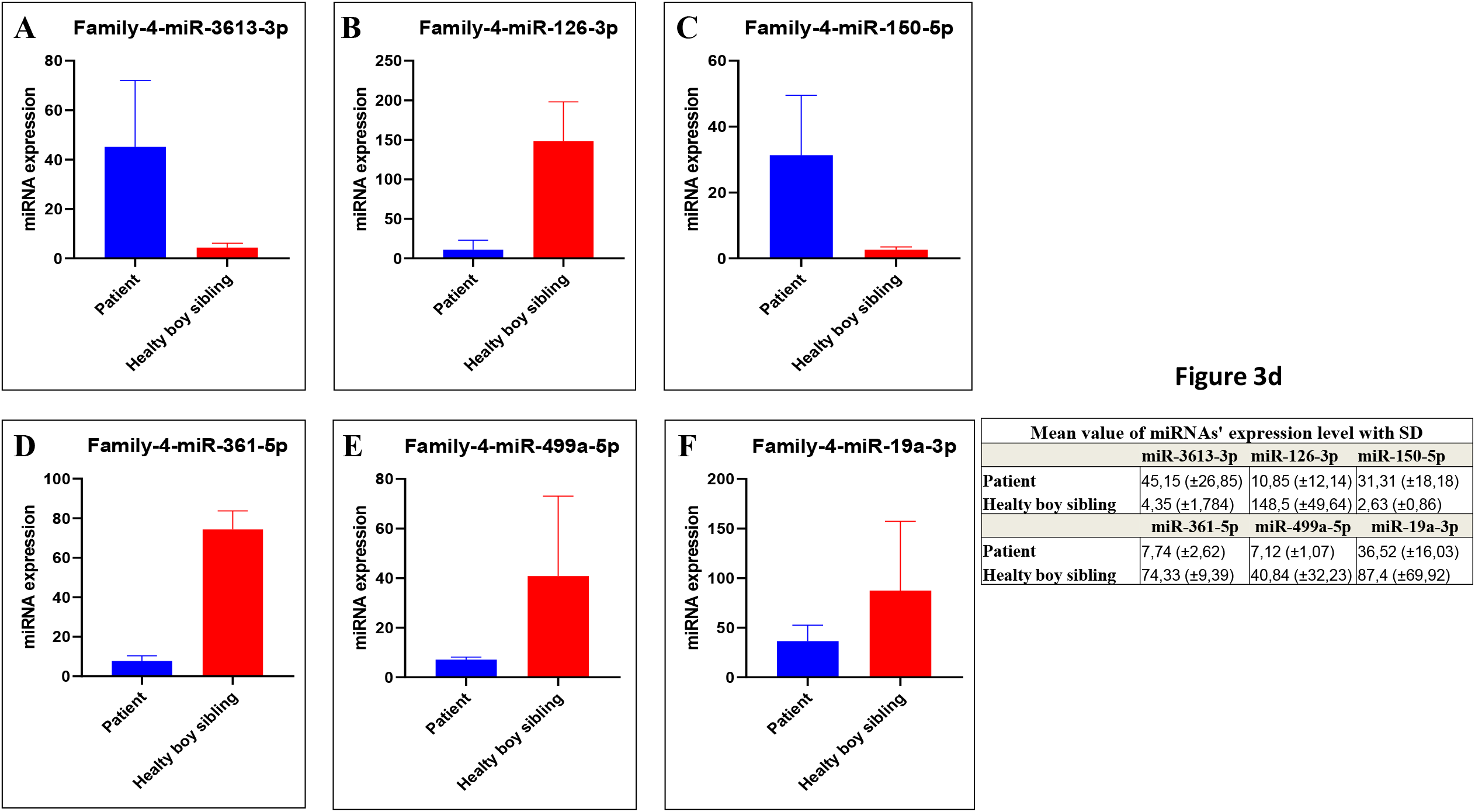
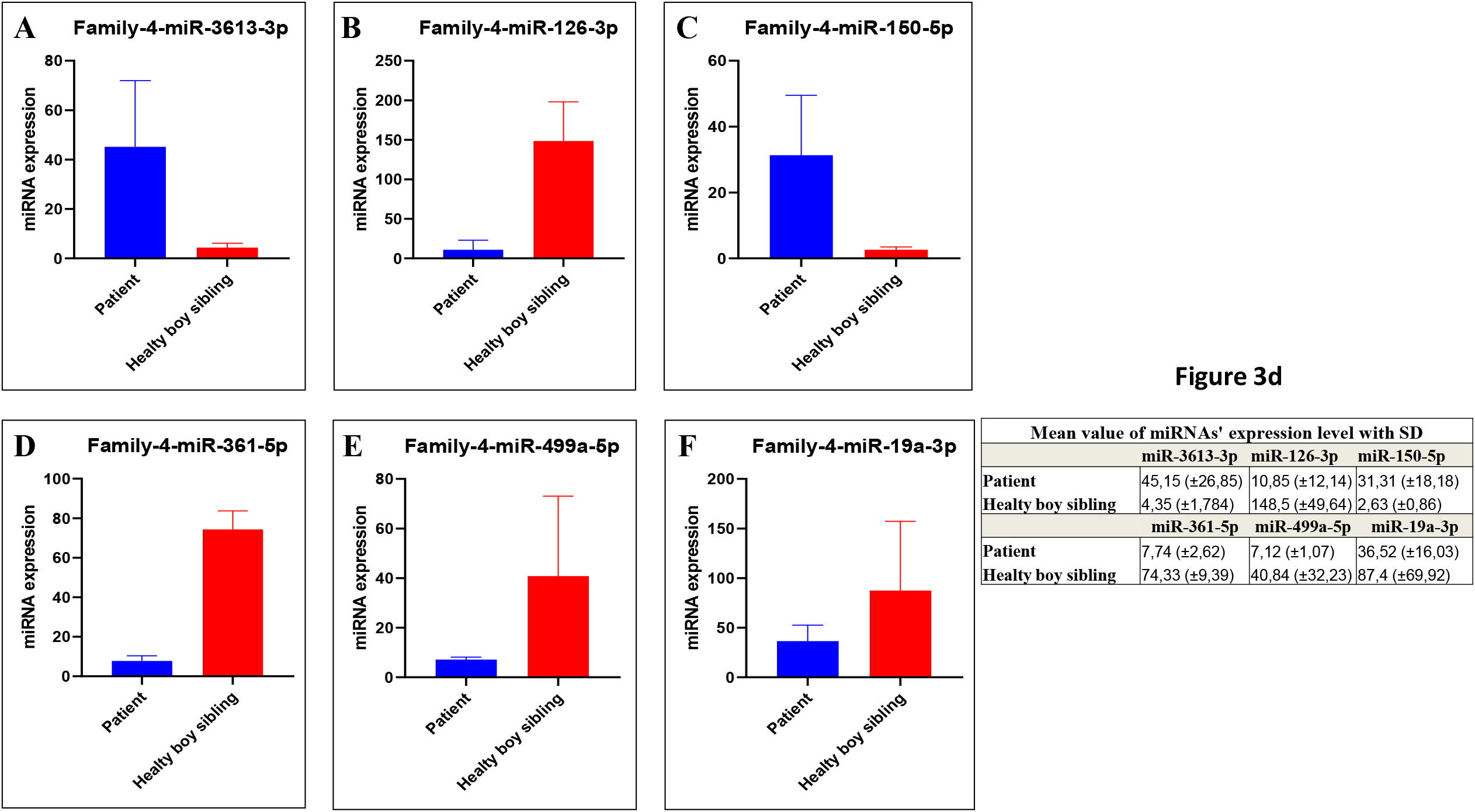
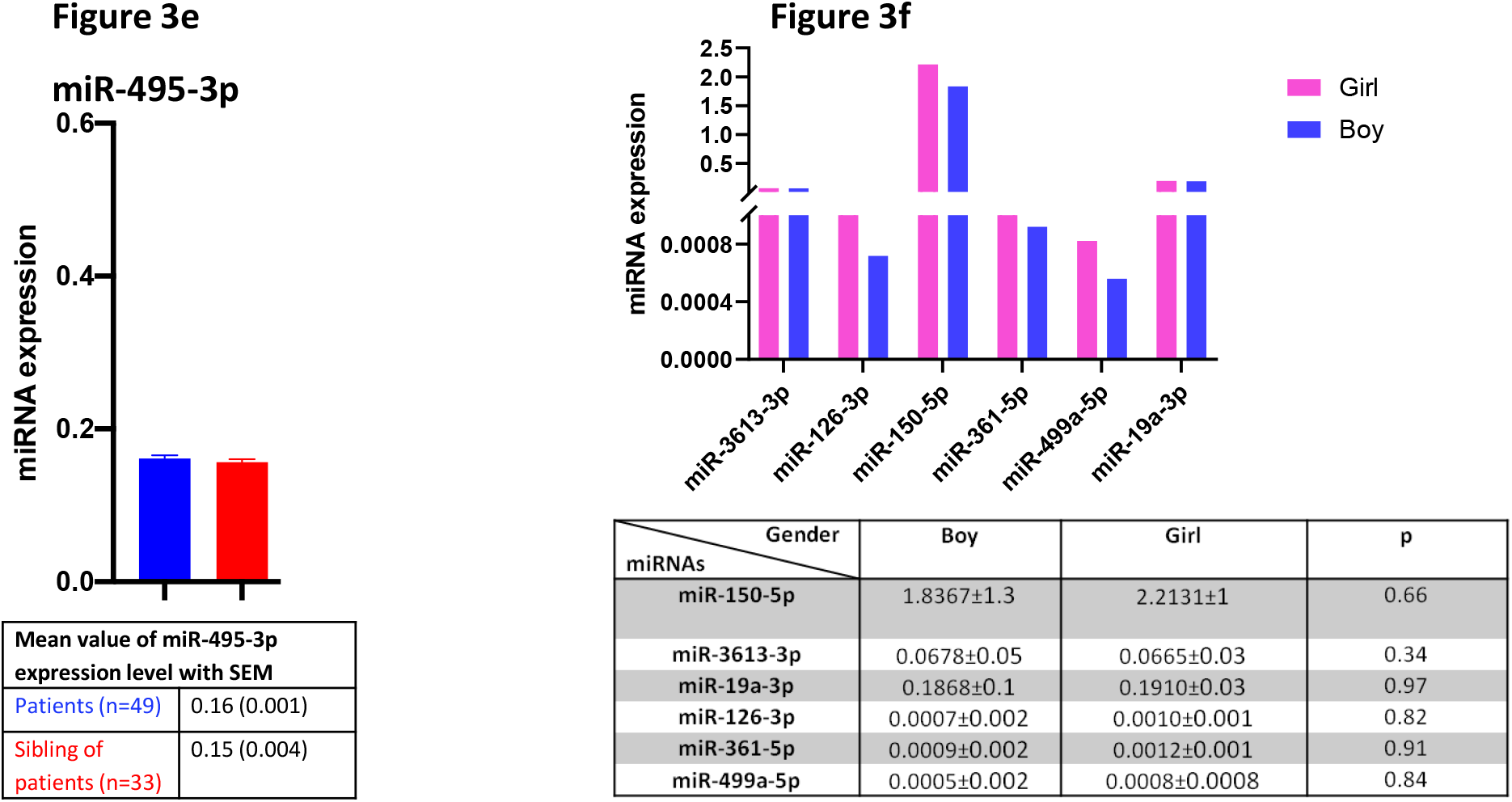
Four examples of comparative distribution profiles of the “six-miRNAs” (miR-19a-3p, miR-361-5p, miR-3613-3p, miR-150-5p, miR-126-3p, and miR-499a-5p) in the same family of patients and their healthy breast siblings. RT-Q-PCR analysis of patients to sibling. Data represents transcription values for “Six-miRNAs” (raw data after normalization). Mean value of miRNAs expression level with standard deviation value (SD). Family 1, 2, 3 and 4 a-d and in e, example of miRNA with unchanged levels between patients and their siblings. Figure 3f - **Decreased expression of the six-miRNAs in patients Girls and Boys with autism**. Distribution of the “six-miRNAs” (miR-19a-3p, miR-361-5p, miR-3613-3p, miR-150-5p, miR-126-3p, and miR-499a-5p) between boy and girl patients. Data represents transcription values for “Six-miRNAs” (raw data after normalization). All raw values from initial screening are presented in Supplementary Tables 1.

In fact, analysis in a given family of each one of the six miRNAs shows that all six miRNAs at the same time are not always at higher levels for the healthy sibling compared to sister or brother patient. Indeed, different combinations are found in each individual familial case and may suggest why some of the siblings did not develop autism, as the six miRNAs are not fully down-regulated to the same level as in the patient.

Examples of the levels of the six miRNAs are shown in Figure 3a-d for four families compared to patients and breast siblings. One of the cases shows in one sibling a down-regulation of one miRNAs while the other five remaining miRNAs are at a higher level compared to the autistic child of the same family (Family 1 in Figure 3a). Another case (Family 2 in Figure 3b) highlights the importance of all six miRNAs at a higher level in two sibling, while in third sibling four of the miRNAs are down-regulated compared to the autistic child. Family 3 (Figures 3c) all six miRNAs are higher in sibling to patient while Family 4 show (Figures 3d) other combinations of distribution of the six miRNAs in the healthy sibling. Figures 3e is a control example of miRNAs that do not vary between siblings and patients with autism. These results show that down-regulation of all six miRNAs is required to express autistic phenotype. Analysis of the healthy sibling of the individual family could help us understand the combination of the necessary levels of the six miRNAs at least in some particular family cases, which may be are required to eventually escape autism. On the other hand, in other families more variations (Figures 3) are observed, these differences again reveal the molecular complexity involved in the disease. In particular, this highlights the variable level distribution of six miRNAs even within the family. Furthermore, these samples of family cases could also provide materials to study the mechanism for the progressive transgenerational silencing events of the six miRNAs.

### Distribution of six miRNAs in autistic boys and girls

Evidence indicates that male gender is a risk factor for ASD^21,22,19^. In Figure 3f the level distribution of six miRNAs (miR-19a-3p, miR-361-5p, miR-3613-3p, miR-150-5p, miR-126-3p, and miR-499a-5p) in the case of boy and girl autistic patients indicates that the average levels of miR-19a-3p and miR-3613-3p are the same in both sexes, while the average levels of four other miRNAs (miR-361-5p, miR-150-5p, miR-126-3p, and miR-499a-5p) are higher in females than in males. Means of statistical deviation is significant with *p* ≤ 0.05. Thus a male sexual bias, which affects the severity of ASD and its possible biological mechanisms, could also be found in the alteration of specific miRNAs.

MiR-150-5p and miR-126 have been reported to be expressed and altered differently in males and females^23^. Hsa-miR-361-5p, in newborn girls, as most significant miRNAs had higher expression^24^. Regarding the remaining miRNAs differences such as the level of miR-499a-5p in girls to boys requires further investigation. Here, we describe the difference in the levels of four miRNAs between boys and girls, to which extent that could help to understand about a sex-dependent trait in autism which requires further investigation. A higher level for these four miRNA transcripts in a girl than in a boy with ASD may suggest common gender-dependent regulation. There are possible biological mechanisms of sex bias that affects the levels of miRNAs, especially with regards to hormones and genetics.

This is the first report of a genetic alteration common to all patients tested in the autism group; and it has been particularly validated in mouse models^13^. However, it is urgent to confirm with separate large cohorts and including the father’s sperm. Early (i.e., very early before parenthood) detection of the autism could possibly be of interest to parents at risk. We assume that the current data will soon either be confirmed in the largest existing patient cohorts, or, failing this, definitively contradicted.

Since there is no reason at this point to suspect the generality of the conclusion, we will consider that ASD is indeed associated with low to very low levels of the “six miRNAs”. However, the initiation event (s) that lead to the coordinated down-regulation of six miRNAs in one generation is far from understood. Consistent with this conclusion are the detection of a partial trait, an intermediate level in close relatives of patients and obviously, by the extensive down-regulation in two of the well-documented animal models (homogenous genome) of the disease.

The down-regulation of six miRNAs is a prominent robust marker for the detection of disorders of the ASD spectrum started in parents; and we now know from mouse models of autism with heterogeneous origins such as chemical and/or genetic that these microRNAs are also down-regulated. In two mouse models of autism we were only able to test five out of six microRNAs (miR-3613 is not present in the mouse genome, but shares most of the miR-19a-3p and miR-499a-5p) and further studies may reveal additional non-coding RNAs also involved in human and mouse models. We do not believe that these six microRNAs are exclusively involved in human and murine models but these six microRNAs are highly plausible marker candidates. In our first human study, we purposely analyzed non-syndromic autistic patients with fear of much heterogeneity; and we excluded from the analysis cases of visible abnormalities of syndromic ASD, with additional phenotypes and dysmorphic features independent of ASD phenotypes (see for the review) ^25^.

Because an independent mutation could be at the origin of multiple variations of gene expression that are may not directly related to autism, it will now be of great interest to know if there are any points of convergence regarding the states of these six miRNAs between patients with syndromic and non-syndromic autism spectrum disorders.

### Inheritance in autism

Individual syndromic autism cases are born with rare *de novo* DNA changes usually associated with chromosomal abnormalities or monogenic mutation^26,27,19,28,29,30,31,29,32,33^. Using genome-wide association studies, it is proposed that in cases of polygenic origin, the common variants which contribute to ASD risks, are a combination of an existing low mutational noise in the parents or an additional *de novo* polymorphism mutation that could lead to the birth of an autistic individual (non-syndromic) in which no additional symptoms are present. Here, following six markers of miRNAs expression, we observe in our cohort of non-syndromic autism, evidences of two stages in successive generations. Thus far we cannot rule out the accumulation of additional low noise mutations responsible for the down-regulation of the six miRNAs in humans; but we do highlight a common mechanism that ultimately leads to their continuous inherited down-regulation.

If the origin of non-syndromic autism has a combination of wild range mutations, then it is difficult to reconcile with stepwise regulation of the six miRNAs (see Ozkul et al.^13^), especially in a genetically homogenous mouse model system. Our data in mice validate the down-regulation of the six miRNAs in non-syndromic autism. The animal model already indicates that males treated with VPA show systematic down-regulation of these six miRNAs, while model analysis of *Cc2d1a+/-* mutants of the six miRNAs shows a systematic modified profile for each of the six miRNAs but with a more complex picture of regulation of descending and/ or rising levels^13^. These results indicate that the levels of the six miRNAs (five in mouse), are sensitive to genetic variation at least in the mouse. We do not yet know, if syndromic ASD patients exhibit modified profiles of the six miRNAs, identical to those of children with non-syndromic ASD.

We hypothesize that the affected non-coding RNAs function cooperatively and are involved in the control of disease-altered functions that are schematically involved in social interactions. Each of the six miRNAs has several hundred targets (Figure 1), which means that many transcripts could be changed at the same time. A lower level of six miRNAs could order cells to be released from cell fate, as their targets are now not degraded but instead maintained. In the event that there is a common regulator for all six miRNAs such as a ncRNA, this should contain at least one sequence complementary to all six miRNAs, either in promotors regions or in transcripts. Their common expression pattern decreases in patients with autism suggests a unique regulatory mechanism from which would derive the expression of distinct patterns of the highly complex genetic systems involved in the control of social and related interactions, as a whole affected by the disease. However, the variable expression pattern in the siblings of patients with autism suggests an alteration already initiated in the parents and partially transmitted to the healthy sibling.

As to the mechanism of the disease and, above all, the mechanisms of the psychological and social interactions affected, these observations could in the future be broadened in two directions starting from the notion that the gene products of the affected miRNAs are themselves regulatory elements. Among the target genes, one can expect to identify genes acting directly in these processes. On the other hand, the common regulation of the six identified miRNA genes may reflect the activities of one or more common upstream regulator(s), the identification of which could hopefully lead us to decipher some of the basic molecular mechanisms responsible for the down-regulation of these six miRNAs.

A possible model of “targeted degradation of miRNAs” (“TDMD”)^14^ is that elicited by other non-coding RNAs such as the long ncRNA induced Cyrano decrease in miR-7 accumulation^34,35^. The transcriptional control, common to the six different miRNAs (and only those), could be at the level of upstream /or downstream transcription will require additional molecular clarification.

In addition, the gradual down-regulation of several miRNAs from parents to children; recall a gene silencing booster (amplification siRNAs) reported in a model system such as plant, mouse or *C. elegans*^36^ with the exception in mammals of the RDRP gene /or activity have yet to be identified. Other mechanisms for persistence of siRNAs are proposed/ or should be identified.

Regardless of the mechanisms of reduced accumulation or alteration of the six miRNAs, it constitutes the hereditary signal which may be the miRNA itself, non-coding RNA or one or more distinct protein(s). In the event that it turns out to be RNA this would be inheritance of a non-Mendelian heterodox type^37^. In support to this last point testes and sperm of mouse models present down-regulation (VPA) and alteration (*Cc2d1a+/-*) of these miRNAs^13^.

In summary, our results and findings, reveal the epigenetic coordinated down-regulation or/ alteration of six distinct miRNAs genes; -two stages of events with progressive down-regulation of six miRNAs: from healthy parents (50% to controls) to healthy children (20% to controls) and to complete autism syndrome in their offspring with expression down to <10% to controls. Distinct profiles of six miRNAs in healthy sibling of children with autism reveal the importance of decreasing the levels of each miRNA.

## Acknowledgments

We thank F. Cuzin for constructive comments on this paper and K. Marcu for English editing of several version.

## Methods

Results of experiments presented and discussed herein are all extracted from data obtained in Ozkul and al. 2020^13^.

We have reported data with the Biomark HD system that provides orders of magnitude higher for real-time PCR compared to conventional platforms using integrated fluidic circuits (IFCs)— nanofluidic circuits containing fluidic networks that automatically combine sets of samples with sets of assays. This innovative solution for real-time PCR provides experiment densities far beyond what is possible with microplate platforms, significantly reducing the number of liquid-handling steps and volumes per reaction^38,39,40,41^. The autoexposure calibration is performed by collecting several images with different exposure times. An optimum exposure time is calculated and is then used for the acquisition of qPCR signals. For this reason, the CCD camera measures continuously in each cycle and thus repeats 40 times in 40 cycles (see method). This method produces robust results.

The real-time PCR step was performed by using a BioMark System with the following protocol: the thermal mixing protocol was followed by heating at 50 °C for 120 sec, 70 °C for 1,800 sec, and 25 °C for 600 sec. Then, the UNG and hot-start protocol were followed by heating at 50 °C for 120 sec and 95 °C for 600 sec. Finally, PCR was performed with 40 cycles at 95 °C for 15 sec and 60 °C for 60 sec. The BioMark system is quantifes low-abundance miRNAs and can detect a single copy at a Ct value of 26–27. Routine qPCR for miR detection and validation (see Supplementary Table 1 for miR sequences) was performed with the miScript PCR control set (catalog number 218380; Qiagen, Germany). The miScript™ miRNA PCR Array Human Serum & Plasma 384HC (Cat No: 331223) was used in this study. The Human Serum & Plasma 384HC miScript miRNA PCR Array profles the expression of 372 miRNAs (see list of the microRNAs in Supplementary Table 3) that are detectable in serum and plasma using the miScript PCR system. SNORD61, SNORD68, SNORD72, SNORD95, SNORD96A, RNU6-2, miRTC, miRTC, and PPC were used as controls. Te data were normalized using the 2−ΔΔct method.

There are some important technical differences between conventional systems using microtiter plates and the microfluidic platforms. The BioMark optical system uses a xenon lamp, optical filters (by default: three out of four filter positions are mounted with filters that are compatible with detection of ROX (6-carboxy-X-rhodamine), FAM (5-carboxyfluorescein), VIC/JOE (6-carboxy4′,5′-dichloro-2′,7′-dimethoxyfluorescein, a charged coupled device (CCD) 9M pixel camera, and a cooling system. The passive reference dye, ROX, is always used in the BioMark platform to identify the chambers and to normalize the signal.

Before the start of cycling, the CCD camera is focused onto the wells. Then, the autoexposure function sets an exposure time for each fluorescence channel based on the background signal. The objective is to set a baseline such that even a minute increase in fluorescence can be detected with high sensitivity by the CCD chip. The autoexposure calibration is performed by collecting several images with different exposure times. An optimum exposure time is calculated and is then used for the acquisition of qPCR signals. For this reason, the CCD camera measures continuously in each cycle and thus repeats 40 times in 40 cycles.

### Data Availability Statement

The datasets generated and analyzed during the current study are available from the corresponding author on reasonable request.

The data underlying this article are available in the article and in its online supplementary material.

#### Ethics Statement

This study was approved by the Hospital Ethics Committee and authorizations from all the patients and the participating relatives were obtained by signing an informed consent form. All parents gave written informed consent before participation (09-20-2011 committee number: 2011/10). All research was performed in accordance with the relevant guidelines and regulations (Erciyes University animal ethics committee 04-11-2012 12/54).

This work was made possible by a grant to Y. Ozkul from Tübitak, 1010 (EVRENA project ID 112S570), and to Minoo Rassoulzadegan Fondation Nestlé Rassoulzadegan 2019-2020.

## Legend of Table and Figures

Supplementary Table 1- Down-regulation of six miRNAs in autistic children and their direct families.

Quantitative RT–PCR was performed by using a high-throughput BioMark Real-Time PCR system (Fluidigm, South San Francisco, CA, USA) see References for all raw data (Ozkul et al.2020). Each value is the CCD camera measures continuously in each cycle and thus repeats 40 times in 40 cycles. The values obtained were normalized by internal control. The expression of 6 serum miRNAs in patients (boys and girls), parents (mothers and fathers) is calculated as % of 37 healthy volunteers. Values are expressed as mean± standard variation. Six-miRNA transcripts (Raw data after normalization) down-regulated profiles in children with autism, sibling and their mothers and fathers compared to age- and sex-matched healthy controls (p<0.0001) (One-way ANOVA tests).

Supplementary Table 2- **Summary of Target Genes for six miRNAs**

Supplementary Table 3- List of 372 microRNAs.

