## Supplementary material for "Progressive decline in the levels of six miRNAs from parents to children in autism": supp tables 1-2-3

**Supplementary Dat**

Table 1a. miRNA expression changes according to their control

| | miR-3613-3p | miR-150-5p | miR-126-3p | miR-361-5p | miR-499a-5p | miR-19a-3p |
|---|---|---|---|---|---|---|
| **Patients (n=45)** | 0.068729 | 1.88095 | 0.001175 | 0.001407 | 0.000916 | 0.181856 |
| | 17.72% | 19.63% | 1.34% | 0.99% | 0.76% | 20% |
| **Patient Boy (n=31)** | 0.059732 | 2.011178 | 0.001262 | 0.001453 | 0.000978 | 0.182325 |
| | 15.40% | 20.99% | 1.44% | 1.02% | 0.81% | 20% |
| **Patient Girl (n=14)** | 0.072791 | 1.822137 | 0.001135 | 0.001386 | 0.000888 | 0.181644 |
| | 18.77% | 19% | 1.30% | 0.97% | 0.73% | 19.99 |
| **Father (n=37)** | 0.14908 | 3.864137 | 0.044443 | 0.056022 | 0.05261 | 0.427877 |
| | 56.66% | 52.8% | 23.3% | 60% | 53.8% | 59.9% |
| **Mother (n=37)** | 0.190171 | 4.844019 | 0.045732 | 0.079835 | 0.065262 | 0.573587 |
| | 49.5% | 46.5% | 16.8% | 42.5% | 39% | 43.1% |
| **Control (n=37)** | 0.38777 | 9.580365 | 0.087165 | 0.141492 | 0.120486 | 0.908365 |

| | miR-3613-3p | miR-150-5p | mi-126-3p | miR-361-5p | miR-499a-5p | miR-19a-3p |
|---|---|---|---|---|---|---|
| **Patient (n=45)** | 0.069±0.05 | 1.900±1.22 | 0.001±0.25 | 0.001±0.05 | 0.001±0.002 | 0.182±0.09 |
| **Patient boy (n=31)** | 0.0678±0.05 | 1.8367±1.3 | 0.0007±0.002 | 0.0009±0.002 | 0.0005±0.002 | 0.1868±0.1 |
| **Patient girl (n=14)** | 2.2131±1 | 2.2131±1 | 0.001±0.001 | 0.0012±0.001 | 0.0008±0.0008 | 0.1910±0.03 |
| **Father Patient (n=37)** | 0.150±0.15 | 3.864±3.84 | 0.044±2.99 | 0.056±0.11 | 0.053±0.09 | 0.428±0.48 |
| **Mother Patient (n=37)** | 0.190±0.19 | 4.844±3.74 | 0.046±0.12 | 0.080±0.06 | 0.065±0.12 | 0.574±0.43 |
| **Control Child (n=21)** | 0.373±0.37 | 10.570±7.06 | 0.066±0.03 | 0.103±0.08 | 0.086±0.05 | 0.817±0.51 |
| **Control Father (n=16)** | 0.344±0.34 | 8.196±4.06 | 0.058±0.03 | 0.140±0.05 | 0.114±0.04 | 1.069±0.47 |
| **Control Mother (n=16)** | 0.380±0.38 | 9.062±3.6 | 0.055±1.56 | 0.139±0.05 | 0.107±0.04 | 1.009±0.45 |

Values are expressed as mean±sd

Six- miRNA transcripts (Raw data after normalization) down-regulated profiles in children with autism, sibling and their mothers and fathers compared to age- and sex-matched healthy controls ($p<0.0001$) (One-way ANOVA tests).

Supplementary Table 1b: **Highest male patient**

| | **miR-3613-3p** | **miR-150-5p** | **miR-126-3p** | **miR-361-5p** | **miR-499a-5p** | **miR-19a-3p** |
|---|---|---|---|---|---|---|
| **Highest male patient** | 0.22677 | 4.60698 | 0.01520 | 0.01699 | 0.01186 | 0.54408 |
| Sibling | - | - | 0.00069 | 0.000951 2 | 0.0005391 | 0.445859 5 |
| Mother | 0.0564882 | 2.543656 | 2.48973 | 0.049558 | 0.0565165 | 1.311286 4 |
| Father | 0.2323707 | 1.879663 3 | 0.01785 | 0.052788 8 | 0.0416584 | 0.811303 8 |
| Highest Control | 0.96770 | 22.93099 | 0.13528 | 0.22768 | 0.22102 | 1.65524 |
| Highest Father Control | 0.55380 | 13.72433 | 0.13292 | 0.26512 | 0.18145 | 2.04072 |
| Highest Mother Control | 0.69384 | 16.73969 | 0.13292 | 0.26512 | 0.18145 | 2.04072 |

Supplementary Table 1c: **Highest female patient**

| | **miR-3613-3p** | **miR-150-5p** | **miR-126-3p** | **miR-361-5p** | **miR-499a-5p** | **miR-19a-3p** |
|---|---|---|---|---|---|---|
| **Highest female patient** | 0.1572 | 4.1346 | 0.0039 | 0.0044 | 0.0031 | 0.2593 |
| Sibling | 0.0992 | 2.7277 | - | - | - | 0.2367 |
| Mother | 0.0711 | 2.3298 | 0.0241 | 0.0472 | 0.0445 | 0.1453 |
| Father | 0.0812 | 1.2989 | 0.0436 | 0.2335 | 0.1843 | 0.1243 |
| Highest Control | 0.9677 | 22.9310 | 0.1353 | 0.2277 | 0.2210 | 1.6552 |
| Highest Father Control | 0.5538 | 13.7243 | 0.1329 | 0.2651 | 0.1814 | 2.0407 |
| Highest Mother Control | 0.6938 | 16.7397 | 0.1329 | 0.2651 | 0.1814 | 2.0407 |

Supplementary Table 1d: **Highest mother patients**

| | miR-3613-3p | miR-150-5p | miR-126-3p | miR-361-5p | miR-499a-5p | miR-19a-3p |
|---|---|---|---|---|---|---|
| Patient | 0.0471441 | 2.404379 | 0.001528 | 0.000744 | 0.0005198 | 0.130406 |
| Sibling | - | 1.173390 | - | - | - | - |
| **Highest mother** | 0.57630 | 17.92144 | 0.19758 | 0.26464 | 0.17495 | 2.10134 |
| Father | 0.2259759 | 0.074709 | 0.001101 | 0.018033 | 0.0200327 | 0.320819 |
| Highest Control | 0.96770 | 22.93099 | 0.13528 | 0.22768 | 0.22102 | 1.65524 |
| Highest Father Control | 0.55380 | 13.72433 | 0.13292 | 0.26512 | 0.18145 | 2.04072 |
| Highest Mother Control | 0.69384 | 16.73969 | 0.13292 | 0.26512 | 0.18145 | 2.04072 |

Supplementary Table 1e: **Highest father patients**

| | miR-3613-3p | miR-150-5p | miR-126-3p | miR-361-5p | miR-499a-5p | miR-19a-3p |
|---|---|---|---|---|---|---|
| Patient | 0.111918 | 2.082017 | 0.00066 | 0.000738 | 0.000516 | 0.163678 |
| Sibling | - | 0.556816 | - | - | - | - |
| Mother | 0.154999 | 4.961163 | 0.08276 | 0.051677 | 0.058853 | 0.308682 |
| **Highest father** | 0.55897 | 17.13459 | 0.88139 | 0.61632 | 0.52845 | 2.26849 |
| Highest Control | 0.96770 | 22.93099 | 0.13528 | 0.22768 | 0.22102 | 1.65524 |
| Highest Father Control | 0.55380 | 13.72433 | 0.13292 | 0.26512 | 0.18145 | 2.04072 |
| Highest Mother Control | 0.69384 | 16.73969 | 0.13292 | 0.26512 | 0.18145 | 2.04072 |

Supplementary Table 1f: **Highest male Sibling patients**

| | miR-3613-3p | miR-150-5p | miR-126-3p | miR-361-5p | miR-499a-5p | miR-19a-3p |
|---|---|---|---|---|---|---|
| Patient | 0.08123 | 2.63195 | 0.00240 | 0.0027 | 0.0019 | 0.112818 |
| **Highest male Sibling** | 0.18791 | 6.01827 | 0.00220 | 0.0025 | 0.0017 | 0.44586 |
| Mother | 0.04919 | 2.06106 | 0.03356 | 0.0660 | 0.1078 | 0.320951 |
| Father | 0.00917 | 0.59173 | 0.00089 | 0.0022 | 0.0007 | 0.46838 |
| Highest Control | 0.96770 | 22.93099 | 0.13528 | 0.2277 | 0.2210 | 1.65524 |
| Highest Father Control | 0.55380 | 13.72433 | 0.06589 | 0.2651 | 0.1815 | 2.04072 |
| Highest Mother Control | 0.69384 | 16.73969 | 0.13292 | 0.2007 | 0.1498 | 1.44065 |

#### Supplementary Table 2

### Summary of Target Genes for miRNAs

**Common targets at least shared between five /or four miRNAs.**

#### Common targets of 5 miRNAs:

- AEBP2 AE binding protein 2
- CFL2 Cofilin 2 (muscle)
- **EIF4E*** Eukaryotic translation initiation factor 4E binding protein 2

#### Common targets of 4 miRNAs:

- **AGO1*** Argonaute RISC catalytic component 1
- **AGO2*** Argonaute RISC catalytic component 2
- AGO3 Argonaute RISC catalytic component 3
- **CDK13*** Cyclin-dependent kinase 13
- **CDK19*** Cyclin-dependent kinase 19
- **DDX3X*** DEAD (Asp-Glu-Ala-Asp) box helicase 3, X-linked
- **FUT9*** Fucosyltransferase 9 (alpha (1,3) fucosyltransferase)
- **IL1RAP*** İnterleukin 1 receptor accessory protein-like 1
- MDM4 Mdm4 p53 binding protein homolog (mouse)
- PLXNC1 Plexin C1
- PTP4A1 Protein tyrosine phosphatase type IVA, member 1
- **PTPRG*** Protein tyrosine phosphatase, receptor type, G
- RAD23B RAD23 homolog B (S. cerevisiae)
- **SLC25A12*** Solute carrier family 25 (aspartate/glutamate carrier), member 12
- TESK2 Testis-specific kinase 2
- **TET3*** Tet methylcytosine dioxygenase 3
- ZBTB10 Zinc finger and BTB domain containing 10

**[1]Autism-related genes demonsrate with star(*)**
**[2]All miRNAs' target genes data were taken from TargetScan Release 7.1 and analyzed.**

**miRNA's Target genes List**

| |
|---|
| **Common target genes of miR-19a-3p, miR-150-5p, miR-361-5p, miR-499a-5p and miR-3613-3p (n=3)** |

| |
|---|
| *Aebp2*, *Cfl2* and *Eıf4e* |

| **Common target genes of miR-19a-3p, miR-361-5p, miR-499a-5p and miR-3613-3p (n=7)** |
|---|
| *Ago2*, *Cdk19*, *Ddx3x*, *Fut9*, *Ptprg*, *Slc25a12* and *Zbtb10*. |

| **Common target genes of miR-19a-3p, miR-150-5p , miR-499a-5p and miR-3613-3p (n=6)** |
|---|
| *Ago3*, *Cdk13*, *Il1rapl1*, *Plxnc1*, *Ptp4a1* and *Tesk2* |

| **Common target genes of miR-19a-3p, miR-150-5p, miR-361-5p and miR-3613-3p (n=4)** |
|---|
| *Ago1*, *Mdm4*, *Rad23b* and *Tet3*. |

| **Common target genes of miR-19a-3p, miR-499a-5p and miR-3613-3p (n=81)** |
|---|
| *Abcb7, Ankrd13c, Arıd5b, Asb7, Atf2, Bnc2, Btg1, C5orf24, Ccrn4l, Celf1, Celsr1, Cnot6l, Cntnap5, Crebzf, Csnk1g1, Csrnp3, Dcp2, Dcun1d1, Ddx3y, Dnal1, Ebf1, Eea1, Eıf4e3, Eıf4g2, Elmod2, Evı5, Fem1c, Fndc3a, Foxp2, Gmfb, Gpbp1, Hspa4l, Ino80, Insıg1, Kctd16, Kıaa1468, Klf7, Klhl42, Kpna3, Lın28b, Lrch2, Map3k2, Mapk6, Med13, Msı2, Nlk, Nrarp, Orc2, Pde3b, Phf17, Prkaa2, Pten, Qkı, Rab21, Rbms3, Reep1, Samd8, Satb1, Secısbp2l, Sh3kbp1, Shcbp1, Sıx1, Slc35f1, Sorl1, Sox4, Sox5, Sox6, Srpk2, Syt1, Tacc1, Thrb, Tmbım6, Trerf1, Ubn1, Ubn2, Vcpıp1, Vkorc1l1, Yıpf6, Yy1, Zdhhc2* and *Znrf2*. |

| **Common target genes of miR-19a-3p, miR-361-5p and miR-3613-3p (n=42)** |
|---|
| *Aak1, Arhgef12, Bmpr2, Cbfa2t2, Cds1, Csmd1, Ctdspl2, Dcun1d3, Dnmt3a, Dync1lı2, Erbb4, Etv3, Gnptg, Hcfc2, Homer1, Igsf10, Ino80d, Kpna4, Lmbr1, Lnpep, Mecp2, Mef2d, Mex3a, Mllt6, Nfıa, Nhs, Nkx6-1, Phax, Prune2, Setd9, Syncrıp, Tgfbr2, Tmem106b, Tmem64, Tsc1, Uhrf1bp1l, Ulk2, Zbtb18, Zhx3, Zmat3, Znf516* and *Znf704*. |

| **Common target genes of miR-361-5p, miR-499a-5p and miR-3613-3p (n=18):** |
|---|
| *Arglu1, Cdyl2, Cmc1, Dcun1d5, Gja9, Gpr180, Larp4b, Lrıg2, Mtx3, Mycbp, Ppm1d, Sh3bgrl2, Slc24a2, Slc41a1, Tmem178b, Ube2v2, Wdr41* and *Znf24*. |

| **Common target genes of miR-19a-3p, miR-150-5p and miR-361-5p (n=1)** |
|---|
| *Prıckle2*. |

| **Common target genes of miR-150-5p, miR-361-5p and miR-3613-3p (n=8)** |
|---|

| |
|---|
| *Ahsa2, Gpr75, Ipo9, Pcnp, Pde7a, Pkp4, Sp1* and *Tenm4.* |

| **Common target genes of miR-19a-3p, miR-150-5p and miR-3613-3p (n=40)** |
|---|
| *Abi2, Adipor2, Arih2, Azin1, Cast, Ccnd2, Cdc73, Clcn6, Cpd, Cpeb4, Cyb561d1, Dkk3, Dsel, Dynll2, Fam179b, Fbxw11, Fosl2, Foxp1, Hif1a, Hnrnpu, Itgb3, Klhl3, Map1b, Map3k12, Nbea, Nfic, Pcdh1, Pik3cb, Rasgef1a, Rimkla, Rims2, Srgap3, Usp13, Vmp1, Vps53, Vsig10, Zbtb20, Zbtb4, Zc3h10* and *Zfp91.* |

| **Common target genes of miR-19a-3p, miR-150-5p and miR-499a-5p (n=1)** |
|---|
| *Pfn2.* |

| **Common target genes of miR-150-5p, miR-499a-5p and miR-3613-3p (n=14)** |
|---|
| *Apc, C6orf120, E2f3, Foxo4, Fzd4, Kcnn3, Mmp16, Pdcd4, Prkar1a, Ptbp2, Srcin1, St3gal6, Stat1* and *Zswim6.* |

| **Common target genes of miR-19a-3p, miR-126-3p and miR-3613-3p (n=1)** |
|---|
| *Itga6.* |

| **Common target genes of miR-126-3p and miR-3613-3p (n=13)** |
|---|
| *Adam9, Herpud1, Kank2, Slc7a5, Fbxo33, Plk2, Crk, Irs1, Dip2c, Efhd2, Gna13, Gbp2* and *Akap13.* |

| **Common target genes of miR-19a-3p and miR-126-3p (n=1)** |
|---|
| *Camsap1.* |

| **Common target genes of miR-499a-5p and miR-3613-3p (n=221)** |
|---|
| *Abi1, Acvr2b, Adam10, Adcyap1, Akr1d1, Ankrd40, Ap1g1, Arid2, Armc2, Atad1, Atrn, Bcl11b, Bhlhe41, C1d, C20orf112, Cacnb2, Capn7, Ccdc50, Ccdc71l, Cd38, Cdc27, Cdc6, Cdca4, Cdh2, Cdk14, Cdk6, Chd9, Chl1, Cisd2, Cldn11, Clybl, Cmip, Cntnap2, Col4a3, Commd2, Cpsf2, Crp, Dcc, Ddx4, Derl2, Dgcr8, Dld, Dnah3, Dsn1, Dyrk2, Efnb1, Eif3j, Eif5, Elavl4, Eml4, Enah, Epb41l2, Epha3, Epha5, Ero1l, Esrrg, Ets1, Eya4, Fam102b, Fam110b, Fam131b, Fam204a, Fam217b, Fam60a, Fam63b, Fam76a, Faxc, Fbxl3, Fbxl4, Fbxo42, Fbxw7, Fgf2, Fgf9, Fkbp5, Fnbp4, Fnip1, Fnip2, Foxg1, Fyn, Fzd8, Gabra1, Gpc6, Greb1, Gxylt1, H3f3b, Haus6, Hic2, Hlcs, Hook3, Hoxd9, Hp1bp3, Hspa4, Iffo2, Ikzf2, Ikzf5, Impg2, Inip, Insr, Irf2bpl, Irx5, Itgbl1, Jag1, Jhdm1d, Jph1, Kiaa1614, Kif26b, Kif5b, Klhl15, Lamp3, Lcor, Lmcd1, Lmnb2, Lmx1a, Lrp8, Lrrc8a, Lrrc8c, Lrrtm1, Mapre2, Marcks, Med7, Meis1, Minos1, Msl1, Mtdh, Mtmr3, Myef2, Mylk, Nav2, Nipal1, Nova1, Nrip1, Pacs2, Paip2b, Pank1, Papola, Pbrm1, Pde10a, Pip5k1c, Pogz, Ppil4, Ppp3ca, Ppp3cb, Ppp6c, Prkab2, Prmt3, Proser2, Prr18, Psd3, Ptar1, Ptbp3, Ptpdc1, Ptpn4, Purb, Rab11fip4, Rab22a, Rab3c, Rad54l2, Rasgrp3, Rcan3, Rfx7, Rims1, Rmnd5b, Rnf114, Rnf115, Rnf152, Rsbn1, Rtkn2, Rybp, Scn2b, Scrg1, Sec62, Sertad2, Setbp1, Sgip1, Sh3bp5, Slc12a2, Slc30a4, Slc7a6, Smc6, Smim15, Snrk, Snx2, Stau2, Stc1, Syap1, Tanc1, Tardbp, Tbc1d15, Tbc1d22b, Tcf12, Tgfbr1, Tle4, Tmem189, Tmem189-Ube2v1, Top1, Tox3, Trim13, Trim67, Tshz2, Tspan12, Tspan6, Ube2n, Ube2nl, Ube2v1, Ube2w, Usp24, Usp47, Vav3, Vgll2, Wdr82, Wwc2, Xkr4, Xrn1, Ybx1, Yipf4, Zdhhc13, Zic3, Znf19, Znf280c, Znf281* and |

| |
|---|
| *Znf518b.* |

| **Common target genes of miR-19a-3p and miR-499a-5p (n=12)** |
|---|
| *Acbd5, Dennd4c, Eıf4a2, Fam168a, Ppfia2, Rnf145, Rnf44, Sec11a, Sos2, Steap2, Sybu and Thra* |

| **Common target genes of miR-19a-3p and miR-3613-3p (n=827)** |
|---|
| *Abca1, Abcd3, Abhd17c, Abhd5, Abr, Acer2, Acp1, Acsbg1, Acsl4, Adam12, Adamts17, Adamts18, Adcy9, Add3, Adnp, Adrb1, Aff1, Aftph, Agpat5, Aıda, Ak3, Akap1, Al590483.1, Alg14, Ammecr1l, Ankrd12, Ankrd29, Ano3, Apold1, Appl1, Ar, Arfgef1, Arfgef2, Arfip1, Arhgap11a, Arhgap12, Arhgap5, Arhgef26, Arıd1a, Arıd4b, Arl8a, Armc8, Arpp19, Asap2, Asxl2, Atg14, Atg16l1, Atp2b2, Atp2c1, Atp6v1b2, Atrx, Atxn1, Atxn1l, Atxn7, Atxn7l1, B3galnt2, B3galt5, B4galt5, Bag5, Bambı, Barhl2, Bcl2l11, Bcl7a, Bend3, Bend4, Bıcd1, Blcap, Bmp2k, Bmp3, Bnıp2, Bptf, Brwd1, Brwd3, Btaf1, Btbd7, Btf3l4, C11orf30, C11orf57, C12orf66, C16orf52, C19orf40, C20orf194, C21orf91, C2orf42, C3orf70, C5orf30, C6orf222, C7orf60, C9orf69, Cab39, Cacna1c, Cacul1, Cadm2, Cald1, Calm1, Camsap2, Cand1, Caprın1, Caprın2, Casz1, Cbfb, Cblb, Cbln2, Cbx6, Ccdc126, Ccdc144a, Ccdc149, Ccdc152, Ccdc6, Ccer1, Ccnd1, Ccnl1, Ccnt2, Ccser2, Cd164, Cd84, Cdc42bpa, Cdh11, Cdh20, Cecr2, Cep170, Cep350, Chd5, Chıc1, Chst1, Chst7, Clıp1, Clıp4, Clock, Cnga3, Cnksr2, Cnot6, Cnot7, Cnr1, Col19a1, Coq10b, Coq6, Cox15, Cox8c, Cpeb1, Cpeb3, Cr2, Crcp, Crebl2, Crebrf, Crmp1, Cs, Cse1l, Csf2rb, Ctgf, Cul5, Cyld, Dag1, Dbn1, Dcaf7, Dcaf8, Dcbld2, Dda1, Ddhd1, Ddı2, Ddx6, Dennd6a, Dgke, Dhx33, Dhx40, Dıcer1, Dıras2, Dkfzp779j2370, Dlc1, Dlg5, Dmxl2, Dnaja2, Dnajb1, Dnajb14, Dnajc16, Dock4, Dock5, Dscam, Dtna, E2f8, Ebf2, Ednrb, Efna5, Efr3a, Egf, Egr3, Ehbp1, Elk3, Ell2, Elovl5, Emc7, Emx2, Enc1, Enpp5, Eogt, Epc1, Epc2, Epg5, Epgn, Epn2, Erbb3, Ereg, Esr1, Etnk1, Etv1, Etv5, Exoc5, Exoc6b, Ext1, F3, Fam104a, Fam114a1, Fam126b, Fam155a, Fam160a1, Fam20b, Fam43a, Fam46a, Fam46b, Fam46c, Fam73a, Fam76b, Fam83d, Fam83f, Fas, Fat3, Fbxo28, Fbxo32, Fbxo36, Fbxo48, Fgf10, Fhl5, Fıcd, Fkbp15, Fmr1, Fnbp1l, Fndc3b, Fras1, Frk, Frmd6, Frs2, Fut4, Fxr1, Fzd3, Fzd6, G2e3, G3bp2, Gabarapl1, Gabrb2, Galnt1, Gapvd1, Garem, Gatad2b, Gatsl2, Gda, Gfpt1, Ghr, Gıgyf1, Gıt2, Gja1, Glccı1, Glra3, Gltp, Gnaq, Gnb1l, Golga8f, Golga8g, Gpr153, Gprın3, Gramd1b, Grb10, Grın2a, Grsf1, Gtf2a1, Gtf2h1, Gulp1, Havcr1, Hdac4, Hectd3, Hecw2, Heg1, Hey2, Hgf, Hhıp, Hıc1, Hıp1, Hıpk1, Hıpk3, Hlf, Hmgcs1, Hnrnpa1, Hnrnpd, Hnrnpf, Hnrnpul1, Hnrnpul2, Hoxa5, Hoxa9, Hoxd1, Hrk, Hs2st1, Hspa2, Hspbap1, Id2, Id4, Igf1, Igf2r, Igfbp3, Igsf3, Ikzf1, Il1r1, Il6st, Inhba, Insıg2, Ipmk, Ireb2, Itch, Itga11, Itga2, Itgb8, Itpkb, Itprıpl2, Jarıd2, Kalrn, Katnal1, Kazn, Kbtbd8, Kcnh1, Kcnh5, Kcnj2, Kcnj6, Kcnk10, Kcnk2, Kcns2, Kcnv1, Kdm2a, Kdm2b, Kdm6a, Kıaa0101, Kıaa0355, Kıaa1211, Kıaa1217, Kıaa1244, Kıaa1462, Kıaa1598, Kıaa1671, Kıaa2013, Kıf13a, Kıf1b, Kıf3a, Kıt, Klf10, Klf12, Klf13, Klf2, Klf6, Klhl11, Klhl20, Klhl28, Knstrn, Kntc1, Kpna6, Kras, L3mbtl3, Lbh, Lclat1, Ldlr, Lgalsl, Lhx9, Lımch1, Lın9, Lmln, Lonrf3, Lpgat1, Lphn2, Lphn3, Lpp, Lppr1, Lrch1, Lrıg3, Lrp12, Lrrc16a, Lrrc40, Lrrc8b, Lrrk1, Lrrtm2, Lsamp, Lsm12, Luzp1, Lzıc, Macf1, Maml1, Man1c1, Map2k3, Map3k1, Map4k3, Mapk1, Mapk14, Mapk8, Marc1, Mark2, Mastl, Mb21d2, Mbd2, Mbnl1, Mbnl3, Mcc, Mctp1, Mdfic, Mecom, Med12l, Mef2c, Memo1, Mfap3l, Mfsd6, Mfsd9, Mgat5, Mıb1, Mıcal2, Mıd1, Mıd1ıp1, Mıer1, Mıer3, Mıpol1, Mkl2, Mlec, Mllt10, Mltk, Mmgt1, Mn1, Mob1b, Mpped2, Mprıp, Mrpl17, Msmo1, Mst4, Mtf2, Mtmr10, Mtmr12, Mtmr6, Mxd1, Mybl1, Mycn, Mypn, Myt1l, Naaladl2, Nacc1, Nacc2, Nanp, Nav3, Nbeal1, Ncald, Nckap1, Nckap5, Nckıpsd, Ndfip2, Ndnf, Nek10, Nek9, Neurl1b, Neurod1, Nf1, Nfatc2, Nfib, Ngdn, Nhsl2, Nkrf, Nmt2, Nol4, Npas2, Npas3, Nptn, Npy2r, Nr3c2, Nrk, Nuak1, Nudcd3, Nufip2, Nup37, Nup54, Nus1, Onecut1, Osbpl6, Otud1, Otud7b, Pacs1, Papd4, Parm1, Parp8, Parvb, Patl1, Pax6, Pcbd2, Pcdh10, Pcgf5, Pde4a, Pde5a, Pdık1l, Pea15, Pex5l, Pfkfb3, Pfn1, Pgk1, Pgm2l1, Phactr4, Phf12, Phf13, Phf14, Phıp, Phlda1, Phlda3, Phlpp2, Phtf2, Pı15, Pık3c2a, Pık3ca, Pık3r3, Pıkfyve, Pıtpnm2, Pknox1, Plaa, Plag1, Plcl1, Plxdc2, Plxna4, Pmepa1, Pnrc1, Pon2, Pou2f1, Pou3f2, Pou4f1, Pp13439, Ppara, Ppıg, Ppm1a, Ppp1r12a, Ppp1r13l, Ppp1r15b, Ppp2r5a, Ppp2r5e, Ppp4r4, Ppp6r2, Ppp6r3, Pptc7, Prc1, Prkaa1, Prkacb, Prkar2a, Prox1, Prr14l, Prrt3, Psd, Ptchd4, Ptges3, Ptpn14, Ptpn21, Ptprd, Purg, R3hdm2, Rab14, Rab18, Rab23, Rab2b, Rab33b, Rab4a, Rab5b, Rab8b, Rad51d, Raı2, Ralgps1, Ralgps2, Rap1a, Rap1b, Rap2b, Rap2c, Rapgef2, Raph1, Rasa1, Rassf2, Rbbp8, Rbfox2, Rbm20, Rbms1, Rcor3, Reep3, Rest, Rfx4, Rgmb, Rıc3, Rıctor, Rın2, Rlım, Rnf11, Rnf111, Rnf165, Rnf2, Rnf38, Ror1, Rora, Rp11-676j12.7, Rp1-170o19.20, Rp11-766f14.2, Rpap3, Rprd2, Rps24, Rps6ka2, Rps6ka5, Rragc, Rragd, Runx1t1, Rxra, S100pbp, S1pr1, S1pr2, Samd4a, Sar1a, Sar1b, Sbf2, Scarf1, Scml2, Scn2a, Scn3a, Scn4b, Scn8a, Scube3, Sdc1, Sec22a, Sec61a1, Sec61a2, Sec63, Sept7, Serınc3, Sesn3, Sfmbt1, Sgcb, Sgcd, Sgk1, Sgms2, Sh3d19, Sh3pxd2a, Sh3rf3, Shc3, Shroom3, Sık3, Sıpa1l1, Sırt5, Skıda1, Skıl, Slc16a7, Slc17a6, Slc22a3, Slc24a3, Slc24a4, Slc25a34, Slc26a4,* |

| |
|---|
| *Slc26a7, Slc2a4rg, Slc30a7, Slc31a2, Slc35c2, Slc35d1, Slc35e2b, Slc48a1, Slc6a11, Slc6a6, Slc8a1, Slc9a2, Slc9a6, Smad5, Smarca2, Smarcd2, Smarce1, Smcr8, Smg1, Smurf2, Snip1, Sntg1, Snx18, Snx27, Socs3, Socs5, Socs6, Sowahb, Sox21, Spata12, Spata2, Spats2l, Spen, Spg20, Spire1, Spock1, Spred2, Spry3, Sptssa, Srpx2, Srsf6, St3gal5, St8sia4, Stam2, Stambpl1, Stk35, Stox2, Strn3, Sulf1, Suz12, Sv2c, Synm, Synpo2, Syt11, Syt2, Syt6, Taok1, Tars, Tbc1d12, Tbc1d25, Tbc1d8, Tbcel, Tbk1, Tbr1, Tbx18, Tceb3, Tfcp2l1, Tfdp2, Tgfbrap1, Tgif1, Tgm5, Tgoln2, Thbs1, Thsd7a, Tia1, Tm9sf3, Tmem110, Tmem132b, Tmem136, Tmem159, Tmem167b, Tmem196, Tmem236, Tmem245, Tmem47, Tmem59, Tmem63b, Tnfrsf1b, Tnks, Tnpo2, Tnrc6a, Tnrc6b, Tnrc6c, Tor1b, Tp53inp1, Tpcn1, Tprg1, Tprg1l, Trabd2b, Trak2, Trim33, Trpm3, Trpm6, Ttc9, Ttyh3, Tub, Txk, Txlna, Txlng, Ube2a, Ube2d2, Ube2d3, Ubl3, Ubxn2b, Ucp3, Uhmk1, Unc13a, Usp12, Usp33, Usp37, Vamp3, Vangl1, Vapa, Vat1l, Vav2, Vgll3, Vgll4, Vps35, Vps37a, Vps37b, Vti1a, Wac, Wbp1l, Wdfy2, Wdfy3, Wdr44, Wee1, Wipf3, Wnk1, Wnk3, Wnt3, Wwp1, Xylt1, Yaf2, Yes1, Yipf5, Ythdc1, Ythdf2, Zbtb14, Zbtb39, Zbtb46, Zbtb47, Zdhhc7, Zer1, Zfand5, Zfand6, Zfyve26, Zfyve9, Zic5, Zmynd11, Znf217, Znf367, Znf445, Znf644, Znf654, Znf70, Znf710, Znf772, Znf827, Znf831, Znrf3, Zpld1* and *Zranb2*. . |

| **Common target genes of miR-361-5p and miR-3613-3p (n=155)** |
|---|
| *Acap2, Adat2, Adcy2, Agps, Ajap1, Aldh1l2, Alg9, Ankfy1, Arhgap20, Asb11, Atl3, Atp6v0a2, Atpaf1, B4galt6, Bcorl1, Bdp1, C11orf44, C15orf61, C16orf87, Calcrl, Camk2d, Ccdc169-Sohlh2, Ccdc30, Ccdc58, Ccny, Cdh1, Cdk17, Cenpbd1, Cisd1, Cldn22, Cldn8, Cpox, Crebbp, Cstf1, Ctbs, Ctnnd1, Dctn6, Ddx52, Diaph2, Dpp10, Dr1, Dyrk3, Ehf, Elovl7, Entpd7, Epb41, Ercc6, Exd2, Fam213a, Fgf7, Fnbp1, Gcom1, Gins2, Glipr1, Gpr155, Gpr22, Gpr63, Grip1, Gsto2, Gtdc1, Gtf2e1, Hmgb2, Hmgcr, Hsp90aa1, Ipo7, Isca2, Kansl1l, Kcmf1, Kcnj3, Kdelc2, Kiaa1549l, Kif5a, Klhl18, Klhl7, Kpnb1, Lpcat1, Lppr5, Luc7l, Lypd6, Map2, Map3k9, Met, Mrps23, Mxi1, Myh9, Nebl, Nfat5, Nfix, Otud4, P4ha2, Pabpc4l, Parp11, Pde4b, Pdk4, Perp, Pgm3, Pik3r1, Pkia, Polr2m, Polr3g, Psmd1, Qsox1, Rap2a, Rassf9, Rcan2, Rchy1, Rhoa, Rnf8, Rqcd1, Rwdd1, Samd12, Sc5d, Scaf8, Scai, Sdk1, Sestd1, Shoc2, Slc16a10, Slc6a5, Smc2, Smim14, Snap25, Snd1, Sohlh2, Spry1, Srek1, St6gal2, St6galnac5, Stx7, Tfg, Tgfb2, Thap1, Timp4, Tmem56, Traf3, Trove2, Twist1, Uba6, Ube2h, Ube2k, Ubr5, Vapb, Vegfa, Vwde, Wdr12, Wnt7a, Wt1, Xkr7, Zbtb24, Zbtb38, Zbtb40, Zbtb8b, Zdhhc17, Zmym4* and *Znf507*. |

| **Common target genes of miR-19a-3p and miR-361-5p (n=3)** |
|---|
| *Zfpm2, Acadsb* and *Mtus1*. |

| **Common target genes of miR-361-5p and miR-499a-5p (n=1)** |
|---|
| *Rragb*. |

| **Common target genes of miR-150-5p and miR-361-5p (n=5)** |
|---|
| *Exo5, Rock1, Capn6, Gid4* and *Ghsr*. |

| **Common target genes of miR-150-5p and miR-3613-3p (n=189)** |
|---|

| |
|---|
| Acbd3, Actr1a, Actrt3, Adam19, Adarb1, Aıfm2, Akırın1, Akt3, Angel1, Ankrd52, Arntl2, Atp8a2, Basp1, Brd1, Btbd9, C3orf72, Cacna1g, Calcr, Calu, Capzb, Cbl, Cbx8, Cc2d1b, Ccdc174, Ccdc73, Ccnt1, Celf3, Cep128, Cnnm2, Csnk1a1, Ctdsp2, Ctnnb1, Cxorf23, Cytıp, Dcaf6, Dennd4a, Dgkı, Drg1, Dst, Dyrk1a, Ebf3, Egr2, Eıf4b, Elmsan1, Elovl3, Ensa, Entpd1, Ep300, Ephb2, Erp44, Etf1, Fahd2a, Fam117b, Fam134c, Fam49a, Fbln7, Fıgn, Foxd3, Fto, Gab1, Gabra4, Gan, Gata6, Glg1, Glıs3, Gosr1, Gpam, Grhl2, Gsk3b, Gtf3c2, H1f0, Hsp90b1, Igf2bp1, Igfbpl1, Irf2bp2, Jph2, Kcnb1, Kctd9, Kıaa0087, Letmd1, Lıx1l, Lrrc58, Map2k6, Map3k3, March6, Mbtd1, Metap1, Mga, Mlxıp, Mmp14, Mpp6, Mtch2, Mtmr9, Muc21, Myb, Myo1c, Myocd, N4bp2l1, Ncs1, Ndc1, Negr1, Nfasc, Nfatc3, Nkx2-4, Nlgn3, Nova2, Nr1d2, Nr2f2, Onecut2, Onecut3, Pafah1b1, Pald1, Papd5, Pappa, Parl, Parva, Pax5, Pcp4l1, Pdha1, Peg10, Phb2, Pıgg, Pık3ap1, Pkhd1, Plxnb1, Prkca, Prps1, Psmd11, Ptgfr, Ptprb, Pura, Rab11a, Rab9a, Rad51, Rc3h1, Rcsd1, Reps2, Rnaseh2c, Rorb, Rp6-24a23.6, Rpp14, Rsbn1l, Rsf1, Scn3b, Sephs1, Sgms1, Skp1, Slc30a5, Smarcd1, Smu1, Soga3, Spred3, Stmn4, Stx3, Styxl1, Suv39h2, Szrd1, Tada1, Tango2, Tbc1d20, Tcf23, Tmcc1, Tmem107, Tmem134, Tmem164, Tmem41a, Tmem55a, Tpm3, Trım66, Trmt1l, Trps1, Tspyl5, Uap1, Ube2r2, Ubfd1, Ubr1, Usp49, Ust, Wdr5b, Wnt2b, Wtap, Zbtb7a, Zbtb7c, Zeb1, Zmat2, Znf189, Znf226, Znf317 and Znf574. |

| **Common target genes of miR-19a-3p and miR-150-5p (n=8)** |
|---|
| Bsn, Chd2, Cısd3, Itsn1, Mbd6, Ppp2r4, Rsph10b and Sema6d. |

| **Target genes of miR-19a-3p (n=291)** |
|---|
| Actn1, Adamtsl3, Adss, Al020996.1, Alg2, Alpk2, Ankıb1, Ankrd34a, Ano1, Anxa7, Aqpep, Arap2, Arc, Arhgap1, Arrdc3, Arrdc4, Asıc4, Asna1, Atg4d, Atp10a, Atp11a, Atp1a2, B4galt2, Bace1, Bcl3, Bıcc1, Bod1l2, Bzrap1, C11orf96, C16orf70, C22orf15, C22orf31, C2cd5, C6orf223, C8orf4, Cab39l, Cadm4, Card10, Cbx7, Cc2d1a, Ccdc88a, Ccm2, Ccna2, Cep55, Cgn, Chmp4b, Chrnb2, Clcn2, Cltc, Cntfr, Col1a2, Cpped1, Csnk2a1, Cspp1, Dennd1a, Dhrs3, Dıp2a, Dırc2, Dlx1, Dlx3, Dock10, Dock3, Dpysl5, Dusp7, Dut, Edaradd, Efnb2, Egflam, Epb41l5, Ephb3, Ephx4, Eps15, Evı5l, Eya1, Fam102a, Fam177b, Fam69a, Fam73b, Fam83g, Fastk, Fbxo10, Fbxo40, Fbxo8, Fgf6, Fıbın, Flnc, Foxd4, Foxd4l1, Foxd4l2, Foxf2, Frmpd2, Furın, Gatsl3, Gga1, Glıs2, Golga6a, Golga6b, Golga6c, Golga6d, Golga8a, Golga8b, Gpcpd1, Gpr137b, Grk6, Grm4, Gsc, Gskıp, Hapln4, Hbp1, Hdac7, Hdlbp, Hoxd4, Hprt1, Hs3st5, Il1rap, Ildr1, Impdh1, Inhbb, Irx4, Itfg3, Ivns1abp, Jakmıp1, Jazf1, Kcna4, Kcnc4, Kcnq5, Kıaa2026, Kıf13b, Klhl17, Lemd3, Lıf, Lımd2, Llgl2, Lonrf1, Lrıg1, Magı2, Mbnl2, Med13l, Med26, Mef2a, Mıcal3, Mınk1, Mon2, Mylıp, Napb, Nbeal2, Ncf1, Neurod4, Neurog1, Nme7, Npepl1, Nr1h3, Nrbf2, Nrbp1, Nrd1, Nudt13, Ocrl, Pak6, Pcdh12, Pcdha1, Pcdha10, Pcdha11, Pcdha12, Pcdha13, Pcdha2, Pcdha3, Pcdha4, Pcdha5, Pcdha6, Pcdha7, Pcdha8, Pcdha9, Pcdhac1, Pcdhac2, Pde7b, Phf20, Pı4ka, Pıanp, Pıtx1, Pla2g10, Plcb1, Plcl2, Postn, Ppp6r1, Prkx, Prr5l, Psap, Ptk2b, Pxdn, Rab13, Rab1a, Rab34, Rabgap1, Racgap1, Raf1, Rapgef4, Rapgefl1, Rdh12, Rfx1, Rgl1, Rgp1, Rgs21, Rhebl1, Rhob, Rnf130, Rnf167, Rnf19a, Rnf216, Robo2, Rtn1, Rtn4rl1, Runx3, Scn1b, Sec24d, Sel1l3, Sema4c, Sın3b, Sıva1, Slc12a7, Slc25a44, Slc25a6, Slc27a1, Slc2a1, Slc37a1, Slc6a8, Slc9a1, Slıt1, Slmap, Smoc1, Smoc2, Smurf1, Snph, Snx17, Socs1, Soga2, Sox12, Sphk2, Spryd3, Sptssb, Spty2d1, Stk33, Stk38, Stk39, Stx12, Stx18, Sun1, Sv2a, Syn1, Taf4, Tgm2, Tgm3, Tıparp, Tln2, Tmem151b, Tmem25, Tnfaıp3, Tnfrsf12a, Tnıp1, Tom1l2, Trım23, Trpc3, Tshz3, Tspan18, Ubap2l, Uhrf2, Unc93b1, Usp6, Usp8, Vps4b, Vsıg10l, Vstm2b, Wbp2, Wbp4, Wdr1, Wdr20, Wdr26, Wdr45b, Wnt1, Wnt10a, Wnt7b, Wrnıp1, Zbtb11, Zbtb7b, Zdhhc18, Zdhhc23, Znf469, Znf521, Znf609, Znf711, Znf800 and Znf804a. |

| **Target genes of miR-361-5p (n=48)** |
|---|
| Ac010327.2, Bdnf, Bend6, Brsk2, Btf3, C10orf128, Calm3, Ccdc178, Cd177, Cısh, Cks1b, Cngb3, Cpsf7, Erc2, Erg, Fadd, Foxm1, Fsd2, Golga4, Gphn, Gucy1b3, Helt, Il10, Il13ra1, Kcnn1, Lad1, Lurap1l, Nfe2, Nr4a2, Ogn, Panx2, Rab3gap1, Rad23a, Ranbp17, Rnf4, Rpp25, Sertm1, Slc25a21-As1, Slc7a6os, Tfap2b, Tgfb1, Tktl2, Tmem184a, Trım44, Txndc2, Vegfb, Vezf1 and Wasf2. |

| Target genes of miR-3613-3p (n=6988) |
|---|
| *A1bg, A1cf, Aacs, Aadacl3, Aaed1, Aagab, Aar2, Aard, Aasdhppt, Aass, Abca13, Abca5, Abca6, Abca8, Abca9, Abcb1, Abcb10, Abcb11, Abcb5, Abcc1, Abcc4, Abcc5, Abcc6, Abcd2, Abce1, Abcf1, Abcf3, Abcg1, Abcg2, Abcg8, Abhd10, Abhd13, Abhd15, Abhd17a, Abhd17b, Abhd2, Abhd6, Abl1, Abl2, Abracl, Ac004899.1, Ac005003.1, Ac005477.1, Ac005481.5, Ac006486.1, Ac007390.5, Ac007431.1, Ac007952.5, Ac007952.6, Ac008443.1, Ac008964.1, Ac011755.1, Ac011897.1, Ac012123.1, Ac012215.1, Ac012360.2, Ac016559.1, Ac024257.1, Ac026310.1, Ac068987.1, Ac074091.13, Ac079210.1, Ac079602.1, Ac105020.1, Ac106017.1, Ac127496.1, Ac132872.2, Ac135178.1, Ac140061.12, Ac140481.2, Acaa2, Acaca, Acacb, Acad11, Acad8, Acan, Acat1, Acbd7, Acer3, Ackr2, Ackr3, Ackr4, Acn9, Aco2, Acot13, Acot2, Acot9, Acox1, Acox3, Acp6, Acrv1, Acsf2, Acsl3, Acsl6, Acsm2a, Acsm3, Acsm4, Acss1, Acss3, Actb, Actbl2, Actg1, Actl8, Actn2, Actn4, Actr10, Actr2, Actr3, Actr3b, Actr6, Actr8, Acvr1c, Acvr2a, Acvrl1, Acyp1, Adal, Adam17, Adam22, Adam23, Adam28, Adam7, Adamts1, Adamts15, Adamts19, Adamts3, Adamts4, Adamts5, Adamts9, Adamtsl1, Adamtsl5, Adap1, Adar, Adarb2, Adat1, Adcy3, Adcy5, Adcy8, Adcyap1r1, Add2, Adh1b, Adh5, Adh6, Adh7, Adhfe1, Adı1, Adıpoq, Adk, Adm, Ado, Adprhl1, Adra1a, Adra2a, Adrbk1, Adrbk2, Adtrp, Aen, Afap1, Afap1l2, Aff2, Aff3, Aff4, Afmıd, Agap1, Agap9, Agbl3, Agbl4, Agbl5, Agfg1, Aggf1, Agmat, Ago4, Agpat3, Agpat4, Agpat6, Agpat9, Agr2, Agrn, Agt, Agtpbp1, Agtr2, Agtrap, Agxt, Agxt2, Ahctf1, Ahcy, Ahcyl2, Ahdc1, Ahı1, Ahnak, Ahr, Ahrr, Aıf1l, Aıg1, Aım1, Aımp1, Aıpl1, Ajuba, Ak1, Ak4, Ak7, Ak9, Akap11, Akap12, Akap2, Akap5, Akap6, Akap7, Akap8, Akıp1, Akırın2, Akna, Akr1c1, Akr1c2, Akr1c4, Akr7a2, Akt1s1, Akt2, Al031663.2, Al079342.1, Al117190.3, Al133373.1, Al161915.1, Al162407.1, Al163636.6, Al353698.1, Al353791.1, Al355390.1, Al359878.1, Al391152.1, Al450307.1, Al583828.1, Al589765.1, Al591684.1, Al603965.1, Al645728.1, Al953854.2, Alad, Alcam, Aldh16a1, Aldh18a1, Aldh1a3, Aldh1b1, Aldh3a2, Aldh4a1, Aldh5a1, Aldh6a1, Aldh7a1, Aldh8a1, Aldoa, Alg1, Alg10, Alg10b, Alg6, Alkbh1, Alkbh4, Alkbh5, Alkbh8, Alox15, Als2, Alx1, Alx4, Amd1, Amdhd1, Amer1, Amer2, Amer3, Ammecr1, Amn1, Amot, Amotl1, Amotl2, Ampd3, Anapc16, Ang, Angel2, Angpt1, Angpt2, Angpt4, Ank2, Ankdd1a, Ankdd1b, Ankef1, Ankfn1, Ankh, Ankle2, Ankmy1, Ankmy2, Ankrd10, Ankrd11, Ankrd13a, Ankrd13b, Ankrd17, Ankrd22, Ankrd26, Ankrd28, Ankrd30b, Ankrd31, Ankrd32, Ankrd33b, Ankrd34b, Ankrd34c, Ankrd42, Ankrd44, Ankrd45, Ankrd46, Ankrd49, Ankrd50, Ankrd55, Ankrd6, Ankrd63, Ankrd9, Anks3, Anks4b, Anks6, Anln, Ano10, Ano4, Anp32a, Anp32b, Anp32e, Antxr2, Anxa11, Anxa4, Anxa5, Anxa6, Ap000695.1, Ap000708.1, Ap001652.1, Ap003068.23, Ap003774.4, Ap005482.1, Ap1ar, Ap1s2, Ap1s3, Ap3b1, Ap3b2, Ap3m1, Ap3s2, Ap4e1, Ap4s1, Ap5b1, Ap5m1, Ap5s1, Apaf1, Apba1, Apba2, Apbb2, Apex1, Aph1a, Aph1b, Apıtd1-Cort, Aplf, Apln, Apoa1, Apob, Apobec3c, Apobec3d, Apobec3f, Apoc3, Apol2, Apol6, Apool, App, Appbp2, Aptx, Aqp11, Aqp4, Aqp6, Aqr, Arap1, Arcn1, Arel1, Arf3, Arf4, Arf6, Arfgap1, Arfgap2, Arg1, Arg2, Argfx, Arhgap15, Arhgap18, Arhgap19, Arhgap21, Arhgap22, Arhgap24, Arhgap25, Arhgap26, Arhgap28, Arhgap29, Arhgap31, Arhgap35, Arhgap42, Arhgap6, Arhgef10, Arhgef11, Arhgef2, Arhgef28, Arhgef3, Arhgef33, Arhgef37, Arhgef38, Arhgef39, Arhgef5, Arhgef7, Arıd3a, Arıd4a, Arl1, Arl10, Arl13b, Arl14ep, Arl14epl, Arl15, Arl2bp, Arl3, Arl4a, Arl4c, Arl5a, Arl5b, Arl5c, Arl6, Arl6ıp1, Arl6ıp5, Arl6ıp6, Arl8b, Armc1, Armc10, Armc3, Armcx3, Armcx5, Arnt, Arpc1a, Arpc2, Arpc5, Arpc5l, Arrb1, Arrb2, Arrdc2, Arsa, Arsb, Arsd, Arse, Arsk, As3mt, Asah2, Asah2b, Asap1, Asb1, Asb14, Asb15, Asb16, Asb18, Asb4, Asb5, Asb6, Ascc1, Ascc2, Asf1a, Ash2l, Aspa, Asph, Aspn, Astn1, Astn2, Asxl3, Asz1, Atad2, Atad2b, Atad3c, Atad5, Ate1, Atf1, Atf5, Atf6, Atf7, Atf7ıp, Atf7ıp2, Atg10, Atg12, Atg2a, Atg2b, Atg3, Atg4c, Atg5, Atıc, Atl2, Atm, Atmın, Atoh7, Atox1, Atp11b, Atp11c, Atp13a3, Atp13a4, Atp1b1, Atp1b2, Atp1b3, Atp1b4, Atp2a2, Atp2b1, Atp2b4, Atp5a1, Atp5b, Atp5c1, Atp5e, Atp5f1, Atp5g1, Atp5g3, Atp5j2-Ptcd1, Atp5s, Atp6ap1l, Atp6v0b, Atp6v0d2, Atp6v0e1, Atp6v1a, Atp6v1c1, Atp6v1c2, Atp6v1d, Atp6v1e1, Atp6v1e2, Atp6v1f, Atp6v1g1, Atp7a, Atp7b, Atp8a1, Atp8b2, Atp8b3, Atp8b4, Atp9a, Atp9b, Atpaf2, Atrnl1, Atxn3, Atxn7l3b, Aunıp, Auts2, Avl9, Avpr1a, Axın1, Axın2, Axl, Azı2, B2m, B3galt1, B3galt2, B3galtl, B3gat2, B3gat3, B3gnt2, B3gnt4, B3gnt5, B3gnt7, B3gnt8, B4galnt2, B4galt1, B4galt4, B4galt7, Baat, Bace2, Bach1, Bach2, Bag1, Bag2, Bag4, Banf1, Bard1, Barx2, Baz1a, Baz2a, Baz2b, Bbc3, Bbıp1, Bbs5, Bbs7, Bbx, Bcar3, Bcas1, Bcas2, Bcas4, Bcat1, Bcdın3d, Bckdhb, Bckdk, Bcl10, Bcl2, Bcl2l13, Bcl2l14, Bcl2l15, Bcl2l2-Pabpn1, Bcl6, Bcl7b, Bcl9, Bclaf1, Bcr, Bdh2, Begaın, Bend2, Bend7, Best1, Best3, Bet1, Bex1, Bex2, Bhlhb9, Bhlhe22, Bhlhe40, Bhmt, Bıd, Bık, Bın1, Bın3, Bırc3, Bırc5, Bırc6, Bıvm, Bloc1s2, Bloc1s3, Bloc1s4, Bloc1s5, Bloc1s6, Blvra, Blzf1, Bmf, Bmı1, Bmp10, Bmp2, Bmp5, Bmp6, Bmp7, Bmp8b, Bmper, Bmpr1a, Bmpr1b, Bms1, Bnc1, Bnıp3l, Bnıpl, Bod1, Bpgm, Bpnt1, Brat1, Brcc3, Brd3, Brd4, Brd7, Brf2, Brı3, Brı3bp, Brıp1, Brıx1, Brms1l, Brox, Bsdc1, Bst2, Btbd10, Btbd19, Btbd3, Btd, Btg2, Btla, Btn1a1, Btnl9, Btrc, Bub1, Bub3, Bves, Bx255923.1, Bysl, Bzw1, Bzw2, C10orf10, C10orf113, C10orf118, C10orf137, C10orf55, C10orf67, C10orf76, C10orf82, C11orf1, C11orf34, C11orf45, C11orf48, C11orf54, C11orf58, C11orf73, C11orf74, C11orf82, C11orf83, C11orf84, C11orf87, C12orf23, C12orf4, C12orf40, C12orf42, C12orf49, C12orf5, C12orf52, C12orf60, C12orf61, C12orf65, C12orf74, C12orf75, C12orf76, C13orf35, C14orf1, C14orf119, C14orf132, C14orf142, C14orf2, C14orf23, C14orf28, C14orf37, C14orf39, C15orf26, C15orf37, C15orf37, C15orf38, C15orf38-Ap3s2, C15orf40, C15orf48, C15orf52, C16orf45, C16orf46, C16orf54, C16orf71, C16orf72, C16orf74, C17orf102, C17orf103,* |

C17orf104, C17orf105, C17orf75, C17orf77, C17orf80, C17orf85, C18orf21, C18orf25, C18orf32, C18orf42, C18orf54, C18orf63, C19orf10, C19orf12, C19orf35, C19orf38, C19orf48, C19orf52, C19orf54, C19orf55, C19orf66, C1galt1, C1galt1c1, C1orf101, C1orf106, C1orf109, C1orf122, C1orf143, C1orf145, C1orf158, C1orf173, C1orf180, C1orf185, C1orf192, C1orf198, C1orf21, C1orf210, C1orf213, C1orf229, C1orf35, C1orf50, C1orf95, C1ql3, C1qtnf2, C1qtnf3, C1qtnf7, C20orf197, C20orf26, C20orf85, C21orf58, C21orf62, C21orf90, C22orf29, C2cd2l, C2orf15, C2orf15, C2orf27a, C2orf27b, C2orf43, C2orf48, C2orf49, C2orf50, C2orf54, C2orf66, C2orf68, C2orf69, C2orf72, C2orf76, C2orf80, C2orf83, C2orf91, C3, C3orf14, C3orf17, C3orf18, C3orf33, C3orf36, C3orf38, C3orf58, C3orf62, C4orf17, C4orf26, C4orf29, C4orf3, C4orf32, C4orf33, C4orf36, C4orf40, C4orf46, C4orf50, C5, C5ar2, C5orf15, C5orf22, C5orf28, C5orf42, C5orf45, C5orf47, C5orf49, C5orf51, C5orf64, C6, C6orf132, C6orf141, C6orf195, C6orf211, C6orf25, C6orf57, C6orf58, C6orf62, C6orf89, C7, C7orf13, C7orf25, C7orf41, C7orf43, C7orf55-Luc7l2, C7orf57, C7orf63, C7orf73, C8a, C8orf17, C8orf33, C8orf37, C8orf44-Sgk3, C8orf46, C8orf47, C8orf59, C9, C9orf156, C9orf170, C9orf3, C9orf40, C9orf41, C9orf47, C9orf64, C9orf66, C9orf72, C9orf78, C9orf84, C9orf91, Ca1, Ca12, Ca13, Ca14, Ca3, Ca5b, Ca6, Ca8, Caap1, Cabin1, Cables2, Cabyr, Cacna1b, Cacna1d, Cacna1e, Cacna1i, Cacna2d1, Cacna2d2, Cacnb4, Cacng2, Cacng4, Cacng8, Cacybp, Cad, Cadm1, Cadps2, Calcb, Calcoco1, Calcoco2, Calhm2, Calm2, Caln1, Camk2a, Camk2n1, Camk4, Camkk1, Camkk2, Camkmt, Camkv, Camlg, Canx, Cap1, Cap2, Capn14, Capns1, Capza1, Capza2, Card8, Carf, Carhsp1, Casc10, Casc3, Casc4, Casc5, Casd1, Cask, Casp1, Casp10, Casp16, Casp2, Casp6, Casp8, Casp9, Casq2, Cass4, Catsper2, Cbll1, Cbr4, Cbs, Cbwd1, Cbx3, Cbx4, Cbx5, Cby1, Cby3, Cc2d2b, Ccar1, Ccar2, Ccbe1, Ccdc103, Ccdc107, Ccdc109b, Ccdc113, Ccdc114, Ccdc115, Ccdc117, Ccdc121, Ccdc122, Ccdc125, Ccdc127, Ccdc129, Ccdc132, Ccdc138, Ccdc14, Ccdc141, Ccdc147, Ccdc15, Ccdc150, Ccdc153, Ccdc158, Ccdc163p, Ccdc170, Ccdc171, Ccdc18, Ccdc23, Ccdc25, Ccdc36, Ccdc38, Ccdc41, Ccdc43, Ccdc47, Ccdc65, Ccdc67, Ccdc68, Ccdc77, Ccdc8, Ccdc80, Ccdc85c, Ccdc89, Ccdc90b, Ccdc92, Ccdc93, Ccl11, Ccl16, Ccl22, Ccl24, Ccl28, Ccl3l1, Ccl3l3, Ccl8, Ccnb1, Ccnb1ip1, Ccndbp1, Ccne1, Ccnf, Ccng1, Ccng2, Ccnh, Ccni2, Ccnj, Ccnjl, Ccnk, Ccnl2, Ccno, Ccnyl1, Ccp110, Ccpg1, Ccr1, Ccr10, Ccr2, Ccr4, Ccs, Ccsap, Ccser1, Cct5, Cd109, Cd163, Cd180, Cd1c, Cd1e, Cd200, Cd200r1, Cd209, Cd226, Cd28, Cd2ap, Cd300ld, Cd302, Cd36, Cd3d, Cd3eap, Cd3g, Cd40lg, Cd44, Cd46, Cd47, Cd5, Cd55, Cd58, Cd59, Cd86, Cd9, Cd93, Cdc14a, Cdc14b, Cdc20b, Cdc23, Cdc37l1, Cdc40, Cdc42, Cdc42bpb, Cdc42ep4, Cdc42se2, Cdc5l, Cdc7, Cdca2, Cdca7l, Cdcp1, Cdh17, Cdh18, Cdh19, Cdh26, Cdh4, Cdh6, Cdh7, Cdh8, Cdhr3, Cdhr4, Cdip1, Cdipt, Cdk1, Cdk12, Cdk16, Cdk2, Cdk2ap1, Cdk3, Cdk5r2, Cdk8, Cdk9, Cdkal1, Cdkl1, Cdkl2, Cdkl4, Cdkn1b, Cdkn2aip, Cdkn2aipnl, Cdkn2b, Cdkn2c, Cdnf, Cdon, Cdr2l, Cds2, Cdv3, Cdx1, Cdx2, Ceacam1, Ceacam8, Cebpb, Cebpg, Cebpz, Cecr5, Celf2, Cenpa, Cenpc, Cenpf, Cenph, Cenpi, Cenpj, Cenpl, Cenpm, Cenpn, Cep104, Cep120, Cep135, Cep164, Cep19, Cep41, Cep44, Cep57, Cep57l1, Cep68, Cep70, Cep72, Cep76, Cep78, Cep85l, Cep89, Cep97, Cercam, Cerk, Cers4, Cers5, Cers6, Ces4a, Cfd, Cfhr4, Cggbp1, Cgnl1, Cgref1, Cgrrf1, Chaf1b, Chchd3, Chchd5, Chchd7, Chd1, Chd1l, Chd6, Chd7, Chdh, Chek1, Cherp, Chfr, Chic2, Chka, Chml, Chmp1b, Chmp2b, Chmp3, Chmp5, Chp1, Chrdl1, Chrfam7a, Chrm2, Chrm3, Chrna4, Chrna7, Chrng, Chst11, Chst14, Chst3, Chst6, Chsy1, Chtop, Ciao1, Ciita, Cinp, Cirbp, Cirh1a, Cited2, Ckap2, Ckap4, Ckb, Cks2, Clasp1, Clasp2, Clca1, Clca2, Clcc1, Clcn3, Clcn4, Clcn5, Cldn1, Cldn12, Cldn19, Cldn2, Clec12a, Clec12b, Clec14a, Clec17a, Clec1a, Clec3a, Clec4c, Clec5a, Clec7a, Clecl1, Clic4, Clic5, Clint1, Clk1, Clk4, Cllu1, Clmn, Clmp, Cln8, Clnk, Clns1a, Clpb, Clpx, Clrn1, Clspn, Clstn1, Clstn2, Clu, Cluap1, Cluh, Clul1, Clvs1, Clvs2, Cmas, Cmbl, Cmpk1, Cmss1, Cmtm4, Cmtm8, Cmtr2, Cnbp, Cnep1r1, Cnih3, Cnih4, Cnksr3, Cnn3, Cnnm3, Cnnm4, Cnot1, Cnot11, Cnot2, Cnot3, Cnot4, Cnot8, Cnpy1, Cnr2, Cnrip1, Cntd1, Cntf, Cntln, Cntn4, Cntn6, Cntnap1, Cntnap3, Cntnap3b, Cntrl, Coa4, Coa5, Cobll1, Coch, Cog3, Cog6, Coil, Col11a1, Col12a1, Col13a1, Col14a1, Col16a1, Col1a1, Col20a1, Col24a1, Col25a1, Col27a1, Col3a1, Col4a1, Col4a3bp, Col5a1, Col6a3, Col6a5, Col8a1, Col8a2, Col9a2, Colca1, Colec10, Colec12, Colgalt2, Commd1, Commd10, Commd3-Bmi1, Commd6, Commd9, Copa, Cops2, Cops3, Cops7a, Cops7b, Cops8, Copz1, Coq4, Coq7, Coro1c, Coro2a, Coro2b, Coro7, Cort, Cotl1, Cox16, Cox17, Cox18, Cox19, Cox20, Cox6b1, Cox6b2, Cox7a2l, Cox7b, Cox7c, Cp, Cpa3, Cpa4, Cpe, Cpeb2, Cped1, Cpm, Cpne1, Cpne3, Cpne4, Cps1, Cpsf1, Cpsf4, Cpsf6, Cpt1a, Cpxm2, Cr1, Cr769776.1, Cradd, Cramp1l, Crb1, Crbn, Creb1, Creb3l2, Creb5, Creg1, Creg2, Creld1, Crim1, Cript, Crispld1, Crispld2, Crkl, Crlf3, Crls1, Crnkl1, Crocc, Crot, Crtam, Crtap, Crx, Cry1, Crybg3, Cryl1, Cryz, Csde1, Csf1, Csgalnact2, Csk, Csmd2, Csmd3, Csn1s1, Csn2, Csnk1a1l, Csnk1g3, Csnk2b, Csnk2b-Ly6g5b-1181, Cspg5, Csrnp1, Csrp1, Csrp3, Csta, Cstf2, Cstf2t, Ctage8, Ctage9, Ctb-167g5.5, Ctb-186h2.3, Ctbp1, Ctc1, Ctc-432m15.3, Ctcf, Ctd-2162k18.4, Ctd-2267d19.3, Ctd-2368p22.1, Ctd-2545m3.6, Ctdspl, Ctf1, Cth, Ctla4, Ctnna1, Ctnna2, Ctnna3, Ctnnal1, Ctnnbl1, Ctns, Ctrc, Ctsb, Ctse, Ctsk, Ctsl, Ctss, Cttn, Cttnbp2, Cttnbp2nl, Cul3, Cul4a, Cul4b, Cuta, Cux2, Cwc22, Cwc25, Cwc27, Cwf19l1, Cwf19l2, Cx3cl1, Cxcl1, Cxcl13, Cxcl14, Cxcl16, Cxcl3, Cxcr2, Cxcr4, Cxcr5, Cxorf21, Cxorf22, Cxorf24, Cxorf30, Cxorf38, Cxorf40a, Cxorf56, Cxxc4, Cxxc5, Cyb561a3, Cyb5a, Cyb5b, Cyb5d1, Cyb5r1, Cyb5r3, Cyb5r4, Cybb, Cybrd1, Cycs, Cyfip1, Cyfip2, Cyp1a2, Cyp1b1, Cyp20a1, Cyp26a1, Cyp26b1, Cyp27a1, Cyp27b1, Cyp27c1, Cyp2a7, Cyp2b6, Cyp2c19, Cyp2u1, Cyp3a4, Cyp3a5, Cyp4f12, Cyp4f3, Cyp4v2, Cyp51a1, Cyp7b1, Cyp8b1, Cystm1, Cyth2, Cyth3, Daam1,

*Dab1, Dab2, Dab2ip, Dach1, Dagla, Dak, Dand5, Dapk1, Dars, Dars2, Dazap1, Dazap2, Dazl, Dbf4, Dbnl, Dbr1, Dbt, Dbx2, Dcaf10, Dcaf12l1, Dcaf12l2, Dcaf13, Dcaf16, Dcaf17, Dcaf4l1, Dcaf4l2, Dcdc1, Dcdc2, Dchs2, Dck, Dclk1, Dcps, Dctn3, Dctn4, Dctn5, Dcun1d2, Dcun1d4, Dcx, Ddah1, Ddhd2, Ddit4l, Ddo, Ddr1, Ddr2, Ddrgk1, Ddx10, Ddx18, Ddx19a, Ddx19b, Ddx20, Ddx25, Ddx46, Ddx5, Ddx53, Ddx60, Decr1, Dedd, Defb105a, Defb105b, Defb118, Defb132, Dennd1b, Dennd2c, Dennd2d, Dennd5a, Dennd5b, Dennd6b, Denr, Depdc4, Depdc5, Depdc7, Derl1, Desi1, Dffa, Dffb, Dfnb59, Dgat2, Dgcr14, Dgkb, Dgkd, Dgkh, Dhcr24, Dhdds, Dhfr, Dhfrl1, Dhodh, Dhtkd1, Dhx35, Dhx36, Dhx37, Dhx58, Diaph3, Diexf, Dimt1, Dip2b, Diras1, Disc1, Dixdc1, Dkfzp434e1119, Dkfzp434o1614, Dkfzp779l1853, Dkk1, Dleu1, Dlg1, Dlg2, Dlg3, Dlg4, Dlgap1, Dlgap2, Dlgap4, Dlk1, Dll1, Dll4, Dlx2, Dlx4, Dlx5, Dmd, Dmgdh, Dmkn, Dmp1, Dmrt2, Dmrta1, Dmrtc2, Dmtf1, Dmxl1, Dna2, Dnaaf2, Dnah17-As1, Dnah5, Dnah7, Dnah9, Dnajb11, Dnajb4, Dnajb5, Dnajb6, Dnajb7, Dnajb9, Dnajc10, Dnajc13, Dnajc14, Dnajc15, Dnajc18, Dnajc21, Dnajc22, Dnajc24, Dnajc25, Dnajc25-Gng10, Dnajc27, Dnajc28, Dnajc3, Dnajc30, Dnajc5, Dnajc6, Dnajc9, Dnase1l2, Dnase1l3, Dnm3, Dnmbp, Dnmt3b, Dnpep, Dnttip2, Dock1, Dock11, Dock7, Dock9, Dok4, Dok5, Dok6, Dolpp1, Dopey1, Dpagt1, Dpf2, Dpf3, Dph3, Dph7, Dpm1, Dpp8, Dppa3, Dppa4, Dpt, Dpy19l3, Dpy19l4, Dpyd, Dpysl2, Dpysl3, Dram2, Draxin, Drp2, Dsc2, Dsc3, Dscr3, Dse, Dsg2, Dsp, Dstn, Dstyk, Dtd2, Dtl, Dtwd2, Dtx3l, Dtx4, Dusp1, Dusp10, Dusp16, Dusp18, Dusp19, Dusp2, Dusp22, Dusp28, Dusp3, Dusp4, Dusp5, Dusp6, Duxa, Dym, Dync1i2, Dync1li1, Dync2h1, Dyrk4, Dysf, Dytn, Dyx1c1, Dzank1, Dzip1, Dzip1l, E2f2, E2f6, E2f7, Eaf1, Eapp, Ebf4, Ebna1bp2, Ebpl, Ece1, Echdc2, Echdc3, Echs1, Ect2l, Eda2r, Edc4, Edem1, Edem2, Edem3, Edil3, Ednra, Eed, Eef1a1, Eef1e1, Eef2k, Efcab1, Efcab11, Efcab12, Efcab13, Efcab14, Efcab2, Efcab5, Efcab6, Efhc1, Efhc2, Efr3b, Egfl6, Egfl8, Egfr, Egln1, Egr1, Ehd1, Ehd2, Ehd3, Ehmt1, Eid2, Eid2b, Eif1, Eif1ad, Eif1ax, Eif1b, Eif2a, Eif2ak1, Eif2ak2, Eif2ak3, Eif2b2, Eif2s1, Eif2s2, Eif2s3, Eif2s3l, Eif3a, Eif3f, Eif3h, Eif3l, Eif3m, Eif4a1, Eif4a3, Eif4ebp1, Eif4ebp2, Eif4enif1, Eif4g3, Eif5a2, Eif5b, Elac1, Elac2, Elavl1, Elavl3, Elf1, Elf2, Elf4, Elf5, Elk1, Elk4, Ell3, Elmo1, Elmo2, Eln, Elovl2, Elovl4, Elovl6, Elp2, Elp3, Elp4, Elp6, Emb, Emc1, Emc10, Emc2, Emc3, Emcn, Emilin2, Eml1, Eml5, Emp1, Emp2, Emr2, En1, En2, Enam, Endod1, Endou, Eno4, Enosf1, Enpp1, Enpp3, Enpp4, Enpp6, Enthd1, Entpd4, Entpd5, Eny2, Eomes, Ep400, Epas1, Epb41l1, Epb41l4a, Epb41l4b, Epb42, Epha10, Epha4, Epha6, Epha7, Ephb4, Ephx2, Epm2a, Epm2aip1, Epor, Eppk1, Eps15l1, Eps8, Epsti1, Ept1, Erap1, Erap2, Erbb2, Erbb2ip, Ercc1, Ercc2, Ercc4, Ercc6l, Ercc6l2, Ercc8, Ergic1, Ergic2, Eri1, Eri2, Erlin2, Ermap, Ermn, Ern1, Ero1lb, Errfi1, Ervmer34-1, Esco1, Esco2, Esd, Esf1, Esm1, Esr2, Esyt1, Esyt2, Esyt3, Etfa, Ets2, Etv6, Etv7, Evc, Evl, Evpll, Evx1, Exd1, Exo1, Exoc1, Exoc3, Exoc6, Exoc8, Exosc1, Exosc2, Exosc3, Exosc6, Exosc9, Exph5, Ext2, Extl2, Extl3, Eya2, Eya3, Eys, Ezr, F11, F11r, F2, F2rl1, F2rl2, Fabp2, Fabp7, Fads1, Fads2, Fads6, Faf1, Fahd1, Fam101b, Fam103a1, Fam105a, Fam105b, Fam106a, Fam107b, Fam110a, Fam110c, Fam111a, Fam114a2, Fam115c, Fam117a, Fam118a, Fam120b, Fam120c, Fam122a, Fam122b, Fam124a, Fam124b, Fam126a, Fam127a, Fam127c, Fam129a, Fam133a, Fam133b, Fam134a, Fam135a, Fam13a, Fam13b, Fam13c, Fam149b1, Fam150a, Fam151b, Fam156a, Fam156b, Fam159b, Fam160b1, Fam161a, Fam162a, Fam162b, Fam169a, Fam171a1, Fam171b, Fam173b, Fam174b, Fam175b, Fam177a1, Fam178a, Fam179a, Fam180a, Fam180b, Fam181b, Fam182b, Fam184b, Fam186a, Fam189a1, Fam192a, Fam193a, Fam193b, Fam195b, Fam196a, Fam198a, Fam198b, Fam199x, Fam19a1, Fam19a4, Fam19a5, Fam207a, Fam208a, Fam20a, Fam210a, Fam210b, Fam211a, Fam214a, Fam216a, Fam216b, Fam219b, Fam220a, Fam221a, Fam227a, Fam227b, Fam228a, Fam228b, Fam229b, Fam26e, Fam26f, Fam3a, Fam3c, Fam45a, Fam46d, Fam49b, Fam53c, Fam65b, Fam65c, Fam71f2, Fam72a, Fam72b, Fam72c, Fam72d, Fam81a, Fam83h, Fam84a, Fam84b, Fam8a1, Fam91a1, Fam92a1, Fam96a, Fam98a, Fam98b, Fam9c, Fanca, Fancb, Fancc, Fancd2, Fance, Fancf, Fancl, Fancm, Far1, Far2, Farsb, Fastkd1, Fastkd2, Fat1, Fat2, Fat4, Fblim1, Fbln1, Fbln2, Fbn1, Fbn2, Fbrsl1, Fbxl13, Fbxl17, Fbxl18, Fbxl20, Fbxl5, Fbxo11, Fbxo15, Fbxo21, Fbxo22, Fbxo25, Fbxo3, Fbxo30, Fbxo31, Fbxo38, Fbxo39, Fbxo4, Fbxo41, Fbxo44, Fbxo45, Fbxo46, Fbxo47, Fbxw2, Fbxw8, Fcar, Fcf1, Fcgr2a, Fcho2, Fchsd2, Fcrl1, Fcrl2, Fcrl5, Fcrlb, Fdft1, Fdx1, Fdxacb1, Fech, Fem1a, Fem1b, Fen1, Fermt1, Fetub, Fez2, Ffar2, Fga, Fgb, Fgd1, Fgd2, Fgd4, Fgd5, Fgd6, Fgf1, Fgf12, Fgf14, Fgf23, Fgf5, Fgfbp1, Fgfbp3, Fgfr1, Fgfr1op, Fgfr2, Fgfr3, Fgfr4, Fgg, Fgl2, Fhdc1, Fhl2, Fig4, Figf, Fignl1, Fip1l1, Fitm2, Fkbp14, Fkbp1a, Fkbp1c, Fkbp7, Fkbp9, Fktn, Flcn, Flg, Flj00104, Flj20373, Flrt2, Flrt3, Flt1, Flvcr1, Flvcr2, Flywch2, Fmn1, Fmnl3, Fmo2, Fmo3, Fn3krp, Fndc1, Focad, Fopnl, Fos, Fosb, Foxa1, Foxa2, Foxa3, Foxb1, Foxc1, Foxc2, Foxd1, Foxd2, Foxf1, Foxi2, Foxj2, Foxj3, Foxk1, Foxk2, Foxl1, Foxl2, Foxn2, Foxn3, Foxo1, Foxo3, Foxp4, Foxq1, Foxr2, Foxred2, Fpgt, Fpr1, Fpr2, Fra10ac1, Frat1, Frem2, Frg1b, Frg2, Frg2b, Frg2c, Frmd3, Frmd4a, Frmd4b, Frmd5, Frmd7, Frmd8, Frmpd4, Frrs1, Frrs1l, Frs3, Fryl, Frzb, Fsbp, Fscn1, Fsd1l, Fshb, Fsip1, Fsip2, Fst, Fstl1, Fstl4, Fstl5, Ftl, Fubp1, Fuca2, Fundc2, Fus, Fut1, Fut10, Fut11, Fut2, Fut8, Fxn, Fyco1, Fyttd1, Fzd1, Fzd2, Fzd5, G3bp1, G6pc, G6pc2, Gab3, Gab4, Gabarap, Gabarapl3, Gabbr1, Gabpa, Gabpb1, Gabpb2, Gabra2, Gabra5, Gabrb1, Gabrb3, Gabrg1, Gabrg3, Gabrp, Gabrq, Gad1, Gadd45a, Gadd45g, Gadd45gip1, Gadl1, Gage1, Gal, Gal3st4, Galc, Galk2, Galnt10, Galnt13, Galnt2, Galnt3, Galnt5, Galnt7, Galntl6, Galr1, Galt, Gap43, Gapt, Gart, Gas1, Gas2, Gas2l3, Gas7, Gata3, Gatad2a, Gatm, Gbp1, Gbp3, Gbp4, Gbp6, Gbx2, Gca, Gcc2, Gcfc2, Gch1, Gclc, Gclm, Gcm1, Gcn1l1, Gcnt1, Gcnt3, Gcnt4, Gcsaml, Gdap1, Gdap2, Gde1, Gdf11,*

Gdf5os, Gdf6, Gdf7, Gdı2, Gdnf, Gdpd1, Gdpgp1, Gem, Gemın2, Gemın6, Gen1, Get4, Gfi1, Gfm1, Gfod1,
Gfpt2, Gfra1, Gfra2, Gfral, Gga2, Ggact, Ggcx, Ggps1, Ghıtm, Gıd8, Gıgyf2, Gımap1, Gımap4, Gınm1,
Gıns1, Gıns3, Gıns4, Gıpc2, Gıpr, Gja3, Gjb3, Gjc1, Gjd2, Gk, Gk5, Glb1l3, Glce, Gldc, Gldn, Gle1, Glı3,
Glı4, Glıpr1l2, Glıpr2, Glmn, Glod4, Glp1r, Glp2r, Glrx2, Glrx3, Gls, Gltscr1, Gltscr1l, Gltscr2, Glud1,
Glud2, Glul, Glyat, Glyatl1, Glyatl3, Glyctk, Gm2a, Gmeb1, Gmnc, Gmpr, Gmps, Gna11, Gna12, Gna14,
Gnaı3, Gnal, Gnao1, Gnat1, Gnaz, Gnb1, Gnb4, Gnb5, Gne, Gng10, Gng2, Gng4, Gng7, Gnl3l, Gnpda2,
Gnpnat1, Gnptab, Gnrhr, Gns, Golga1, Golga3, Golga6l1, Golga6l10, Golga6l4, Golga6l6, Golga6l9,
Golım4, Golm1, Golph3, Golph3l, Golt1b, Gon4l, Gopc, Gorab, Gorasp2, Gosr2, Got1, Gp5, Gpalpp1,
Gpank1, Gpatch1, Gpatch11, Gpatch2, Gpatch2l, Gpatch8, Gpbp1l1, Gpc3, Gpc4, Gpc5, Gpd1l, Gpd2, Gpı,
Gpkow, Gpld1, Gpm6a, Gpm6b, Gpn2, Gpn3, Gpr1, Gpr110, Gpr111, Gpr113, Gpr116, Gpr125, Gpr132,
Gpr133, Gpr135, Gpr137c, Gpr141, Gpr157, Gpr160, Gpr17, Gpr174, Gpr179, Gpr25, Gpr27, Gpr34,
Gpr37, Gpr37l1, Gpr39, Gpr4, Gpr50, Gpr56, Gpr64, Gpr65, Gpr78, Gpr82, Gpr83, Gpr85, Gpr88,
Gprasp1, Gprasp2, Gprc5a, Gprc5b, Gprın2, Gpx5, Gpx7, Gpx8, Gramd1c, Gramd2, Gramd4, Grap2,
Grasp, Grb7, Greb1l, Grem1, Grem2, Grhl1, Grıa1, Grıa2, Grıd2, Grık1-As2, Grık2, Grık3, Grık4, Grık5,
Grın2b, Grın3a, Grk4, Grk5, Grm3, Grm6, Grpel2, Grwd1, Gsdma, Gsdmb, Gse1, Gsg2, Gspt1, Gsr, Gsta4,
Gstcd, Gstk1, Gstm2, Gstm5, Gsto1, Gsx2, Gtf2b, Gtf2f1, Gtf2f2, Gtf2h5, Gtf3c1, Gtf3c3, Gtf3c4, Gtf3c6,
Gtpbp1, Gtpbp10, Gtpbp3, Gtpbp4, Gtpbp8, Guca1b, Gucd1, Gucy1a2, Gucy1a3, Gucy2c, Guf1, Gvqw1,
Gxylt2, Gyg2, Gzf1, H1fx, H2afj, H2afv, H2afy, H2afy2, H3f3c, H6pd, Haao, Habp2, Habp4, Hace1, Hadh,
Hapln1, Hapln3, Harbı1, Has2, Has3, Hat1, Haus2, Haus3, Haus5, Havcr2, Hbs1l, Hcar1, Hccs, Hcfc1, Hck,
Hcn1, Hdac1, Hdac2, Hdac3, Hdac9, Hdhd1, Hdhd2, Hdx, Heatr3, Heatr5a, Heatr6, Hebp2, Heca, Hectd1,
Hectd2, Hells, Helz, Hemk1, Henmt1, Hepacam, Hepacam2, Heph, Hephl1, Herc3, Herc4, Herpud2, Hes1,
Hexa, Hexb, Heyl, Hfe, Hgsnat, Hhat, Hhıpl1, Hıat1, Hıatl1, Hıatl2, Hıbadh, Hıbch, Hıf1an, Hıf3a, Hıgd1a,
Hınfp, Hınt1, Hınt3, Hıst1h1c, Hıst1h2ab, Hıst1h2ag, Hıst1h2ah, Hıst1h2aı, Hıst1h2ak, Hıst1h2bd,
Hıst1h2bf, Hıst1h2bj, Hıst1h2bn, Hıst1h2bo, Hıst1h3b, Hıst1h3e, Hıst1h3g, Hıst1h3h, Hıst1h4e, Hıst2h2be,
Hıst2h4a, Hıst2h4b, Hıst3h2bb, Hıvep1, Hıvep2, Hıvep3, Hjurp, Hla-A, Hla-Doa, Hla-Dob, Hla-Dqb1, Hltf,
Hlx, Hmbox1, Hmces, Hmcn1, Hmg20a, Hmga1, Hmgb1, Hmgb3, Hmgcl, Hmgcll1, Hmgn1, Hmgn2, Hmgn4,
Hmha1, Hmox1, Hmsd, Hmx3, Hnf4a, Hnf4g, Hnmt, Hnrnpa0, Hnrnpa1l2, Hnrnpa3, Hnrnpdl, Hnrnph1,
Hnrnpk, Hnrnpl, Hnrnpll, Hnrnpm, Hnrnpr, Hoga1, Homer3, Homez, Hook1, Hormad1, Hormad2, Hoxa1,
Hoxa10, Hoxa13, Hoxa2, Hoxa3, Hoxb3, Hoxb4, Hoxb7, Hoxb9, Hoxc11, Hoxc13, Hoxc8, Hoxd11, Hoxd3,
Hpgd, Hps3, Hps4, Hpse, Hpse2, Hrasls5, Hrh1, Hrh2, Hrh4, Hrnr, Hs1bp3, Hs3st1, Hs3st3a1, Hs3st3b1,
Hs6st1, Hs6st3, Hsd11b1, Hsd17b11, Hsd17b12, Hsd17b13, Hsd17b3, Hsd17b6, Hsd17b8, Hsd3b7, Hsdl1,
Hsdl2, Hsf2, Hsf2bp, Hsf5, Hspa12a, Hspa13, Hspa14, Hspa1a, Hspa1b, Hspa6, Hspb3, Hspd1, Hspe1-Mob4,
Hsph1, Htr1d, Htr1f, Htr2c, Htr5a, Htr5a-As1, Htr7, Hunk, Hus1, Huwe1, Hydın, Hyou1, Hypk, Iah1,
Iapp, Iars2, Iba57, Ibtk, Ica1l, Ick, Icmt, Icoslg, Id1, Ide, Idh1, Idh3a, Ids, Ier2, Ier5, Iffo1, Ifi16, Ifi44l, Ifi6,
Ifit1, Ifit2, Ifit3, Ifit5, Ifltd1, Ifna6, Ifnar1, Ifnar2, Ifne, Ifngr2, Ifnlr1, Ifrg15, Ift140, Ift20, Ift57, Ift80, Ift81,
Ift88, Igf1r, Igf2, Igf2bp2, Igf2bp3, Igfbp2, Igfbp5, Igj, Igsf11, Igsf23, Igsf6, Igsf9b, Ik, Ikbkap, Ikbkg, Ikzf3,
Ikzf4, Il10ra, Il11, Il12a, Il12rb1, Il12rb2, Il15, Il17ra, Il17rb, Il17rd, Il1b, Il1r2, Il1rl1, Il1rn, Il20, Il21r,
Il22ra2, Il23r, Il24, Il33, Il4, Il5ra, Il6r, Ildr2, Ilf2, Ilf3, Imp4, Impa1, Impa2, Impact, Impad1, Inadl, Incenp,
Ing1, Ing2, Ing5, Inhbc, Inhbe, Inmt, Inpp4a, Inpp4b, Inpp5a, Inpp5b, Inpp5e, Inpp5f, Insc, Insm1, Insm2,
Ints10, Ints2, Ints3, Ints6, Ints7, Intu, Ip6k1, Ip6k3, Ipcef1, Ipo5, Ipo8, Ipp, Ippk, Iqce, Iqcg, Iqcj, Iqgap1,
Iqgap2, Iqgap3, Iqub, Irak1bp1, Irak2, Irak3, Irak4, Irf4, Irf6, Irgq, Irs2, Irs4, Isca1, Isl1, Isl2, Islr2, Isoc1,
Ispd, Ist1, Isy1, Itga1, Itga10, Itga3, Itga4, Itga8, Itga9, Itgam, Itgav, Itgax, Itgb1, Itgb3bp, Itgb5, Itgb6,
Itm2a, Itm2b, Itpa, Itpk1, Itpr1, Itpr2, Itprıp, Iws1, Iyd, Jak1, Jak2, Jak3, Jakmıp2, Jam2, Jdp2, Jkamp,
Jmjd1c, Jmy, Josd1, Jph3, Jrkl, Junb, Jund, Kal1, Kank4, Kansl1, Kars, Kat2b, Kat6a, Kat6b, Kat7, Katnal2,
Katnbl1, Kazald1, Kb-1507c5.2, Kbtbd12, Kbtbd4, Kbtbd6, Kbtbd7, Kcna1, Kcna3, Kcna6, Kcnab1, Kcnc1,
Kcnc2, Kcnd2, Kcnd3, Kcne3, Kcne4, Kcng1, Kcng3, Kcnıp4, Kcnj13, Kcnj14, Kcnj15, Kcnj16, Kcnj5,
Kcnk1, Kcnk13, Kcnk3, Kcnk5, Kcnk6, Kcnk9, Kcnmb1, Kcnmb2, Kcnmb3, Kcnmb4, Kcnq3, Kcns1, Kcnt2,
Kctd1, Kctd10, Kctd12, Kctd2, Kctd3, Kctd5, Kctd7, Kctd8, Kdelr1, Kdelr2, Kdm1b, Kdm3a, Kdm3b, Kdm4a,
Kdm4b, Kdm4c, Kdm5a, Kdm5b, Kdm5c, Kdm5d, Kdm6b, Kdm8, Kdr, Kdsr, Kera, Khdc3l, Khdrbs2, Khsrp,
Kıaa0141, Kıaa0226, Kıaa0232, Kıaa0247, Kıaa0319, Kıaa0319l, Kıaa0368, Kıaa0391, Kıaa0408,
Kıaa0430, Kıaa0586, Kıaa0754, Kıaa0825, Kıaa0895, Kıaa0907, Kıaa0922, Kıaa0930, Kıaa0947, Kıaa1009,
Kıaa1024, Kıaa1033, Kıaa1107, Kıaa1109, Kıaa1143, Kıaa1191, Kıaa1210, Kıaa1239, Kıaa1324,
Kıaa1324l, Kıaa1429, Kıaa1430, Kıaa1432, Kıaa1456, Kıaa1467, Kıaa1522, Kıaa1524, Kıaa1549,
Kıaa1551, Kıaa1586, Kıaa1715, Kıaa1737, Kıaa1755, Kıaa1841, Kıaa1875, Kıaa1919, Kıaa1958, Kıaa2018,
Kıaa2022, Kıdıns220, Kıf11, Kıf14, Kıf16b, Kıf18b, Kıf1c, Kıf21b, Kıf24, Kıf27, Kıf2a, Kıf5c, Kıf6, Kın,
Kır2dl4, Kırrel, Kıtlg, Kl, Klf11, Klf16, Klf17, Klf3, Klf5, Klf8, Klf9, Klhdc1, Klhdc10, Klhdc8a, Klhl13,
Klhl2, Klhl21, Klhl23, Klhl24, Klhl26, Klhl29, Klhl30, Klhl31, Klhl32, Klhl36, Klhl38, Klhl4, Klhl5, Klhl6,
Klhl8, Klhl9, Klk10, Klk13, Klk4, Klk7, Klln, Klrc4, Klrd1, Kmo, Kmt2a, Kmt2c, Kmt2d, Knop1, Kpna1,
Kpna5, Krba1, Krba2, Krbox4, Krcc1, Krıt1, Krr1, Krt12, Krt2, Krt222, Krt26, Krt37, Krt74, Krt77, Krt78,
Krt8, Krt86, Krtap1-5, Krtap19-6, Krtap20-2, Krtap4-2, Krtap5-4, Krtap9-2, Krtap9-6, Krtap9-8, Ksr1, Ksr2,

*Ktı12, Ktn1, Kxd1, Kynu, L2hgdh, L3hypdh, L3mbtl4, Lacc1, Lace1, Lactb, Laır1, Lama4, Lamc3, Lamp2, Lamtor1, Lamtor3, Lancl1, Lancl2, Lancl3, Lap3, Laptm4a, Laptm4b, Laptm5, Larp1, Lars, Lat2, Lats1, Lats2, Lax1, Layn, Lbr, Lbx1, Lbx2, Lca5, Lcorl, Lcp1, Lcp2, Ldb1, Ldb2, Ldha, Ldhal6a, Ldlrad3, Ldoc1l, Lef1, Lefty2, Lemd2, Leng8, Lepr, Leprot, Leprotl1, Letm1, Letm2, Lfng, Lgals8, Lgı2, Lgmn, Lgr4, Lgsn, Lhfpl4, Lhpp, Lhx1, Lhx3, Lhx4, Lhx5, Lhx6, Lhx8, Lıfr, Lıg3, Lılra1, Lılra2, Lılra5, Lılrb4, Lılrb5, Lıma1, Lımd1, Lımk1, Lın52, Lın54, Lın7a, Lın7c, Lınc00346, Lınc00908, Lınc00998, Lıngo2, Lıngo3, Lıpc, Lıpg, Lıph, Lıtaf, Lıx1, Llph, Lman1, Lman2l, Lmbrd1, Lmbrd2, Lmnb1, Lmo3, Lmo7, Lmod2, Lmtk2, Lmx1b, Lnx1, Lnx2, Loh12cr1, Lonp2, Lonrf2, Lox, Loxl2, Lpar2, Lpar3, Lpar4, Lpcat2, Lpcat3, Lpcat4, Lpın1, Lpın2, Lpl, Lppr4, Lrat, Lrba, Lrıf1, Lrıt3, Lrp11, Lrp1b, Lrp2bp, Lrp3, Lrp4, Lrpap1, Lrpprc, Lrrc1, Lrrc10, Lrrc17, Lrrc19, Lrrc2, Lrrc25, Lrrc27, Lrrc28, Lrrc39, Lrrc3c, Lrrc47, Lrrc49, Lrrc4b, Lrrc55, Lrrc69, Lrrc8d, Lrrd1, Lrrfıp1, Lrrfıp2, Lrrk2, Lrrn1, Lrrn4cl, Lrrtm3, Lrrtm4, Lrtomt, Lsg1, Lsm10, Lsm11, Lsm14a, Lsm3, Lsm4, Lsm5, Lst1, Ltbp1, Ltbp2, Ltbr, Ltn1, Ltv1, Luc7l2, Luc7l3, Lum, Lurap1, Luzp2, Luzpp1, Ly6g5b, Ly75, Ly75-Cd302, Lyar, Lyn, Lypd1, Lypd3, Lypla1, Lyplal1, Lyrm1, Lyrm2, Lyrm4, Lyrm5, Lyrm7, Lyrm9, Lysmd1, Lysmd3, Lysmd4, Lyst, Lyz, Lztfl1, Lzts1, Lzts2, M6pr, Mab21l1, Mab21l3, Macc1, Macrod2, Mad2l1, Mad2l1bp, Maf, Mafb, Mafg, Mafk, Mag, Magea9, Magea9b, Mageb1, Mageb10, Mageb18, Mageb2, Mageb4, Mageb5, Magı1, Magı3, Magohb, Magt1, Mak, Mal, Mal2, Mall, Malsu1, Malt1, Mamdc2, Maml2, Maml3, Man1a1, Man1a2, Man2a1, Mansc1, Maob, Map10, Map1lc3c, Map2k2, Map3k4, Map3k7, Map4, Map4k4, Map4k5, Map7, Map7d2, Map7d3, Map9, Mapk13, Mapk1ıp1l, Mapk4, Mapkapk5, Mapre1, Marc2, March1, March2, March3, March7, March8, Marcksl1, Mark1, Marveld2, Mast3, Mat1a, Mat2b, Matn1, Matn2, Matn3, Matr3, Mavs, Mb, Mb21d1, Mbd3, Mbd5, Mbıp, Mbl2, Mblac2, Mboat1, Mboat2, Mbtps1, Mbtps2, Mc2r, Mccc2, Mcf2l, Mcf2l2, Mcıdas, Mcl1, Mcm10, Mcm3, Mcm3ap, Mcm4, Mcm7, Mcm8, Mcm9, Mcmbp, Mcmdc2, Mcoln2, Mcoln3, Mcph1, Mctp2, Mcts1, Mcu, Mcur1, Mdc1, Mdfı, Mdm1, Mdm2, Me1, Me2, Meaf6, Med1, Med14, Med16, Med17, Med18, Med20, Med21, Med22, Med28, Med4, Med6, Med9, Mef2bnb, Mefv, Megf9, Meı4, Meıs2, Melk, Mesdc1, Mesdc2, Mesp2, Mest, Metap2, Metrnl, Mettl10, Mettl12, Mettl14, Mettl16, Mettl17, Mettl20, Mettl21a, Mettl21b, Mettl24, Mettl2a, Mettl2b, Mettl4, Mettl6, Mettl7a, Mettl8, Mettl9, Mex3b, Mex3d, Mfap1, Mfap3, Mff, Mfhas1, Mfn1, Mfn2, Mfsd1, Mfsd2a, Mfsd4, Mfsd8, Mgam, Mgat1, Mgat2, Mgat3, Mgat4a, Mgat4c, Mgat5b, Mgea5, Mgp, Mgrn1, Mgst1, Mıa3, Mıca, Mıcalcl, Mıcall1, Mıcb, Mıd2, Mıdn, Mına, Mınos1-Nbl1, Mınpp1, Mıp, Mıs12, Mıs18bp1, Mıtf, Mkı67, Mkı67ıp, Mkln1, Mkrn3, Mks1, Mlana, Mlf1, Mlf2, Mlh1, Mlh3, Mlıp, Mllt11, Mllt3, Mllt4, Mmaa, Mmab, Mmachc, Mmadhc, Mmd, Mme, Mmp15, Mmp17, Mmp24, Mmp7, Mmp8, Mmp9, Mms22l, Mnat1, Mnda, Mns1, Mob1a, Mob3a, Mob3b, Mob3c, Mob4, Mobp, Mocos, Mocs2, Mocs3, Mogat2, Mon1b, Morc1, Morc2, Morc3, Morn2, Morn4, Mospd2, Moxd1, Mpc2, Mpdz, Mphosph6, Mphosph8, Mpı, Mplkıp, Mpp3, Mpp5, Mpp7, Mppe1, Mpped1, Mpst, Mpv17l, Mpzl1, Mpzl3, Mr1, Mrap2, Mre11a, Mreg, Mrfap1, Mrfap1l1, Mrı1, Mrm1, Mro, Mroh1, Mrp63, Mrpl12, Mrpl15, Mrpl19, Mrpl3, Mrpl30, Mrpl39, Mrpl4, Mrpl44, Mrpl45, Mrpl46, Mrpl47, Mrpl48, Mrpl49, Mrpl50, Mrpl51, Mrps10, Mrps11, Mrps14, Mrps16, Mrps17, Mrps21, Mrps25, Mrps27, Mrps30, Mrps36, Mrps5, Mrps6, Mrrf, Mrs2, Mrto4, Ms4a1, Ms4a10, Ms4a6a, Ms4a7, Msantd2, Msantd3, Msantd3-Tmeff1, Msantd4, Msh2, Msl2, Msl3, Msmb, Msn, Msr1, Msrb1, Msrb2, Mss51, Mstn, Mt1g, Mt1x, Mta1, Mta3, Mt-Atp6, Mtbp, Mterf, Mterfd3, Mtf1, Mtfmt, Mtfr2, Mthfd1, Mthfd1l, Mthfd2, Mthfr, Mthfsd, Mtıf2, Mtl5, Mtm1, Mtmr1, Mtmr11, Mtmr14, Mtmr2, Mtmr4, Mt-Nd4l, Mto1, Mtpap, Mtr, Mtrr, Mtss1, Mttp, Mtx2, Muc13, Muc15, Muc20, Muc7, Mum1l1, Murc, Mutyh, Mxra5, Mxra7, Mybbp1a, Mybl2, Mybpc1, Myc, Mycbp2, Mycl, Myct1, Myf6, Myh15, Myl12b, Mylk3, Mylk4, Mynn, Myo10, Myo18a, Myo1b, Myo1f, Myo1h, Myo3a, Myo3b, Myo5a, Myo5b, Myo5c, Myo6, Myo9a, Myog, Myom2, Myot, Myoz2, Mysm1, Mzt1, N4bp2, N4bp2l2, N4bp3, N6amt1, Naa11, Naa25, Naa30, Naa35, Naa38, Naa50, Naalad2, Nab1, Nabp1, Naca, Naca2, Nadk, Nadk2, Nagk, Nags, Naıf1, Naıp, Nalcn, Nampt, Nanog, Nanos1, Nanos2, Nap1l1, Nap1l3, Nap1l5, Napepld, Napg, Narf, Narg2, Nars, Nat1, Nat10, Nat8, Nat8l, Nav1, Nbl1, Nbn, Nbpf10, Nbpf11, Nbpf14, Nbpf24, Nbpf6, Ncam1, Ncam2, Ncan, Ncapd3, Ncapg2, Ncaph, Ncbp1, Nccrp1, Nceh1, Nck2, Nckap5l, Ncl, Ncmap, Ncoa1, Ncoa2, Ncoa3, Ncoa4, Ncoa5, Ncoa7, Ncor1, Ncor2, Ncr1, Ncr3lg1, Nde1, Ndnl2, Ndp, Ndrg1, Ndrg3, Ndrg4, Ndst1, Ndst3, Ndst4, Ndufa10, Ndufa4, Ndufa5, Ndufa7, Ndufa9, Ndufaf3, Ndufaf4, Ndufaf5, Ndufaf6, Ndufb5, Ndufb6, Ndufc1, Ndufc2, Ndufs1, Ndufs5, Ndufv3, Necab3, Necap2, Nedd1, Nedd4l, Nedd9, Neıl2, Nek1, Nek2, Nek3, Nek4, Nek5, Nek7, Nelfa, Nell1, Nenf, Neo1, Net1, Neto1, Neto2, Neu3, Neurl, Neurod2, Neurog2, Nf2, Nfam1, Nfatc2ıp, Nfe2l1, Nfe2l2, Nfe2l3, Nfkbız, Nfrkb, Nfxl1, Nfya, Nfyb, Nfyc, Ngb, Nhej1, Nhlh2, Nhlrc2, Nhsl1, Nıcn1, Nıd1, Nıf3l1, Nın, Nınj1, Nıpa1, Nıpa2, Nıpal2, Nıpal3, Nıpal4, Nıpsnap3a, Nıpsnap3b, Nıt2, Nkaın1, Nkaın2, Nkaın3, Nkap, Nkapl, Nkıras1, Nkx1-2, Nkx2-1, Nkx2-2, Nkx3-1, Nlgn1, Nlgn2, Nlgn4x, Nln, Nlrc3, Nlrc5, Nlrp3, Nlrp9, Nme6, Nmnat1, Nmnat2, Nmnat3, Nnmt, Nnt, Noa1, Noc2l, Noc3l, Nol10, Nol9, Nolc1, Nom1, Nono, Nop14, Nop9, Nos1, Notch1, Notch2, Notch2nl, Noto, Nox1, Nox5, Npap1, Npat, Npepps, Npffr1, Nphp1, Npl, Npnt, Npr1, Npr3, Nptx1, Npy1r, Npy4r, Npy5r, Nqo1, Nqo2, Nr1ı2, Nr2c1, Nr2c2, Nr2f1, Nr2f6, Nr3c1, Nr4a1, Nr4a3, Nr5a2, Nr6a1, Nras, Nrbp2, Nrcam, Nrde2, Nrg1, Nrg2, Nrg4, Nrıp2, Nrıp3, Nrn1, Nrp1, Nrxn1, Nrxn2, Nrxn3, Nsd1, Nsg2, Nsl1, Nsmce1, Nsun3, Nsun4, Nt5c1a, Nt5dc1, Nt5dc3, Ntm, Ntn1, Ntng1, Ntng2, Ntpcr, Ntrk2, Nts, Nuak2, Nub1, Nubpl, Nucb1, Nucb2, Nucks1, Nudcd1, Nudcd2, Nudt1, Nudt10, Nudt11, Nudt16,*

*Nudt19, Nudt21, Nudt3, Nudt4, Nudt7, Nufıp1, Nuggc, Numbl, Nup107, Nup153, Nup155, Nup205, Nup214, Nup35, Nup43, Nup50, Nup62cl, Nup85, Nup93, Nupl1, Nupl2, Nusap1, Nutm2d, Nxf1, Nxn, Nxpe3, Nxph2, Nxt1, Nxt2, Oaf, Oard1, Oat, Oaz2, Ocıad2, Ocln, Odf2, Odf2l, Ogfod1, Ogfod3, Ogfrl1, Ogt, Oıp5, Ola1, Olfm2, Olfm3, Olfml1, Olfml2a, Olfml2b, Olıg2, Olr1, Omd, Opa1, Ophn1, Opn3, Opn5, Oprk1, Optn, Or12d3, Or2a4, Or2c3, Or2h1, Or51e2, Or7d2, Or9q1, Oraı2, Oraı3, Oraov1, Orc1, Orc4, Orc6, Ormdl1, Ormdl3, Os9, Osbp, Osbpl10, Osbpl11, Osbpl2, Osbpl3, Osbpl8, Oser1, Osgep, Osgepl1, Osgın2, Osmr, Ost4, Ostc, Ostf1, Ostm1, Ostn, Otc, Otog, Otub2, Otud3, Otud6b, Otud7a, Ovol1, Oxgr1, Oxnad1, Oxsr1, Oxtr, P2rx7, P2ry1, P2ry10, P2ry14, P2ry2, P2ry8, P4hb, Pabpc1, Pabpc5, Pabpn1, Pacrgl, Padı4, Paf1, Pafah1b2, Pag1, Pah, Paıcs, Paıp2, Pak1, Pak1ıp1, Pak2, Pak3, Pak4, Palb2, Palld, Palm2, Palm2-Akap2, Pan3, Pank2, Pank3, Paox, Papd7, Papln, Papolb, Papolg, Pappa2, Papss1, Paqr3, Paqr5, Paqr7, Paqr8, Pard3, Pard3b, Pard6b, Pard6g, Parg, Parp1, Parp14, Parp15, Parp16, Parp2, Parpbp, Patz1, Pawr, Pax1, Pax3, Pax9, Paxbp1, Pbx1, Pbx2, Pbx4, Pcbd1, Pcbp1, Pcbp2, Pcbp4, Pccb, Pcdh11x, Pcdh11y, Pcdh19, Pcdh7, Pcdh9, Pcdhb10, Pcdhb11, Pcdhb12, Pcdhb13, Pcdhb15, Pcdhb16, Pcdhb17, Pcdhb2, Pcdhb3, Pcdhb4, Pcdhb5, Pcdp1, Pcf11, Pcgf1, Pcgf2, Pcgf3, Pck1, Pclo, Pcm1, Pcmt1, Pcmtd1, Pcmtd2, Pcna, Pcnx, Pcnxl2, Pcnxl3, Pcsk1, Pcsk2, Pcsk5, Pcsk7, Pcsk9, Pcyt1a, Pcyt1b, Pdcd10, Pdcd11, Pdcd1lg2, Pdcd6ıp, Pdcd7, Pdcl, Pdcl3, Pde11a, Pde12, Pde1c, Pde3a, Pde4d, Pde4dıp, Pde6b, Pde6d, Pdf, Pdgfa, Pdgfc, Pdgfd, Pdgfra, Pdhb, Pdhx, Pdıa3, Pdk1, Pdk3, Pdlım2, Pdlım3, Pdlım4, Pdlım5, Pdp1, Pdp2, Pdpn, Pdpr, Pdrg1, Pds5a, Pds5b, Pdxdc1, Pdxk, Pdzd2, Pdzd8, Pdzd9, Pdzrn4, Peak1, Peg3, Pelı1, Pelı2, Pelo, Pelp1, Penk, Per1, Per2, Per3, Pet112, Pet117, Pex1, Pex10, Pex11a, Pex11b, Pex13, Pex19, Pex2, Pex26, Pex3, Pex7, Pf4, Pfas, Pfdn2, Pfkfb2, Pfkm, Pgam5, Pgap1, Pgap2, Pgbd1, Pgbd2, Pgbd4, Pgbd5, Pgd, Pggt1b, Pgls, Pgm2, Pgpep1, Pgr, Phactr2, Phactr3, Phb, Phc3, Phex, Phf10, Phf11, Phf16, Phf2, Phf20l1, Phf21b, Phf3, Phf5a, Phf6, Phf7, Phf8, Phka1, Phkb, Phkg2, Phox2a, Phox2b, Pı4k2a, Pı4k2b, Pıas1, Pıas2, Pıas3, Pıas4, Pıezo2, Pıga, Pıgk, Pıgm, Pıgo, Pıgp, Pıgr, Pıgu, Pıgv, Pıgw, Pıgy, Pıh1d3, Pık3c2b, Pık3c3, Pık3cd, Pık3ıp1, Pım1, Pım2, Pınx1, Pıp4k2a, Pıp4k2b, Pıp4k2c, Pıp5k1a, Pırt, Pısd, Pıthd1, Pıtpnb, Pıtpnc1, Pıwıl1, Pıwıl2, Pıwıl3, Pja1, Pkd1l1, Pkd2, Pkhd1l1, Pkıb, Pkn2, Pknox2, Pkp1, Pkp2, Pla1a, Pla2g12a, Pla2g16, Pla2g2c, Pla2g4a, Pla2g4d, Pla2g4e, Pla2g7, Pla2r1, Plac8, Plagl2, Plb1, Plbd2, Plcb2, Plcd3, Plce1, Plcg1, Plcg2, Plcxd1, Plcxd3, Pld1, Pld5, Pld6, Plek, Plekha1, Plekha2, Plekha3, Plekha5, Plekha6, Plekha7, Plekha8, Plekhb2, Plekhf2, Plekhg1, Plekhg2, Plekhg3, Plekhg4b, Plekhg7, Plekhh2, Plekhm3, Plekho2, Plg, Plglb1, Plglb2, Plın1, Plk4, Pllp, Pln, Plod2, Plrg1, Pls1, Pls3, Plscr1, Plscr4, Plxdc1, Plxna1, Plxna2, Plxna3, Pm20d2, Pmaıp1, Pml, Pmp22, Pmpca, Pmpcb, Pms2, Pmvk, Pnısr, Pnlıprp3, Pnma1, Pnn, Pno1, Pnoc, Pnp, Pnpla1, Pnpla3, Pnpla4, Pnpo, Podxl, Pof1b, Pofut1, Poglut1, Pola2, Pole3, Pole4, Polh, Polı, Polm, Polr1a, Polr1b, Polr1c, Polr1d, Polr1e, Polr2a, Polr2d, Polr2j2, Polr2j3, Polr2k, Polr3a, Polr3d, Polr3e, Polr3k, Pom121l2, Pomk, Pomp, Pomt2, Pon1, Pon3, Pop1, Pop5, Popdc2, Popdc3, Pot1, Poted, Potee, Poteg, Poteı, Potej, Potem, Pou2af1, Pou2f3, Pou3f1, Pou5f1b, Pou6f1, Pou6f2, Ppa2, Ppapdc1a, Ppapdc2, Ppargc1a, Ppargc1b, Ppat, Ppbp, Ppef2, Ppfıa4, Ppfıbp1, Pphln1, Ppıc, Ppıe, Ppıf, Ppıh, Ppıl2, Ppıl3, Ppıl6, Ppıp5k2, Ppm1e, Ppm1f, Ppm1h, Ppm1k, Ppm1l, Ppm1n, Ppme1, Ppp1cb, Ppp1cc, Ppp1r10, Ppp1r12b, Ppp1r12c, Ppp1r13b, Ppp1r14c, Ppp1r15a, Ppp1r1a, Ppp1r1c, Ppp1r2, Ppp1r3a, Ppp1r3b, Ppp1r3c, Ppp1r3d, Ppp1r3e, Ppp1r9a, Ppp2r1a, Ppp2r1b, Ppp2r2a, Ppp2r2b, Ppp2r2c, Ppp2r3a, Ppp3cc, Ppp3r1, Ppp4r1l, Ppp4r2, Ppp5c, Ppt1, Pqlc2, Pramef16, Pramef17, Prcp, Prdm1, Prdm13, Prdm15, Prdm16, Prdm2, Prdm6, Prdm7, Prdm9, Prelıd2, Prelp, Prep, Prepl, Prex1, Prex2, Prh2, Prıckle1, Prıckle3, Prım1, Prım2, Prkab1, Prkaca, Prkag2, Prkar2b, Prkcb, Prkce, Prkcı, Prkd2, Prkd3, Prkg1, Prkg2, Prkrıp1, Prkrır, Prlr, Prmt10, Prmt2, Prmt6, Prnd, Prom2, Prosc, Prox2, Prpf3, Prpf31, Prpf38a, Prpf38b, Prpf39, Prpf4, Prpf40a, Prpf40b, Prpf4b, Prpf6, Prps2, Prr11, Prr12, Prr13, Prr23a, Prr23c, Prr24, Prr26, Prrc1, Prrc2b, Prrc2c, Prrg1, Prrg3, Prrg4, Prrt1, Prss16, Prss21, Prss23, Prss35, Prss53, Prss54, Psat1, Psd2, Psd4, Psen1, Psenen, Psg11, Psıp1, Psma4, Psma5, Psma8, Psmb1, Psmb2, Psmb3, Psmb5, Psmb6, Psmb9, Psmc2, Psmc4, Psmc6, Psmd10, Psmd12, Psmd3, Psmd4, Psmd5, Psmd9, Psme4, Psmf1, Psmg1, Psmg4, Pspc1, Psph, Pstpıp2, Ptcd1, Ptcd2, Ptcd3, Ptch1, Ptch2, Ptchd1, Ptchd3, Ptdss2, Pter, Ptgdr, Ptger3, Ptger4, Ptges3l, Ptgfrn, Ptgıs, Ptgr2, Ptgs2, Pthlh, Ptk2, Ptk6, Ptk7, Ptma, Ptn, Ptp4a2, Ptpla, Ptplad1, Ptplad2, Ptplb, Ptpn1, Ptpn11, Ptpn12, Ptpn20a, Ptpn20b, Ptpn3, Ptpre, Ptprf, Ptprj, Ptprk, Ptprm, Ptprq, Ptprr, Ptprs, Ptprt, Ptpru, Ptrf, Ptrh2, Pttg1ıp, Ptx3, Pum1, Pum2, Pus7, Pus7l, Pvr, Pvrl1, Pvrl3, Pwwp2b, Pxk, Pxmp4, Pycard, Pygo1, Pygo2, Pyroxd1, Pyurf, Qdpr, Qpct, Qpctl, Qrfpr, Qrıch1, Qser1, Qtrt1, Qtrtd1, R3hdm1, Rab10, Rab11fıp1, Rab11fıp2, Rab11fıp5, Rab12, Rab19, Rab27a, Rab27b, Rab2a, Rab30, Rab31, Rab32, Rab36, Rab37, Rab39a, Rab3d, Rab3gap2, Rab3ıp, Rab40a, Rab40b, Rab40c, Rab5a, Rab6b, Rab6c, Rab7l1, Rab9b, Rabac1, Rabep1, Rabgap1l, Rabgef1, Rabıf, Rabl2a, Rabl2b, Rabl3, Rabl5, Rac1, Rad1, Rad18, Rad21l1, Rad50, Rad51ap2, Rad51b, Rad51l3-Rffl, Rad52, Rad54b, Rad54l, Rad9b, Raet1e, Rag1, Rag2, Raı14, Rala, Ralb, Ralbp1, Ralgapa2, Ralgapb, Ramp2, Ran, Ranbp1, Ranbp10, Ranbp3l, Ranbp6, Rangap1, Rap1gds1, Rapgef1, Rapgef3, Rapgef5, Rapgef6, Rara, Rarb, Rarres1, Rars2, Rasa2, Rasa3, Rasa4, Rasal2, Rasef, Rasgef1b, Rasgrf2, Rasgrp1, Rasgrp4, Rasl10b, Rasl11b, Rassf3, Rassf4, Rassf5, Rassf6, Rassf8, Raver2, Rb1, Rb1cc1, Rbak, Rbbp4, Rbbp5, Rbbp6, Rbbp7, Rbbp9, Rbfox1, Rbl1, Rbl2, Rbm11, Rbm12b, Rbm15, Rbm15b, Rbm17, Rbm18, Rbm19, Rbm22, Rbm23, Rbm25, Rbm27, Rbm28, Rbm3,*

*Rbm33, Rbm34, Rbm38, Rbm41, Rbm43, Rbm45, Rbm46, Rbm47, Rbm48, Rbm4b, Rbm7, Rbm8a, Rbms2, Rbmx2, Rbmxl1, Rbmy1a1, Rbmy1b, Rbmy1d, Rbmy1e, Rbmy1f, Rbmy1j, Rbp2, Rbpj, Rbpms2, Rcan1, Rcbtb1, Rcbtb2, Rcl1, Rcn2, Rcor1, Rd3, Rdh10, Rdh11, Rdh13, Rdh16, Rdx, Reck, Recql, Reep5, Rel, Relb, Reln, Rer1, Rere, Rergl, Ret, Retsat, Rfc2, Rfc3, Rfc5, Rffl, Rfk, Rfng, Rft1, Rftn2, Rfwd3, Rfx3, Rfxap, Rgma, Rgpd1, Rgpd2, Rgpd4, Rgpd5, Rgpd6, Rgpd8, Rgr, Rgs1, Rgs16, Rgs17, Rgs18, Rgs22, Rgs4, Rgs7bp, Rgs9bp, Rhbdd1, Rhbg, Rhcg, Rhd, Rhno1, Rhobtb1, Rhof, Rhoj, Rhoq, Rhou, Rhpn2, Ric8b, Rif1, Rilpl1, Rimklb, Riok2, Riok3, Ripk4, Ripply1, Ripply3, Rit2, Rlf, Rln2, Rmdn1, Rmdn2, Rmi1, Rmi2, Rmnd5a, Rnase13, Rnase4, Rnaseh1, Rnaseh2b, Rnd2, Rnd3, Rnf10, Rnf103, Rnf103-Chmp3, Rnf122, Rnf125, Rnf126, Rnf13, Rnf138, Rnf139, Rnf14, Rnf141, Rnf144a, Rnf144b, Rnf146, Rnf149, Rnf150, Rnf168, Rnf169, Rnf170, Rnf182, Rnf183, Rnf187, Rnf20, Rnf207, Rnf212, Rnf213, Rnf214, Rnf217, Rnf24, Rnf41, Rnf43, Rnf6, Rnf7, Rnft1, Rngtt, Rnmt, Rnmtl1, Rnpc3, Rnps1, Robo1, Rock2, Ror2, Ros1, Rp1, Rp11-1026m7.2, Rp11-105c20.2, Rp11-1070n10.3, Rp11-1102p16.1, Rp11-111m22.2, Rp11-113d6.10, Rp11-116d17.1, Rp11-122a3.2, Rp11-156e8.1, Rp11-162a12.2, Rp11-169f17.1, Rp11-181c3.1, Rp11-192h23.4, Rp11-210m15.2, Rp11-268j15.5, Rp11-302b13.5, Rp11-322l20.1, Rp11-332o19.5, Rp11-343c2.12, Rp11-366l20.2, Rp11-382j12.1, Rp11-383h13.1, Rp11-422n16.3, Rp11-455g16.1, Rp11-497e19.2, Rp11-625h11.1, Rp11-644f5.10, Rp11-664d7.4, Rp11-67h2.1, Rp11-762i7.5, Rp11-817j15.3, Rp11-826n14.2, Rp11-9b6.1, Rp13-996f3.4, Rp13-996f3.5, Rp2, Rp5-1052i5.2, Rpa1, Rpa2, Rpa3, Rpa3-As1, Rpap2, Rpe, Rpe65, Rpf1, Rpf2, Rpgr, Rpgrip1l, Rph3al, Rpl10, Rpl10a, Rpl10l, Rpl12, Rpl13, Rpl14, Rpl15, Rpl17-C18orf32, Rpl18a, Rpl22, Rpl22l1, Rpl23, Rpl24, Rpl27a, Rpl28, Rpl32, Rpl35a, Rpl36a-Hnrnph2, Rpl37, Rpl37a, Rpl3l, Rpl4, Rpl7, Rpl7a, Rplp1, Rpn2, Rpp30, Rpp40, Rprd1a, Rprd1b, Rprml, Rps12, Rps16, Rps19, Rps21, Rps27l, Rps28, Rps29, Rps3, Rps6, Rps6ka3, Rps6ka6, Rps6kb1, Rps6kc1, Rpsap58, Rptn, Rraga, Rreb1, Rrh, Rrm1, Rrm2, Rrm2b, Rrn3, Rrp1, Rrp15, Rrp36, Rrp8, Rs1, Rsad2, Rsl1d1, Rsl24d1, Rsph3, Rsph9, Rspo2, Rspo3, Rspry1, Rsrc1, Rsrc2, Rsu1, Rtca, Rtcb, Rtf1, Rtn3, Rtn4ip1, Rtp1, Rufy2, Rufy3, Rundc1, Rundc3b, Runx1, Runx2, Rwdd2a, Rwdd2b, Rwdd3, Rwdd4, Rxfp1, Ryk, S100a16, S100a2, S100a6, S1pr5, Sacs, Safb, Sall1, Sall3, Sall4, Samd15, Samd5, Samd9, Samhd1, Sap30, Sap30l, Sart3, Sash1, Sass6, Satb2, Sav1, Sbk2, Sbno1, Sbspon, Scamp1, Scand3, Scarb2, Scd, Scd5, Scel, Scfd2, Scg3, Scg5, Scgn, Scimp, Scin, Scly, Scn11a, Scn1a, Scn9a, Scnm1, Scnn1g, Sco1, Scoc, Scrn3, Scube2, Scxa, Scxb, Scyl3, Sdad1, Sdc3, Sdc4, Sdcbp, Sdccag3, Sde2, Sdf4, Sdha, Sdhaf1, Sdhaf2, Sdhc, Sdhd, Sdk2, Sdpr, Sdr16c5, Sec14l1, Sec14l2, Sec14l3, Sec14l5, Sec16a, Sec16b, Sec22c, Sec23a, Sec23b, Sec23ip, Sec24a, Sec24b, Sec31a, Secisbp2, Seh1l, Sel1l, Sele, Selk, Selp, Selt, Sema3a, Sema3c, Sema4g, Sema5a, Sema6a, Sema6b, Sema6c, Senp1, Senp2, Senp5, Senp6, Senp7, Senp8, Sep15, Sepn1, Sepsecs, Sept10, Sept11, Sept14, Sept2, Sept3, Sept6, Serac1, Serbp1, Serf1a, Serf1b, Serf2, Serinc1, Serinc4, Serinc5, Serp1, Serp2, Serpina5, Serpinb1, Serpinb13, Serpinb5, Serpinb7, Serpinb8, Serpinb9, Serpine1, Serping1, Serpinh1, Sertad1, Sertad3, Sertad4, Sesn1, Sesn2, Set, Setd1b, Setd3, Setd7, Setd8, Setx, Sez6l, Sez6l2, Sf1, Sf3a1, Sf3b1, Sf3b14, Sf3b2, Sf3b3, Sf3b5, Sfmbt2, Sfpq, Sfr1, Sfrp1, Sfrp4, Sfrp5, Sfswap, Sft2d1, Sft2d2, Sft2d3, Sfxn1, Sfxn2, Sfxn3, Sfxn5, Sgce, Sgk223, Sgk3, Sgk494, Sgol1, Sgol2, Sgpl1, Sgpp1, Sgpp2, Sgsh, Sgsm1, Sgsm2, Sgtb, Sh2b3, Sh2d1a, Sh2d1b, Sh2d4b, Sh3bgr, Sh3bgrl, Sh3bp1, Sh3bp2, Sh3bp4, Sh3gl3, Sh3glb1, Sh3pxd2b, Sh3rf1, Sh3rf2, Sh3tc1, Sh3tc2, Shank2, Shb, Shc1, Shc4, She, Shh, Shisa2, Shisa7, Shisa9, Shmt1, Shmt2, Shox, Shox2, Shpk, Shprh, Shq1, Shroom2, Shroom4, Siae, Siah1, Siah3, Siglec11, Siglec14, Siglec6, Siglec9, Sik2, Sike1, Sim1, Sim2, Simc1, Sin3a, Sipa1l2, Sirt1, Sirt3, Six3, Six4, Ska1, Ska2, Skap2, Ski, Skor1, Skp2, Sla2, Slain1, Slain2, Slbp, Slc10a6, Slc10a7, Slc11a2, Slc12a1, Slc12a3, Slc12a6, Slc12a8, Slc13a1, Slc14a1, Slc15a1, Slc15a2, Slc15a5, Slc16a1, Slc16a13, Slc16a3, Slc16a4, Slc16a6, Slc16a9, Slc17a2, Slc17a3, Slc17a8, Slc18a2, Slc18b1, Slc19a1, Slc19a2, Slc19a3, Slc1a2, Slc1a3, Slc1a5, Slc20a1, Slc20a2, Slc22a1, Slc22a15, Slc22a23, Slc22a24, Slc22a25, Slc23a2, Slc24a1, Slc24a5, Slc25a11, Slc25a15, Slc25a16, Slc25a17, Slc25a23, Slc25a24, Slc25a26, Slc25a30, Slc25a32, Slc25a33, Slc25a36, Slc25a37, Slc25a38, Slc25a40, Slc25a42, Slc25a43, Slc25a45, Slc25a46, Slc25a51, Slc25a52, Slc25a53, Slc26a2, Slc26a3, Slc27a4, Slc28a3, Slc29a3, Slc29a4, Slc2a10, Slc2a12, Slc2a13, Slc2a2, Slc2a4, Slc2a5, Slc2a9, Slc30a1, Slc30a10, Slc30a6, Slc30a8, Slc30a9, Slc31a1, Slc33a1, Slc35a2, Slc35a3, Slc35a5, Slc35b4, Slc35d2, Slc35d3, Slc35e1, Slc35e2, Slc35e3, Slc35f2, Slc35f3, Slc35f5, Slc35f6, Slc35g1, Slc35g3, Slc36a1, Slc36a2, Slc36a3, Slc36a4, Slc37a4, Slc38a1, Slc38a2, Slc38a7, Slc39a10, Slc39a11, Slc39a12, Slc39a2, Slc39a7, Slc39a9, Slc3a1, Slc40a1, Slc41a2, Slc43a1, Slc43a2, Slc44a1, Slc44a3, Slc45a2, Slc47a1, Slc4a10, Slc4a1ap, Slc4a4, Slc4a7, Slc5a1, Slc5a12, Slc5a3, Slc5a5, Slc5a8, Slc6a1, Slc6a14, Slc6a20, Slc7a1, Slc7a11, Slc7a14, Slc7a7, Slc7a8, Slc9a4, Slc9a5, Slc9a7, Slc9a8, Slc9a9, Slco1b1, Slco1b3, Slco2b1, Slco4c1, Slco5a1, Slfn11, Slfn12, Slfn12l, Slfn5, Slit2, Slit3, Slitrk2, Slitrk3, Slitrk4, Slk, Slmo2, Sltm, Slu7, Slx4, Slx4ip, Smad1, Smad2, Smad3, Smad4, Smad6, Smad7, Smad9, Smagp, Smarca1, Smarca5, Smarcad1, Smarcc1, Smarcc2, Smc1a, Smc4, Smc5, Smchd1, Smco3, Smco4, Smek1, Smg7, Smim12, Smim13, Smim17, Smim19, Smim4, Smim7, Smndc1, Smpd3, Smpd4, Smpdl3a, Sms, Smtnl2, Smug1, Smyd4, Snai2, Snap23, Snap29, Snap47, Snap91, Snapc3, Snapc5, Snca, Sned1, Snf8, Snn, Snrnp35, Snrnp40, Snrnp48, Snrpa, Snrpb2, Snrpd1, Snrpd3, Snrpf, Snrpn, Sntb1, Sntb2, Sntn, Snx1, Snx10, Snx12, Snx13, Snx14, Snx15, Snx19, Snx20, Snx22, Snx24, Snx29, Snx3, Snx30, Snx31, Snx4, Snx6, Soat1, Sobp, Socs2, Socs4, Socs7, Sod2, Sorbs1, Sorbs2, Sorcs1, Sorcs3, Sord, Sort1, Sos1, Sost, Sostdc1, Sowaha,*

*Sowahc, Sox1, Sox11, Sox13, Sox17, Sox18, Sox2, Sox3, Sox30, Sox7, Sox8, Sox9, Sp110, Sp140l, Sp2, Sp3, Sp4, Sp8, Sp9, Spa17, Spaca1, Spag1, Spag16, Spag6, Spag9, Spam1, Spast, Spata13, Spata17, Spata18, Spata5, Spata6, Spata6l, Spats1, Spats2, Spc24, Spc25, Spcs2, Spcs3, Spdye1, Spdye3, Specc1, Specc1l, Spem1, Sphar, Spib, Spic, Spice1, Spidr, Spin1, Spin3, Spin4, Spint2, Spn, Spo11, Spocd1, Spop, Spopl, Sppl2a, Sppl3, Sprr2f, Sprtn, Spry4, Spryd4, Spryd7, Spta1, Sptbn1, Sptbn2, Sptlc1, Sptlc2, Sptlc3, Spz1, Sqle, Srbd1, Srcap, Srd5a1, Srd5a3, Srek1ip1, Srfbp1, Srgap1, Srgn, Srl, Srp19, Srp72, Srp9, Srpk1, Srprb, Srr, Srrd, Srrm1, Srrt, Srsf1, Srsf10, Srsf11, Srsf12, Srsf2, Srsf3, Srsf5, Srsf7, Srsf9, Ss18, Ss18l1, Ssb, Ssbp2, Ssbp3, Ssh2, Ssh3, Ssr1, Ssr3, Sstr1, Sstr2, Ssx2ip, St13, St18, St5, St6gal1, St6galnac2, St6galnac3, St7l, St7-Ot4, St8sia2, Stag2, Stam, Stambp, Stap1, Stard10, Stard13, Stard3nl, Stard4, Stard5, Stard7, Stard9, Stat2, Stat3, Stat5a, Stat5b, Stath, Stau1, Steap4, Stil, Stim1, Stim2, Stk11, Stk17a, Stk17b, Stk24, Stk25, Stk3, Stk32a, Stk32b, Stk36, Stk38l, Stk4, Stmn1, Stmn2, Stoml1, Ston1, Ston2, Stox1, Stpg1, Stradb, Strap, Strbp, Strip2, Strn, Sts, Stt3a, Stt3b, Stx11, Stx16, Stx17, Stx1b, Stx2, Stx4, Stx6, Stxbp3, Stxbp4, Stxbp5, Stxbp5l, Stxbp6, Styx, Sub1, Sucla2, Sucnr1, Suco, Suds3, Sugt1, Sulf2, Sult1a3, Sult1a4, Sult1b1, Sult1c4, Sult1e1, Sumf2, Sumo1, Sumo2, Supt16h, Supt3h, Supt7l, Surf4, Surf6, Susd1, Susd5, Suv420h1, Sv2b, Svil, Svip, Svop, Swap70, Sycp2, Sycp2l, Syde2, Syf2, Syn3, Sync, Synj1, Synj2, Synj2bp, Synrg, Sypl1, Sypl2, Syt10, Syt13, Syt14, Syt15, Syt17, Syt4, Syt5, Sytl3, T, Tab2, Taco1, Tacr1, Tacr2, Tacr3, Tacstd2, Tada2a, Tada2b, Taf11, Taf13, Taf1a, Taf1b, Taf2, Taf3, Taf4b, Taf5, Taf5l, Taf8, Taf9b, Tagln2, Tal1, Tamm41, Tanc2, Tank, Taok3, Tap2, Tapbp, Tapbpl, Tarsl2, Tas2r14, Tas2r20, Tas2r4, Tas2r5, Tatdn2, Tatdn3, Tax1bp1, Taz, Tbc1d1, Tbc1d14, Tbc1d19, Tbc1d23, Tbc1d24, Tbc1d2b, Tbc1d30, Tbc1d4, Tbc1d5, Tbc1d8b, Tbc1d9, Tbca, Tbcb, Tbccd1, Tbce, Tbck, Tbl1x, Tbl1xr1, Tbpl1, Tbrg1, Tbx20, Tbx21, Tbx5, Tc2n, Tcea1, Tcea3, Tceal4, Tceanc, Tceb1, Tcerg1, Tcf15, Tcf24, Tcf3, Tcf4, Tcf7, Tcf7l2, Tcfl5, Tchp, Tcof1, Tcp10l2, Tcp11l1, Tctex1d1, Tctn1, Tctn2, Tdgf1, Tdp1, Tdp2, Tdrd12, Tdrd15, Tdrd5, Tdrkh, Tdrp, Tead1, Tead3, Teddm1, Tenm2, Tenm3, Tep1, Tepp, Terf2, Terf2ip, Tert, Tes, Tesc, Tet2, Tex101, Tex15, Tex22, Tex26, Tex30, Tex9, Tfam, Tfap2c, Tfb1m, Tfcp2, Tfdp3, Tfpi, Tg, Tgif2, Tgif2lx, Tgif2ly, Tgs1, Thap2, Thap5, Thap6, Thap9, Thbd, Thbs2, Them4, Thg1l, Thnsl1, Thoc1, Thoc5, Thrap3, Thsd7b, Thumpd2, Thumpd3, Tiaf1, Tial1, Tiam1, Ticam2, Ticrr, Tifa, Tigd2, Tigd6, Timd4, Timeless, Timm10b, Timm22, Timm23, Timm50, Timm8a, Timm9, Timp2, Timp3, Tinagl1, Tipin, Tiprl, Tirap, Tjap1, Tjp1, Tk2, Tkt, Tlcd2, Tldc2, Tle1, Tle3, Tlk2, Tll1, Tll2, Tln1, Tlr3, Tlr4, Tlr6, Tlr7, Tlr8, Tm2d1, Tm4sf1, Tm4sf20, Tm4sf4, Tm4sf5, Tm6sf1, Tm7sf3, Tm9sf2, Tm9sf4, Tma16, Tmc5, Tmcc3, Tmco1, Tmco5a, Tmed10, Tmed2, Tmed4, Tmed5, Tmed7, Tmed7-Ticam2, Tmed8, Tmeff1, Tmem101, Tmem105, Tmem106a, Tmem106c, Tmem108, Tmem11, Tmem117, Tmem119, Tmem120b, Tmem123, Tmem126b, Tmem127, Tmem132c, Tmem133, Tmem135, Tmem140, Tmem145, Tmem14a, Tmem14e, Tmem150c, Tmem154, Tmem158, Tmem161b, Tmem163, Tmem167a, Tmem168, Tmem169, Tmem17, Tmem170a, Tmem170b, Tmem179b, Tmem182, Tmem183a, Tmem184b, Tmem184c, Tmem185a, Tmem185b, Tmem186, Tmem19, Tmem192, Tmem194b, Tmem2, Tmem200c, Tmem206, Tmem209, Tmem213, Tmem215, Tmem216, Tmem218, Tmem219, Tmem220, Tmem229a, Tmem229b, Tmem230, Tmem231, Tmem233, Tmem237, Tmem239, Tmem241, Tmem242, Tmem246, Tmem248, Tmem251, Tmem252, Tmem254, Tmem255a, Tmem261, Tmem27, Tmem30a, Tmem30b, Tmem30c, Tmem33, Tmem35, Tmem38a, Tmem38b, Tmem39a, Tmem40, Tmem41b, Tmem42, Tmem43, Tmem44, Tmem45b, Tmem50b, Tmem51, Tmem52b, Tmem57, Tmem65, Tmem66, Tmem67, Tmem78, Tmem86a, Tmem8a, Tmem91, Tmem92, Tmem97, Tmem99, Tmem9b, Tmf1, Tmod2, Tmod3, Tmpo, Tmppe, Tmprss11a, Tmprss11bnl, Tmprss11d, Tmprss11e, Tmprss12, Tmprss13, Tmprss2, Tmprss3, Tmprss4, Tmsb15a, Tmsb15b, Tmsb4x, Tmsb4y, Tmtc1, Tmtc2, Tmtc3, Tmx1, Tmx3, Tmx4, Tnfaip1, Tnfaip2, Tnfaip8, Tnfaip8l1, Tnfaip8l3, Tnfrsf10a, Tnfrsf10b, Tnfrsf10d, Tnfrsf11a, Tnfrsf13c, Tnfrsf1a, Tnfrsf6b, Tnfrsf9, Tnfsf10, Tnfsf11, Tnfsf15, Tnfsf4, Tnfsf8, Tnfsf9, Tnik, Tnks1bp1, Tnks2, Tnni1, Tnpo1, Tnpo3, Tnr, Tnrc18, Tns1, Tns3, Tns4, Tob2, Tomm20, Tomm22, Tomm34, Tomm40, Tomm6, Tomm7, Tomm70a, Top2b, Topaz1, Topors, Tor1a, Tor1aip1, Tor1aip2, Tor2a, Tor3a, Tor4a, Tox, Tox4, Tp53bp2, Tp53inp2, Tp53rk, Tp53tg3, Tp53tg3b, Tp53tg3c, Tp53tg3d, Tp63, Tp73, Tpbg, Tpcn2, Tpd52, Tpd52l3, Tpgs1, Tph1, Tpi1, Tpmt, Tpp1, Tpp2, Tpr, Tprkb, Tpst2, Tpt1, Tpte, Tra2a, Tra2b, Trabd2a, Traf1, Traf3ip1, Traf3ip2, Traf5, Traf6, Tram1, Tram1l1, Tram2, Trank1, Trappc13, Trappc2, Trappc2l, Trappc2p1, Trappc6b, Trappc9, Trat1, Trdmt1, Trdn, Trhde, Triap1, Trib1, Trim11, Trim14, Trim16, Trim16l, Trim2, Trim22, Trim24, Trim26, Trim27, Trim36, Trim38, Trim4, Trim40, Trim43, Trim45, Trim49, Trim49c, Trim49d1, Trim5, Trim52, Trim58, Trim59, Trim6, Trim64b, Trim65, Trim68, Trim71, Trim72, Trim8, Trio, Trip11, Trip12, Trip13, Trip4, Triqk, Trmt10a, Trmt10c, Trmt11, Trmt112, Trmt2b, Trmt5, Trmt6, Trnp1, Troap, Trpc1, Trpc6, Trpm5, Trpm7, Trpm8, Trpv1, Trpv2, Trpv3, Trpv5, Trub1, Trub2, Tsc22d2, Tsc22d3, Tsen15, Tsen34, Tsga10, Tshz1, Tsn, Tsnax, Tspan11, Tspan13, Tspan14, Tspan15, Tspan2, Tspan31, Tspan5, Tspan9, Tspo, Tspyl1, Tspyl4, Tspyl6, Tsr2, Tsr3, Tstd2, Ttc14, Ttc19, Ttc21b, Ttc22, Ttc26, Ttc28, Ttc30a, Ttc30b, Ttc33, Ttc37, Ttc39a, Ttc39b, Ttc39c, Ttc5, Ttc6, Ttc7a, Ttc7b, Ttc9c, Ttf2, Ttl, Ttll1, Ttll11, Ttll4, Ttll5, Ttll7, Ttll9, Ttn, Ttpa, Ttpal, Ttr, Tubal3, Tubb, Tubb2b, Tubd1, Tubgcp2, Tubgcp3, Tulp3, Tulp4, Tvp23a, Tvp23c, Twf1, Twistnb, Twsg1, Txn2, Txndc15, Txndc16, Txndc17, Txndc9, Txnip, Txnl1, Txnl4a, Txnl4b, Txnrd3, Tyro3, Tyrp1, Tyw5, U2surp, Uaca, Uap1l1, Uba2, Uba5, Uba52, Ubac1, Ubap2, Ubash3b, Ubb, Ubc, Ube2b, Ube2d1, Ube2e2, Ube2f, Ube2g1, Ube2g2, Ube2i, Ube2j1,*

| |
|---|
| *Ube2j2, Ube2l3, Ube2o, Ube2q1, Ube2t, Ube2z, Ube3a, Ube3b, Ube4a, Ubıad1, Ubl4b, Ubl7, Ublcp1, Ubox5, Ubp1, Ubqln1, Ubqln2, Ubqln3, Ubr2, Ubr4, Ubr7, Ubtd2, Ubtf, Ubxn10, Ubxn2a, Ubxn4, Ubxn7, Uchl3, Uchl5, Uck2, Uevld, Ufm1, Ugcg, Ugdh, Uggt1, Ugt2a3, Ugt2b10, Ugt2b4, Ugt8, Uhrf1bp1, Ulbp3, Umps, Unc13c, Unc45b, Unc5c, Unc5d, Unc79, Unc80, Uncx, Upf2, Upf3a, Upf3b, Upk3bl, Uprt, Uqcr11, Uqcrb, Uqcrc2, Uqcrfs1, Uqcrq, Urb1, Urb2, Urgcp, Ush2a, Usp1, Usp10, Usp15, Usp19, Usp22, Usp25, Usp26, Usp27x, Usp28, Usp3, Usp31, Usp36, Usp38, Usp4, Usp42, Usp43, Usp45, Usp46, Usp48, Usp51, Usp53, Usp6nl, Usp7, Usp9x, Usp9y, Utp11l, Utp14c, Utp15, Utp20, Utp23, Utp6, Utrn, Uvssa, Uxs1, Vamp1, Vamp4, Vangl2, Vash2, Vasp, Vbp1, Vcam1, Vcan, Vcpkmt, Vdac1, Vegfc, Ventx, Veph1, Vezt, Vgll1, Vhl, Vhll, Vıl1, Vım, Vıpr2, Vldlr, Vma21, Vmac, Vnn1, Vnn2, Vopp1, Vps13a, Vps13c, Vps18, Vps26a, Vps26b, Vps29, Vps33a, Vps33b, Vps36, Vps37d, Vps41, Vps45, Vps4a, Vps52, Vps8, Vsıg1, Vsıg4, Vstm2a, Vta1, Vtı1b, Vwa2, Vwa5a, Vwa7, Vwc2, Vwc2l, Wapal, Wasf1, Wasf3, Wasl, Wbp5, Wdfy1, Wdhd1, Wdr11, Wdr13, Wdr3, Wdr31, Wdr33, Wdr36, Wdr37, Wdr4, Wdr43, Wdr46, Wdr47, Wdr5, Wdr55, Wdr59, Wdr7, Wdr70, Wdr72, Wdr73, Wdr75, Wdr76, Wdr77, Wdr87, Wdr91, Wdr92, Wfdc1, Wfdc13, Wfdc8, Wfikkn2, Whamm, Whsc1, Whsc1l1, Wıpf1, Wıpf2, Wıpı1, Wıpı2, Wısp1, Wısp2, Wız, Wnt11, Wnt16, Wnt4, Wnt5a, Wnt8b, Wnt9b, Wrb, Wrn, Wsb1, Wsb2, Wscd1, Wtıp, Wwc1, Wwc3, Wwp2, Wwtr1, Xcl1, Xcr1, Xıap, Xırp2, Xk, Xkr8, Xkr9, Xpnpep3, Xpo1, Xpo4, Xpo5, Xpot, Xpr1, Xrcc2, Xrcc3, Xrcc4, Xrcc5, Xrcc6, Xxylt1, Yae1d1, Yap1, Yars2, Yeats2, Yeats4, Yıpf1, Yıpf2, Ylpm1, Yme1l1, Yod1, Ypel1, Ypel2, Ypel4, Ypel5, Ythdc2, Ythdf1, Ythdf3, Ywhab, Ywhag, Ywhah, Ywhaz, Zadh2, Zar1l, Zbed1, Zbed3, Zbed4, Zbtb16, Zbtb21, Zbtb25, Zbtb26, Zbtb3, Zbtb33, Zbtb34, Zbtb37, Zbtb41, Zbtb44, Zbtb5, Zbtb6, Zbtb8a, Zbtb8os, Zc2hc1a, Zc3h12b, Zc3h12c, Zc3h13, Zc3h14, Zc3h15, Zc3h18, Zc3h6, Zc3h8, Zc3hav1l, Zc3hc1, Zcchc11, Zcchc14, Zcchc2, Zcchc24, Zcchc3, Zcchc4, Zcchc5, Zcchc6, Zcchc7, Zcchc8, Zcchc9, Zcwpw2, Zdbf2, Zdhhc14, Zdhhc15, Zdhhc20, Zdhhc21, Zdhhc24, Zdhhc3, Zdhhc9, Zeb2, Zfand1, Zfand3, Zfand4, Zfc3h1, Zfhx3, Zfhx4, Zfp1, Zfp14, Zfp28, Zfp3, Zfp30, Zfp36, Zfp36l1, Zfp36l2, Zfp37, Zfp62, Zfp64, Zfp69b, Zfp82, Zfpm1, Zfr, Zfx, Zfy, Zfyve16, Zfyve20, Zhx1, Zhx2, Zıc1, Zık1, Zım2, Zım3, Zkscan1, Zkscan2, Zkscan3, Zkscan4, Zkscan8, Zmat1, Zmız1, Zmpste24, Zmym1, Zmym2, Zmym3, Zmym6, Zmynd19, Zmynd8, Znf100, Znf101, Znf106, Znf107, Znf117, Znf124, Znf131, Znf132, Znf134, Znf135, Znf136, Znf14, Znf140, Znf141, Znf146, Znf148, Znf154, Znf157, Znf160, Znf169, Znf17, Znf174, Znf175, Znf177, Znf195, Znf197, Znf2, Znf20, Znf200, Znf207, Znf208, Znf211, Znf212, Znf22, Znf221, Znf222, Znf223, Znf225, Znf227, Znf23, Znf234, Znf236, Znf25, Znf250, Znf251, Znf253, Znf254, Znf256, Znf257, Znf259, Znf26, Znf260, Znf263, Znf264, Znf266, Znf267, Znf273, Znf275, Znf277, Znf28, Znf280b, Znf280d, Znf283, Znf284, Znf285, Znf286a, Znf286b, Znf292, Znf318, Znf320, Znf321p, Znf324b, Znf326, Znf331, Znf333, Znf334, Znf33a, Znf33b, Znf343, Znf345, Znf347, Znf35, Znf354b, Znf354c, Znf365, Znf37a, Znf383, Znf384, Znf385d, Znf391, Znf394, Znf398, Znf404, Znf417, Znf418, Znf419, Znf426, Znf429, Znf43, Znf430, Znf431, Znf436, Znf440, Znf442, Znf446, Znf449, Znf45, Znf451, Znf460, Znf461, Znf462, Znf468, Znf470, Znf471, Znf473, Znf484, Znf486, Znf487, Znf490, Znf491, Znf492, Znf493, Znf497, Znf500, Znf503, Znf506, Znf512, Znf514, Znf517, Znf519, Znf525, Znf527, Znf528, Znf529, Znf532, Znf541, Znf543, Znf546, Znf548, Znf549, Znf550, Znf551, Znf552, Znf554, Znf555, Znf556, Znf557, Znf558, Znf559, Znf560, Znf561, Znf562, Znf566, Znf568, Znf57, Znf570, Znf572, Znf573, Znf578, Znf581, Znf582, Znf583, Znf584, Znf585a, Znf585b, Znf587, Znf587b, Znf589, Znf592, Znf594, Znf595, Znf597, Znf600, Znf605, Znf606, Znf607, Znf608, Znf610, Znf611, Znf613, Znf614, Znf616, Znf619, Znf620, Znf621, Znf622, Znf623, Znf624, Znf626, Znf629, Znf638, Znf639, Znf641, Znf652, Znf655, Znf66, Znf660, Znf664, Znf665, Znf667, Znf669, Znf670, Znf674, Znf675, Znf677, Znf678, Znf680, Znf681, Znf688, Znf689, Znf69, Znf695, Znf697, Znf699, Znf701, Znf703, Znf705a, Znf705g, Znf706, Znf708, Znf709, Znf713, Znf714, Znf716, Znf720, Znf721, Znf724p, Znf726, Znf730, Znf736, Znf737, Znf738, Znf74, Znf740, Znf746, Znf747, Znf749, Znf75a, Znf75d, Znf766, Znf770, Znf773, Znf776, Znf778, Znf780a, Znf780b, Znf781, Znf783, Znf786, Znf788, Znf790, Znf791, Znf793, Znf799, Znf8, Znf805, Znf808, Znf81, Znf813, Znf816, Znf829, Znf83, Znf830, Znf835, Znf839, Znf84, Znf841, Znf844, Znf845, Znf85, Znf850, Znf852, Znf860, Znf878, Znf879, Znf880, Znf891, Znf90, Znf91, Znf92, Znf93, Znf99, Znfx1, Znhıt6, Znrf1, Zranb1, Zranb3, Zrsr1, Zscan1, Zscan12, Zscan16, Zscan2, Zscan22, Zscan23, Zscan29, Zscan31, Zswım1, Zswım2, Zwınt, Zxda, Zxdb, Zxdc, Zyg11a, Zyg11b* and *Zzz3*. |

| **Target genes of miR-150-5p (n=72)** |
|---|
| *Ac027763.2, Aco1, Actr1b, Acvr1b, B3gnt1, Bcl9l, C17orf82, Camk2g, Cdan1, Cdk5r1, Cep85, Cers3, Cmtm6, Cnıh1, Cnppd1, Col4a4, Csf3, Dusp13, Dynlt1, Elovl1, Fam47e, Fdxr, Fndc5, Gabrg2, Gcm2, Gdı1, Ggnbp2, Grıpap1, Hılpda, Hmga2, Hnrnph3, Ikbıp, Itıh6, Kcnıp1, Lmo4, Mdga1, Metap1d, Mrvı1, Ndor1, Nefm, Notch3, Nup98, Nxph3, Osbpl9, Pa2g4, Pan2, Pdıa6, Plp2, Pom121c, Ppp2cb, Prr3, Prrt2, Rnf121, Rnf32, Sbf1, Scp2, Sec31b, Sh3bp5l, Shısa4, Slc16a2, Sparc, Steap3, Stx5, Tcte1, Tek, Tlr5, Tp53, Ube2e3, Upf1, Zcchc17, Znf322* and *Znf865* |

| Target genes of miR-126-3p (n=8) |
|---|
| *Plxnb2, Rgs3, Egfl7, Bak1, Pex5, Traf7, Ptpn9* and *Samd14.* |

| Target genes of miR-499a-5p (n=53) |
|---|
| *Abat, Adc, Adprh, Ahcyl1, Anks1b, Ap3s1, Aurka, C2cd2, Camta2, Cdca7, Cdkn1a, Chsy3, Cntn5, Colq, Comt, Dcaf12, Dnajb2, Dnm1l, Dusp11, Edar, Elavl2, Fam168b, Fubp3, Gata4, Glyr1, H2afz, Hnrnpa2b1, Hnrnpc, Hoxd10, Kıf3c, Lıms1, Mrps35, Msh6, Myh10, Nup88, Osbpl1a, P4ha1, Pcyox1, Polb, Pqlc3, Rab5c, Ralyl, Ranbp9, Rbm12, Rnf220, Scn4a, Scrn1, Shısa6, Slc45a4, Spns2, Srgap2, Srsf4* and *Tspan3.* |

**Human Plasma miRNA Assay Set 1**

| | 1 | 2 | 3 | 4 | 5 | 6 | 7 | |
|---|---|---|---|---|---|---|---|---|
| A | miR-495-3p | miR-3141 | miR-532-5p | miR-1231 | miR-223-5p | miR-4291 | miR-3651 | |
| B | miR-26b-3p | miR-146a-5p | miR-21-3p | miR-338-3p | miR-3200-5p | miR-769-5p | miR-4651 | |
| C | miR-624-5p | miR-139-3p | miR-1539 | miR-130a-3p | miR-339-5p | miR-132-3p | miR-22-5p | |
| D | miR-151a-3p | miR-877-5p | miR-100-5p | miR-18a-3p | miR-134-5p | miR-181c-5p | miR-1537-3p | |
| E | miR-98-5p | miR-20a-5p | miR-3191-3p | miR-766-3p | miR-140-5p | miR-328-3p | miR-133a-3p | |
| F | miR-331-3p | miR-487b-3p | miR-3176 | miR-181a-5p | miR-378e | miR-671-3p | miR-450a-5p | |
| G | miR-181d-5p | miR-330-3p | miR-4289 | miR-744-5p | miR-16-5p | miR-34c-3p | miR-106b-5p | [illegible] |
| H | miR-193b-3p | miR-576-5p | miR-10b-5p | miR-1277-3p | miR-188-5p | miR-30a-3p | miR-93-5p | |

**Human Plasma miRNA Assay Set 2**

| | 1 | 2 | 3 | 4 | 5 | 6 | 7 | |
|---|---|---|---|---|---|---|---|---|
| A | let-7c-5p | miR-184 | miR-152-3p | miR-3911 | miR-1909-5p | miR-205-5p | miR-1587 | |
| B | miR-16-2-3p | miR-3200-3p | let-7g-5p | miR-2110 | miR-196b-5p | miR-29c-3p | miR-127-3p | [illegible] |
| C | miR-4689 | miR-4516 | miR-377-3p | miR-4323 | miR-501-5p | miR-29a-3p | miR-596 | [illegible] |
| D | miR-196a-5p | miR-148b-3p | miR-4422 | miR-15a-5p | miR-421 | miR-337-5p | miR-7-5p | |
| E | miR-144-3p | miR-195-5p | miR-4454 | miR-99b-5p | let-7d-3p | miR-652-3p | miR-379-5p | [illegible] |
| F | miR-30c-5p | miR-32-5p | miR-192-5p | miR-1287-5p | miR-320b | miR-374a-5p | miR-3613-3p | [illegible] |
| G | miR-99a-5p | miR-335-5p | miR-146b-5p | miR-2355-5p | miR-490-3p | miR-136-5p | miR-196b-3p | |
| H | miR-425-3p | miR-590-3p | miR-96-5p | miR-486-5p | miR-615-5p | miR-378i | miR-154-5p | |

**Human Plasma miRNA Assay Set 3**

| | 1 | 2 | 3 | 4 | 5 | 6 | 7 | |
|---|---|---|---|---|---|---|---|---|
| A | miR-3610 | miR-1286 | miR-18b-5p | miR-34a-5p | miR-660-5p | miR-151b | miR-4687-5p | [illegible] |
| B | miR-373-3p | miR-1 | miR-422a | miR-631 | miR-627-5p | miR-451a | miR-194-5p | |
| C | miR-484 | miR-3653 | miR-224-5p | miR-4732-3p | miR-125b-5p | let-7i-5p | miR-3159 | [illegible] |
| D | miR-485-5p | miR-532-3p | miR-199a-3p | miR-4301 | miR-412-3p | miR-136-3p | miR-1301-3p | |
| E | miR-122-5p | miR-542-5p | miR-140-3p | miR-24-3p | miR-215-5p | miR-130b-3p | let-7a-5p | [illegible] |
| F | miR-574-3p | miR-191-3p | miR-7-1-3p | miR-675-3p | miR-23b-5p | let-7f-5p | miR-301b | |
| G | miR-223-3p | miR-25-3p | miR-19a-3p | miR-151a-5p | miR-625-3p | miR-30a-5p | miR-4538 | |
| H | miR-18a-5p | miR-487a-3p | miR-1233-3p | miR-637 | miR-942-5p | miR-197-3p | cel-miR-39-3p | |

**Human Plasma miRNA Assay Set 4**

| | 1 | 2 | 3 | 4 | 5 | 6 | 7 | |
|---|---|---|---|---|---|---|---|---|
| A | miR-199b-5p | miR-497-5p | miR-214-3p | miR-4732-5p | miR-3120-3p | miR-423-5p | miR-93-3p | |
| B | miR-3646 | miR-3907 | miR-338-5p | miR-143-3p | let-7i-3p | miR-29b-3p | miR-605-5p | [illegible] |
| C | miR-1281 | miR-423-3p | miR-877-3p | miR-22-3p | miR-203a | miR-454-3p | let-7f-1-3p | |
| D | miR-3183 | miR-542-3p | miR-337-3p | miR-381-3p | miR-505-3p | miR-363-3p | let-7g-3p | |
| E | miR-424-3p | miR-433-3p | miR-1976 | let-7d-5p | miR-548d-5p | miR-1183 | miR-4505 | |

| | | | | | | | |
|---|---|---|---|---|---|---|---|
| **F** | miR-4524a-3p | miR-17-5p | miR-720 | miR-128-3p | miR-181c-3p | miR-1225-3p | miR-26b-5p |
| **G** | miR-219a-1-3p | miR-210-3p | miR-17-3p | miR-320a | miR-15b-3p | miR-206 | miR-1237-3p |
| **H** | miR-142-5p | miR-145-5p | miR-369-3p | miR-202-3p | miR-4688 | miR-126-5p | cel-miR-39-3p |
